# Quantitative modelling of fate specification in the *C. elegans* postembryonic M lineage reveals a missing spatiotemporal signal

**DOI:** 10.1101/2024.09.01.610667

**Authors:** Benjamin Planterose Jiménez, Alexander R. Blackwell, João J. Ramalho, Sander van den Heuvel, Kirsten ten Tusscher, Erika Tsingos

## Abstract

The invariant lineages of *C. elegans* provide tractable cell fate models to study how developing organisms robustly integrate spatial signals at the single-cell level via gene regulatory networks. For instance, during postembryonic development, a mesoderm lineage arises through a sequence of oriented cell divisions from a single progenitor. This mesoblast initially gives rise to 18 cells with three distinct fates – 14 body wall muscles (BWMs), 2 coelomocytes (CCs; dorsal), and 2 sex myoblasts (SMs; ventral). The latter cells migrate and then proliferate to contribute 16 smooth muscles to the nematode’s reproductive organs. Prior work identified key symmetry breaking cues: i) ventrally restricted activation of the LIN-12 Notch pathway promoting SM over CC fate and ii) asymmetric re-distribution of SYS-1 *β*-catenin and POP-1 TCF among daughter cells along the anteroposterior (A-P) axis, i.e. the Wnt/*β*-catenin asymmetry pathway. However, it remains unclear whether these pathways are sufficient to specify all cell fates accordingly or whether additional symmetry breaking cues are necessary. In this study, we use quantitative modelling to better understand fate specification in the postembryonic M lineage. Specifically, we focus on the anteroposterior symmetry break by creating increasingly complex models towards robustly reproducing fate specification in wild type larvae and mutants. This iterative process resulted in two alternative models that explain the experimental observations by either introducing an additional spatial (spatial symmetry break) or temporal cue (temporal symmetry break). Finally, we evaluate their plausibility and propose a series of experiments to provide support for alternative models. Overall, our study highlights how a quantitative examination of mechanistic ideas can identify knowledge gaps and guide experimental follow-up.

## Introduction

In the development of multicellular organisms, cells integrate spatiotemporal signals via gene regulatory networks (GRNs). Ultimately, the output of GRNs modulate different cellular behaviors, including cell division, terminal differentiation, cell migration, apoptosis, cell fusion and quiescence. The *C. elegans* invariant cell-lineages have been widely studied for their remarkable reproducibility since they allow decision-making processes in the near absence of inter-individual variation (1). In particular, the postembryonic M lineage captures the interplay between proliferation and fate diversification in the context of already-formed larval structures (2, 3). Newly hatched L1 larvae contain a single mesodermal blast (M) cell. The M cell is an early descendant of MS (MS.apaapp) and is generated at the anterior left side of the embryo, and subsequently migrates to the posterior right side where it remains quiescent during embryogenesis (4). After hatching and triggered by the presence of food, possibly via the insulin-like growth factor signaling pathway in a cell non-autonomous fashion (5, 6), the M cell begins to divide midway through the L1 stage in a sequence of oriented cell divisions: First along the dorso-ventral (d/v), then left-right (l/r), and finally twice along anteroposterior (a/p) axes (1) (stages 1-M, 2-M, 4-M, 8-M and 16-M, respectively). Cells M.d(l/r)pa differentiate into large ovoid cells called coelomocytes (CCs), that are adjacent to the somatic musculature. CCs are capable of absorbing and concentrating soluble materials, although there is no evidence for active phagocytosis or migration (7). Cells M.v(l/r)pa undergo an additional cell division along the anteroposterior axis, giving rise to stage 18-M; the resulting cells M.v(l/r)paa become sex myoblasts (SMs) which later migrate towards the vulval region during the L2 stage. During L3, the SMs undergo three additional rounds of anteroposterior oriented cell divisions whose final products are 8 vulval muscle (4 VM1, 4 VM2) and 8 uterine muscle (4 UM1, 4 UM2) cells required for egg-laying (2, 8). All remaining M-derived cells acquire the cellular fate of body wall muscle cells (BWMs). Thus, by the end of L1 (i.e. 18-M), the M cell generated a total of 2 CCs, 2 SMs and 14 BWMs in hermaphrodites, making this lineage attractive to study the relationship between proliferation and cell fate specification.

Although fate specification and proliferation in the M lineage are invariant in the wild type, GRNs that give rise to such levels of precision remain incompletely defined. In principle, computational models can help determine to what extent the essence of a biological process has been revealed or whether critical steps are likely missing. Importantly, several key fate transcription factors have been previously identified which impact the M lineage cell fate, including *hlh-1* (BWM) (2), *ceh-34* and *eya-1* (CC) (9), and *sem-2* (SM) (10). Moreover, several signaling pathways are known to be involved in the fate specification of the postembryonic M lineage. For example, the Notch pathway is thought to provide a key dorso-ventral cue (11), whereby direct contact between cells expressing Notch ligand and cells expressing Notch receptor triggers the release of the Notch receptor’s intracellular domain (ICD) through several proteolytic steps. This ICD peptide translocates to the nucleus and associates with a CSL transcription factor and coactivator (LAG-1 and SEL-8, respectively in *C. elegans*) to trigger transcriptional activation (12). The *C. elegans* genome encodes two Notch receptors: *glp-1* and *lin-12*. Whilst *glp-1(0)* larvae have no alterations in the M lineage (11), mutations in *lin-12* disrupt M lineage dorsoventral asymmetries, with the ventral pattern changing to dorsal in *lin-12(0)* and reciprocal alterations (D→V) in *lin-12(gf)* mutants (11). These results indicate that the dorsal fate may be default and that LIN-12 Notch signaling is needed for ventral-specific M lineage fate induction. Consistent with such a role for LIN-12, the neighboring hypodermal cells on the ventral side upregulate Notch ligands APX-1 and LAG-2 at the spatiotemporal context of the M lineage (4-M according to translational fusion reporters) (11). In addition, a translational fusion reporter simultaneously labeling both the ICD and the full-length Notch receptor (LIN-12^ICD+FL^) can be detected from the 4-M stage onward (11). The ventrally-restricted activation of the LIN-12 Notch pathway has been further confirmed by reporters of LIN-12 proteolytic cleavage (13), LIN-12^ICD^ nuclear translocation (14) or a translational fusion reporter for the downstream target LAG-1 (15). Although the Notch pathway alone can explain the dorsoventral asymmetries of the M lineage, intriguing evidence indicates a role for the BMP pathway. BMP mutants display no ectopic M lineage fate transformations, however, in a *sma-9(0)* background, with D→V transformations similar to *lin-12(gf)*, compromising BMP signaling results in rescue of wild type fate specification. In fact, *lin-12(0); sma-9(0)* larvae experience a dorsoventral inversion of the M lineage (DV). SMA-9 is a homolog of Schnurri and was thought to act downstream of BMP signaling. To explain these observations, Foehr *et al* proposed that the BMP signaling can promote ventral fates on the dorsal side but is normally repressed by SMA-9 in the wild type (16).

The anteroposterior fate asymmetry in the M lineage is thought to be mediated by the Wnt/*β*-catenin asymmetry (W*β*A) pathway (9). This possibly nematode-specific variant of the canonical Wnt pathway participates in a large number of embryonic (e.g. EMS) and larval asymmetric cell divisions (e.g. T cell, seam cells, distal tip cells) (17–19). The subcellular localization and levels of Wnt/*β*-catenin pathway components guides cell fate decision-making. Ultimately, the nuclear levels of POP-1 TCF and SYS-1 *β*-catenin influence on whether their target genes are transcriptionally activated or repressed, and thereby the corresponding cellular fates. Indeed, nuclear POP-1 TCF and SYS-1 *β*-catenin show a characteristic asymmetric re-distribution following asymmetric M lineage divisions in the M lineage (9). Moreover, mutations in W*β*A pathway genes disrupt the normal anteroposterior asymmetry in M descendants (9).

Of note, both the regulation of the W*β*A pathway and the exact M daughter cell fate are highly complex. The *C. elegans* genome encodes four *β*-catenins: *bar-1, hmp-2 sys-1* and *wrm-1* (20). BAR-1 is the core *β*-catenin in the canonical Wnt pathway and can form transactivating heterodimers with POP-1. In contrast, HMP-2 does not interact with POP-1 but with cadherin HMR-1 and colocalizes at adherens junctions (though parallel signaling properties have been described in the context of the EMS cell (21)). WRM-1 and SYS-1 are both involved in the W*β*A pathway, though only the latter can form a trans-activating heterodimer with POP-1 (22). In the well-studied EMS cell, Wnt ligands secreted from a posterior source (i.e. P2) lead to the polarization of EMS by which the posterior pole becomes enriched in Wnt receptors (MOM-5 Frizzled) and Dishevelled proteins. As mitosis progresses, the membrane-tethered APR-1 APC preferentially localizes to the anterior pole, reaching its maximum asymmetry during telophase (23). APR-1, together with PRY-1 Axin, GSK-3 GSK3*β* and KIN-19 CK1*α*, form the destruction complex (hereon simply referred to as APC/Axin) (18). Upon cytokinesis, the anterior daughter cell inherits an excess of APC/Axin, which leads to down-stream asymmetries in SYS-1 and POP-1 via two distinct mechanisms: Firstly, APC/Axin promotes the phosphorylation of SYS-1, which triggers its ubiquitination and proteosomal degradation (asymmetric degradation). Secondly, APC/Axin promotes the nuclear export of WRM-1 *β*-catenin in a microtubule-dependent manner (asymmetric nuclear export) (23). WRM-1 is required for the phosphorylation of POP-1 by the Nemo-like kinase LIT-1, to trigger its export from the nucleus. This leads to the nuclear depletion of POP-1 specifically in the posterior daughter cell (where WRM-1 is enriched).

Importantly, POP-1 not only activates gene expression, together with SYS-1, but also acts as a transcriptional repressor. When nuclear POP-1 levels are high and SYS-1 levels low, as in anterior daughter cells, POP-1 forms repressor complexes, together with co-repressors UNC-37 Groucho and HDA-1 histone deacetylase (17). As a result, the asymmetric distribution of WβA pathway components leads to differential expression of POP-1 target genes in anterior versus posterior daughter cells. It is not fully understood how Wnt ligands create directionality in this process. In the M lineage, SYS-1 and POP-1 display posterior and anterior nuclear enrichment, respectively (9). Considering similarities with the EMS blastomere, a posterior Wnt source may be expected, but Wnt’s could also contribute instructive regulation instead of a directional cue (24). There are five Wnt genes in *C. elegans* (*cwn-1, cwn-2, egl-20, lin-44* and *mom-2*), which are expressed in a series of partially overlapping domains along the antero-posterior axis in L1 larvae. In addition, four Frizzled-like receptor genes (*mom-5, lin-17, mig-1* and *cfz-2*) are expressed as well as LIN-18 RYK and CAM-1 ROR Wnt receptors. Of potential relevance to the M lineage, EGL-20 Wnt forms an anteroposterior gradient originating from the tail during the L1 stage (25). Opposing gradients of EGL-20 and the inhibitor protein secreted Frizzled-related protein 1 (SFRP-1) have been implicated in the cell migration of Q neuroblasts with similar spatiotemporal context as the M lineage (26), suggesting a possible additional role in regulating the M lineage.

Adding another dimension of complexity, the prior picture of the W*β*A pathway focuses on the sequence of events surrounding a single cell division. However, M.d(l/r) and M.v(l/r) undergo two or three anteroposterior cell divisions, respectively. If and how SYS-1 and POP-1 asymmetries accumulate across several cell divisions has not been quantified in the M lineage. Zacharias *et al* have reported quantitative cumulative effects of this factors in early embryonic development, where cell divisions separated by as little as 15 minutes can occur (27). Nonetheless, it is uncertain whether the same picture applies to cell divisions in the M lineage, which occur approximately every 2 hours. Mutants in the W*β*A pathway display variable fate and proliferation alterations in the M lineage and are notably hard to interpret. For instance, loss of *sys-1* results in partial transformations affecting the most posterior part of the M lineage (M.(d/v)(l/r)pp →M.(d/v)(l/r)pa, with dorsal and ventral penetrance of 26 % and 84.2 %, respectively). However, the opposite trend is observed in *pop-1(RNAi)* larvae (M.(d/v)(l/r)pa →M.(d/v)(l/r)pp; dorsal and ventral penetrance of 48.4 % and 25 %). Strangely, the most anterior cells in the lineage (M.(d/v)(l/r)a(a/p)) very rarely experience any ectopic transformations in either mutants (28). Altogether, while the Notch and Wnt/*β*-catenin asymmetry pathways are clearly involved, it remains unclear whether and how these signaling pathways can instruct all different cell fates (BWM, CC, or SM) of the M lineage during L1 development.

In summary, although a wealth of experimental observations has identified the key pathways and genes, many details on how the different levels of regulation fit together remain elusive. In this study, we develop a series of quantitative mathematical models of increasing complexity to integrate multiple layers of evidence and to assess whether the known signaling pathways are sufficient to properly specify cell fates across the M lineage. Our modelling approach reveals that an additional spatial or temporal symmetry breaking cue is necessary to specify the anteroposterior fate determination. Overall, our study underscores how putting mechanistic ideas to the test via quantitative modelling can identify knowledge gaps that might otherwise go unnoticed, but also challenge established preconceptions and instruct future experimental investigations.

## Materials and Methods

All data analysis and modelling was performed in the R programming environment (29). Specifically, we used R-packages `deSolve` (30), `rootSolve` (31, 32), `PolynomF` (33) and `pracma` (34) to simulate systems of differential equations, to numerically compute steady-states and to solve polynomial equations with real coefficients. Data visualizations were achieved with `ggplot2` (35), `ggsci` (36), `reshape2` (37) and `scales` (38). We implemented symbolic calculations in the Python programming language (39) using the `SymPy` library (40). All scripts employed for the analysis of this manuscript have been made available at https://github.com/BenjaminPlanterose/MlineageAPsymmetrybreak.

## Results

### An apparent non-monotonic dose-response between Wnt pathway activity and cell fate decision

Towards building quantitative models of fate specification in the M lineage, we first sought to decompose the problem of fate specification by identifying axes of symmetry breaking. We noted that the fate specification in the M lineage is left-right symmetric (2). We can thus project the resulting fates at 18-M to a 2-dimensional grid (Fig 1A). The assumption of left-right symmetry also holds for mutants, for example, in mutants in key fate-promoting transcription factors such as *sem-2(0)* and *ceh-34(RNAi)* or *eya-1(0)*, either the SM or the CC transforms to BWM on both sides of the larvae (Fig 1B). Given this projection, the problem of fate specification can be decomposed into “a dorso-ventral symmetry break” (Why do fates in *row 1* differ from those in *row 2*?) and “an anteroposterior symmetry break” (Why do fates in *columns 3 or 3A* differ from those in *columns 1, 2, 3B and 4*?) (Fig 1A). We next sought to investigate the correspondence between published genetic data and our abstract symmetry break decomposition. Previous genetic screens identified the LIN-12 Notch pathway as a major regulator controlling the dorso-ventral symmetry break. Loss-or gain-in-function mutations in *lin-12* result in a dorso-ventral symmetric lineage (Fig 1B). By contrast, the contribution of BMP signaling is more complex. Foehr *et al* propose a model in which the dorsal M lineage may also receive a BMP signal that paradoxically promotes ventral fates, but is constitutively repressed by the presumed BMP antagonist SMA-9 Schnurri (11, 16). Several lines of evidence support this model: i) loss-of-function of the BMP signaling components pathway does not disrupt wild type M lineage fate specification (16); ii) loss of *sma-9* phenocopies *lin-12(gf)* (Fig 1B) (11); iii) compromising BMP in a *sma-9(0)* background results in a complete rescue of wild type fate specification (16); iv) *sma-9(0);lin-12(0)* larvae experience a dorso-ventral inversion of fate specification (11) (Fig 1B). The latter inversion can be explained by a simple superposition of *lin-12(0)* transforming the ventral to the dorsal fate (V→D), with *sma-9(0)* transforming the dorsal to the ventral fate (D→V). We further propose that the dorsally-restricted BMP signal may be functional in male M lineage fate specification, where SM cells are generated both dorsally and ventrally (1). Thus, SMA-9 may be required to induce sex-specific differences in the M lineage. In summary, BMP is most likely not required for hermaphrodite M lineage specification, while the LIN-12 Notch pathway is solely responsible for the ventral fate specification in hermaphrodites. Accompanying dorso-ventral signaling, the W*β*A pathway provides the only known anteroposterior cue in the M lineage. Complete loss-of-function of SYS-1 *β*-catenin mutant (*sys-1(0)*) and knock-down of POP-1 TCF (*pop-1(RNAi)*) larvae display variable ectopic transformations and proliferation patterns in the posterior *columns 3 and 4* (cells M.(d/v)(l/r)p(a/p)) whilst the anterior *columns 1 and 2* (cells M.(d/v)(l/r)a(a/p)) mostly retain their wild type identity (Fig 1B). All *columns* are produced by successive divisions along the anteroposterior axis. The first anteroposterior division generates the progenitors of the anterior *columns 1, 2* and the progenitors of the posterior *columns 3, 4*, while the second anteroposterior division generates the individual *columns 1, 2, 3, and 4*. The cell in *row 2, column 3* undergoes an additional round of anteroposterior division to generate the cells at *row 2, columns 3A and 3B*. The W*β*A pathway promotes asymmetries in nuclear content of SYS-1 and POP-1 in daughter cells upon asymmetric cell division in the anteroposterior axis (9). Thus, we expect that modulating this pathway should affect all *columns* to some extent. Instead, partially compromising the W*β*A pathway mainly affects the posterior side *columns 3 and 4*, which is unexpected. Since differences between *columns 1 and 2* and *columns 3 and 4* are generated during the first cell division, this raises the question to how the first anteroposterior cell division can be more robust to alterations in the W*β*A pathway than the latter ones. Thus, we focus on the time between 4-M and 18-M stage and ignore the variability in proliferation patterns seen in mutants since accounting for this would require a more complex modelling approach that integrates cell cycle entry and exit, and the interplay between proliferation and differentiation, which goes beyond the scope of this work.

**Fig. 1.**
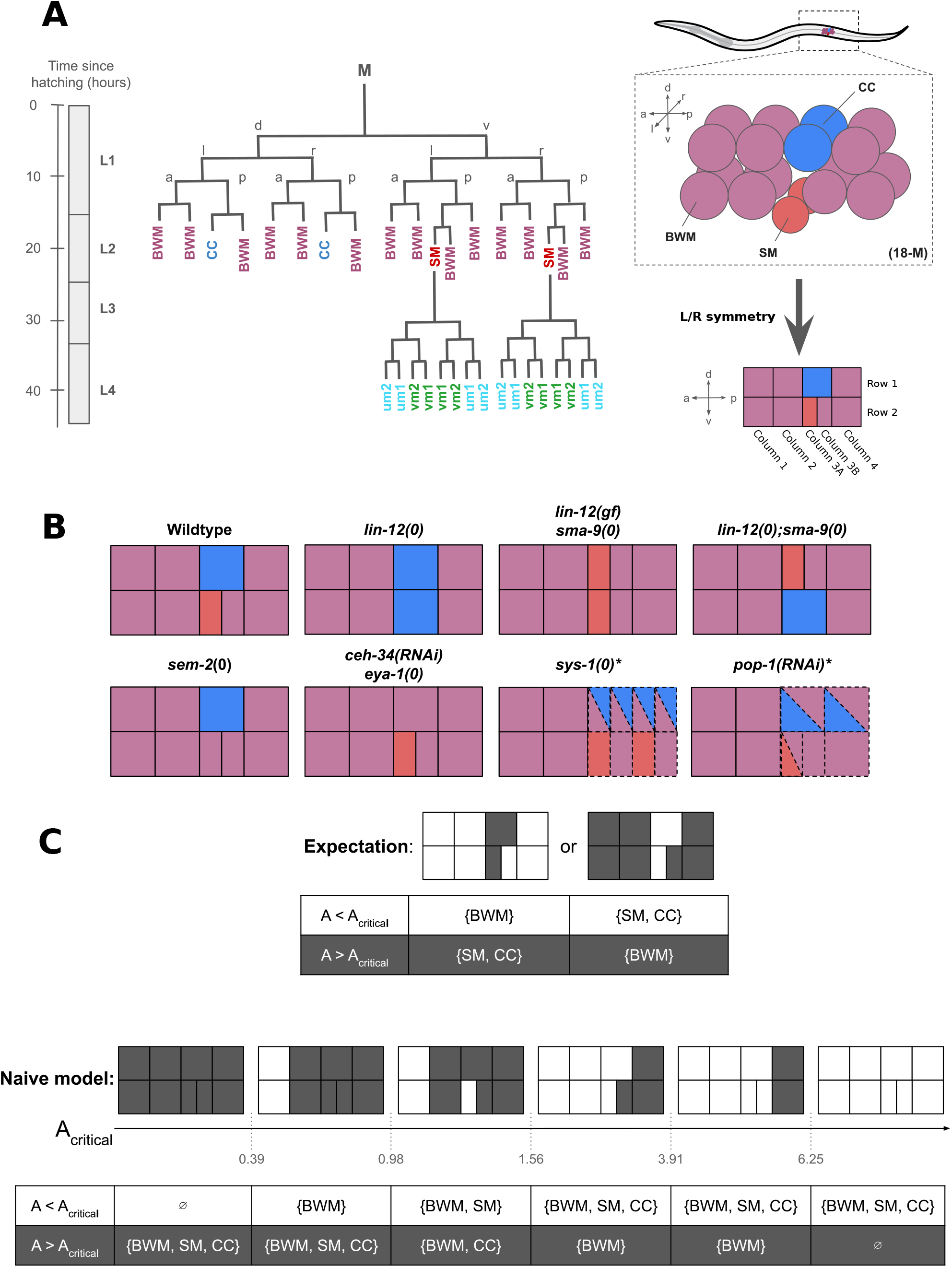
Wild type and mutant fate specification in the postembryonic M lineage and predicted anteroposterior symmetry breaks in the naive model. (A) Schematic of early M lineage in wild type hermaphrodites highlighting left-right symmetry in fate specification and the corresponding 2-D projection. (B) Fate specification in mutants projected into a 2-D grid, including *lin-12(0)* (11), *sma-9(0)* (16), *lin-12(gf)* (11), *lin-12(0);sma-9(0)* (16), *sem-2(0)* (10), *ceh-34(RNAi)* (9), *eya-1(0)* (9), *sys-1(0)* (9) and *pop-1(RNAi)* (9). Dashed lines symbolize variable cell divisions, whilst colored triangles represent variation in fate specification. (C) Naive model predictions across different values of *A*^Wnt^ and corresponding fates in the M lineage. anteroposterior symmetry break occurs when 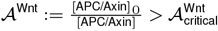. Values obtained assuming *ϕ*:= 0.8 (details in Suppl. Methods). M: M mesoblast, d: dorsal, v: ventral, l: left, r: right, a: anterior, p: posterior, BWM: body wall muscle; CC: coelomocyte; SM: sex myoblast; um: uterine muscle; vm: vulval muscle

To better understand how the successive anteroposterior divisions of *columns 1 through 4* may impact on the activity of the W*β*A pathway and anteroposterior symmetry break, we first consider a simple toy model, hereafter referred to as “the naive model”. We envision the M.(d/v)(l/r) cells at 4-M stage as having an initial number of APC/Axin molecules, which are redistributed among daughter cells by a proportion *ϕ* ∈ [0.5, 1], where *ϕ* = 1 corresponds to complete anterior redistribution, whilst *ϕ* = 0.5 corresponds to a symmetric cell division. For the sake of simplicity, we assume that APC/Axin is neither produced nor degraded, that the volume of the daughter cells and their nuclei shrink by a factor of two after every cell division (i.e. cleavage cell divisions with constant karyoplasmic ratio). We further assume that the activation of downstream Wnt targets is proportional to the ratio of nuclear SYS-1 and POP-1,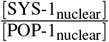, in line with the current understanding of the W*β*A pathway. This is equivalent to assuming that Wnt target activation is proportional to 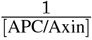, since asymmetric distribution of SYS-1 and POP-1 depends on APC/Axin. To estimate the asymmetric inheritance proportion *ϕ*, we turn to data from the ABpl lineage, which undergoes two successive asymmetric cell divisions under the influence of the W*β*A pathway. Here, the measured average 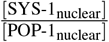 ratios were 0.2, 3.6, 1.1, 9.6 for aa, ap, pa and pp cellular products, respectively (27). We thus obtained a fit 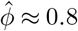 (*R*^2^ = 0.96, see Suppl. Method for details). We can express the relative activation of downstream Wnt targets as the ratio of the initial to the final APC/Axin content,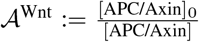. Our naive model predicts that this ratio follows a graded expression pattern decreasing along the anteroposterior axis, with the following hierarchy: 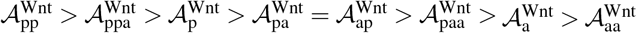. We assume that fate decision occurs at the 18-M stage after all divisions, and that there is a critical threshold of Wnt pathway activation, 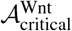, above which a cellular fate transition takes place. Under the strong assumptions of the naive model, the W*β*A pathway activity displays a monotonic dose-response and is unable to properly discriminate *columns 1, 2, 3B, 4* from *columns 3 and 3A* regardless of the chosen value of 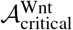 (Fig 1C). Furthermore, the naive model predicts that enrichment and depletion are commutative in the sense that the order of cell divisions does not affect the final result. In other words, a cell that is the posterior daughter in the first division and anterior in the second is equivalent to a cell that is the anterior daughter in the first and posterior in the second 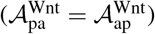. However, this contradicts the ABpl lineage data (27), since 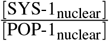 for ap (3.6) is significantly greater than for pa (1.1). In conclusion, the naive model predicts a graded monotonic Wnt pathway activity along the anteroposterior axis, which cannot explain why only *column 3* selects non-BWM fates in wild types nor can it explain the mutant phenotypes affecting only *columns 3 and 4*. Given all of the above, we decided to relax some of our assumptions, and construct a more faithful model of the W*β*A pathway.

### Detailed dynamical model of the Wnt/*β*-catenin asymmetry pathway is monostable

An unrealistic premise of the naive model was that APC/Axin is neither produced nor degraded, but solely re-distributed. In order to take into consideration the continuous production and degradation of APC/Axin and downstream components over time, we adopt the formalism of ordinary differential equations (ODEs). Here, we envision cellular fates as the stable fixed points of a multidimensional dynamical system (41). We compatibilize this formalism with cell divisions by introducing multiplicative discontinuities on the APC/Axin kinetics (i.e. hybrid system). In brief, we start from some given initial conditions, updating the system of ODEs until some prespecified time, *t*_division_, splitting the system into two copies, each representing a daughter cell. We then use the final condition of the mother cell as the initial conditions of the two daughter cells but having asymmetrically-distributed APC/Axin among daughter cells. As in the naive model, we still assume cleavage-like cell divisions with constant karyoplasmic ratio and a partition ratio *ϕ*.

To better understand how the W*β*A pathway enacts fate decision-making, we aimed to test to what extent the asymmetries in APC/Axin affect the downstream components in the pathway. At the same time, to make the model tractable, we restricted the number of components by assuming RNA quasi-steady state. To better represent the subcellular compartmentalization of the cell, we incorporated two compartments (cytosol and nucleus), and scaled kinetic equations for corresponding import/export shuttling terms for all components (42). We introduced ODEs for SYS-1, POP-1, SYS-1·POP-1 and their dimerization equilibrium dynamics (in both compartments), and LIT-1·WRM-1 complex (without explicitly including LIT-1 or WRM-1 as monomers, also to limit the number of components), all with constant production and first-order mass-action degradation (both in a cytosol-exclusive manner). We linked APC/Axin dynamics with the downstream components by introducing SYS-1 degradation promoted by APC/Axin in parallel to its basal degradation, cytosolic import of WRM-1·LIT-1 enabled by APC/Axin, and POP-1 nuclear export promoted by WRM-1·LIT-1. These additional processes were implemented as multiplicative terms based on the Hill equation (saturation curve, Hill coefficient of two to favor non-linear dynamics). Additionally, we assume that APC/Axin does not promote the degradation of cytosolic SYS-1·POP-1 and that SYS-1·POP-1 nuclear export is not favored by nuclear LIT-1·WRM-1. We hereon refer to this model as “Model W*β*A” (Fig 2A).

**Fig. 2.**
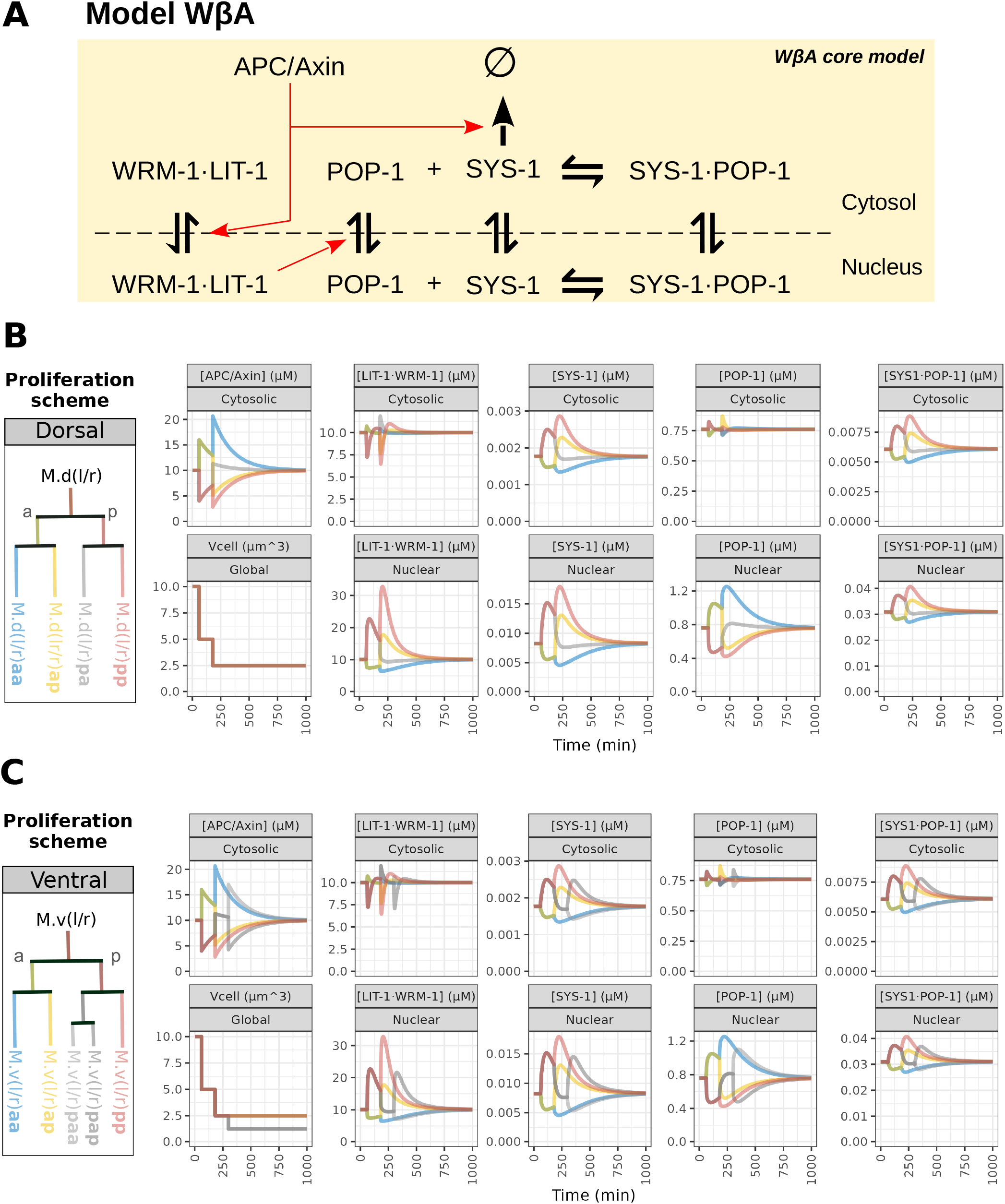
Model W*β*A in the cumulative regime. (A) Model diagram. Red arrows indicate regulatory interactions, bold black arrows indicate binding/unbinding. Arrows for production and degradation terms are omitted for clarity except for SYS-1 degradation. (B) Numerical simulations of time dynamics across W*β*A pathway components using dorsal and (C) ventral M lineage proliferation schemes.

Model W*β*A does not include any regulation of APC/Axin by the other components of the W*β*A pathway. The result of this is that the half-life of a perturbation on [APC/Axin] can be expressed solely as a function of the degradation rate *d*_APC/Axin_, given by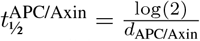, and thus, does not depend on the initial condition nor on the production rate *p*_APC/Axin_. A perturbed initial condition evolves towards the steady-state with rate 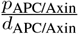. As a result, APC/Axin asymmetries generated in prior cell divisions tend to dampen over time. Thus, if cell divisions are spread too far apart in time, prior asymmetries are extinguished before they can interfere with present ones. In other words, cumulative effects in APC/Axin can occur only if Δ*t*_division_ is in the order of 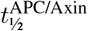. Under this regime, APC/Axin behaves in a non-commutative manner (enrichment-depletion is not equivalent to depletion-enrichment in APC/Axin). For 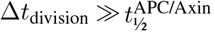, cell divisions are effectively independent of one another. Since it is unknown which option applies to the M lineage, we studied this model under both regimes: cumulative 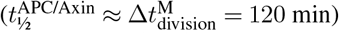 and a non-cumulative 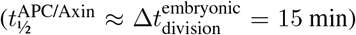. Though APC/Axin is set to evolve in time to a prespecified steady-state in Model W*β*A, APC/Axin asymmetries could potentially push multistable downstream components towards one steady-state or another. By analytically solving the steady-states of all the downstream components and with the help of symbolic calculus (40) (`Suppl. Methods`), we show that the entire system effectively evolves towards a single fix point in ℝ ^+^. Pathway-wide monostability has several implications.

Firstly, since each fate should correspond to a different stable fix point (in ℝ^+^), our modelling suggests that bistable gene regulatory motifs are likely located downstream of the W*β*A pathway rather than within it. Secondly, as for APC/Axin, we can construct a similar argument on the transient nature of the asymmetries of downstream W*β*A pathway components, with the caveat that their persistence in time may not adhere to 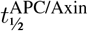. As these results are intrinsic to the model’s structure, they will generally be true regardless of the values chosen for model parameters.

In spite of the multiple simplifying assumptions, Model W*β*A still has a total of 26 free parameters which complicates its study especially given that no appropriate dataset exists to simultaneously fit all the model parameters. In fact, quantitative modelling is an excellent tool to explore parameter and solution space in the absence of such data (43). At this exploratory phase, we selected parameter values in the right order of magnitude range. For instance, we made sure that the time dynamics hierarchy was respected for different (sub)cellular processes: 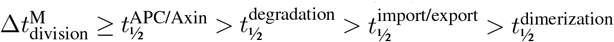. We chose kinetic parameters such that, in steady state, nuclear proteins are mostly nuclear, and that protein heterodimers are favored over the monomers (details in Suppl. Methods).

Furthermore, we were able to restrict the range of plausible values for production parameters for SYS-1 and POP-1 by considering experimental observations of fluorescent reporters (9), as explained in the following. Since translational fusion reporters of SYS-1 and POP-1 cannot distinguish their monomeric-from their complexed-form, what is observed under the confocal microscope are combined fractions. For a SYS-1::GFP construct, that means [SYS-1_nuc_] + [SYS-1·POP-1_nuc_], and for a POP-1::GFP construct that means [POP-1_nuc_] +[SYS-1·POP-1_nuc_]. Given that the SYS-1·POP-1 complex is the putative pathway activator, it is reasonable to expect that the asymmetry in its abundance across the M lineage anti-correlates with POP-1’s (the repressor). Consider the hypothetical case that [SYS-1·POP-1_nuc_] » [POP-1_nuc_], and that we are observing a POP-1::GFP construct. In this case, because the complex is more abundant than POP-1, and the GFP signal is the sum of [POP-1_nuc_] + [SYS-1·POP-1_nuc_], we would expect to see stronger signal in posterior cells. This is the opposite of what is observed.

In fact, M lineage cells display clear POP-1::GFP anterior enrichment (9). Thus, it is likely that the inverse is true: [SYS-1·POP-1_nuc_] « [POP-1_nuc_]. Going deeper into this point, nuclear SYS-1·POP-1 complex is in fact the point in the signaling pathway where two opposite asymmetries collide (i.e. SYS-1: p > a; and POP-1: a > p). Which asymmetry dominates over the other and how is destructive interference avoided? To complicate matters, the balance is also affected by cytosolic fractions and the different regulatory mechanisms acting on SYS-1 (i.e. degradation) and POP-1 (i.e. nuclear export). Leveraging the exploratory power of our model, we tested a range of productions rates for SYS-1 and POP-1 (**Fig S1-2**). We observe that excess of SYS-1, POP-1, or both abolished the observable asymmetries in [POP-1_nuc_] + [SYS-1·POP-1_nuc_], [SYS-1_nuc_] + [SYS-1·POP-1_nuc_], or both, respectively (**Fig S1-2**). Simply put, high production rates result in an excess of background levels of the SYS-1·POP-1 complex. This is because the SYS-1·POP-1 complex is not asymmetrically regulated (only its constituent monomers are). Thus, high background levels of the SYS-1·POP-1 complex can hide the effects of upstream dynamics in SYS-1 and POP-1. We used this information to select the reference values for the production rates of SYS-1 and POP-1 in our simulation (see Supp. Methods for more details). We also evaluated the influence of all parameters on the maximum transient asymmetries of nuclear POP-1 and SYS-1·POP-1 (**Fig S3-4**). For example, we find that the degradation rate of SYS-1 promoted by APC/Axin (*d*_APC/Axin:SYS-1_) contributed the most to the SYS-1·POP-1 complex asymmetries. This indicates that regulation of SYS-1 nuclear export by APC/Axin is a major control point of the pathway.

Having parameterized the model with the above considerations, we ran numerical simulations of the time dynamics across components for the cumulative (Fig 2B-C) and the non-cumulative regimes of Model W*β*A (**Fig S5**). All observed asymmetries are transient, as predicted by our symbolic calculations on steady-states.

Model Wnt*β*A reveals several interesting insights. Firstly, since nuclear import/export kinetics are proportional to nuclear surface area which scales as 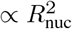 while the cellu-lar volume scales as 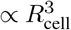, every cleavage cell division increases transport kinetic rate by a factor of 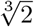, assuming constant karyoplasmic ratio. We used our derived analytical solution of the steady-states to observe the effects of *V*_cell_ on the steady-state concentrations of nuclear SYS-1, POP-1 and SYS-1·POP-1 (**Fig S6**). Interestingly, the influence of cellular volume is marginal. Thus, our model predicts that the volumetric effects after the third round of cell division of the SM mother (M.v(l/r)pa) in *row 2, column 3* are likely negligible. We also tested the influence of the production rates of SYS-1 and POP-1 (*p*_sys-1_, *p*_pop-1_ and *p*_wrm-1·lit-1_) on corresponding steady-states. Both *p*_sys-1_ and *p*_pop-1_ positively correlate with the steady state level of the complex [SYS-1·POP-1]_∞_. Further, *p*_sys-1_ correlates with [SYS-1]_∞_ and anti-correlates with [POP-1]_∞_, whilst the opposite occurs for *p*_pop-1_. In other words, lower *p*_sys-1_ (like in a *sys-1(0)* mutant) is associated to a larger pool of POP-1 since there is a lower copy number of SYS-1 to sequester POP-1 in its heterodimeric form.

### Adding bistable motifs downstream is insufficient to reproduce the non-monotonic dose-response

We previously showed that model W*β*A is inherently monostable and thus it is unable to break the anteroposterior symmetry and make decisions on cellular fates. To address this limitations, we can however append additional elements downstream of W*β*A in order to supplement the GRN with the property of bistability (41). As the most simple scenario, we consider including a hypothetical downstream target, activated by the nuclear SYS-1·POP-1 complex and repressed by nuclear POP-1. To impart bistability, we further propose that our hypothetical target can self-activate and thus, self-maintain even in the absence of the activating SYS-1·POP-1 complex. Here, to avoid explicitly modelling the multitude of regulatory subprocesses acting upon this hypothetical target (e.g. transcriptional activation and repression by SYS-1·POP-1 and POP-1; target RNA degradation, shuttling and translation; target protein shuttling; binding of target protein to target promoter; transcriptional self-activation by target protein), we grouped these subprocesses into a single equation as an approximation to ensure the tractability of the model. Altogether, we refer to this model as “Model W*β*A+B” (Fig 3A).

**Fig. 3.**
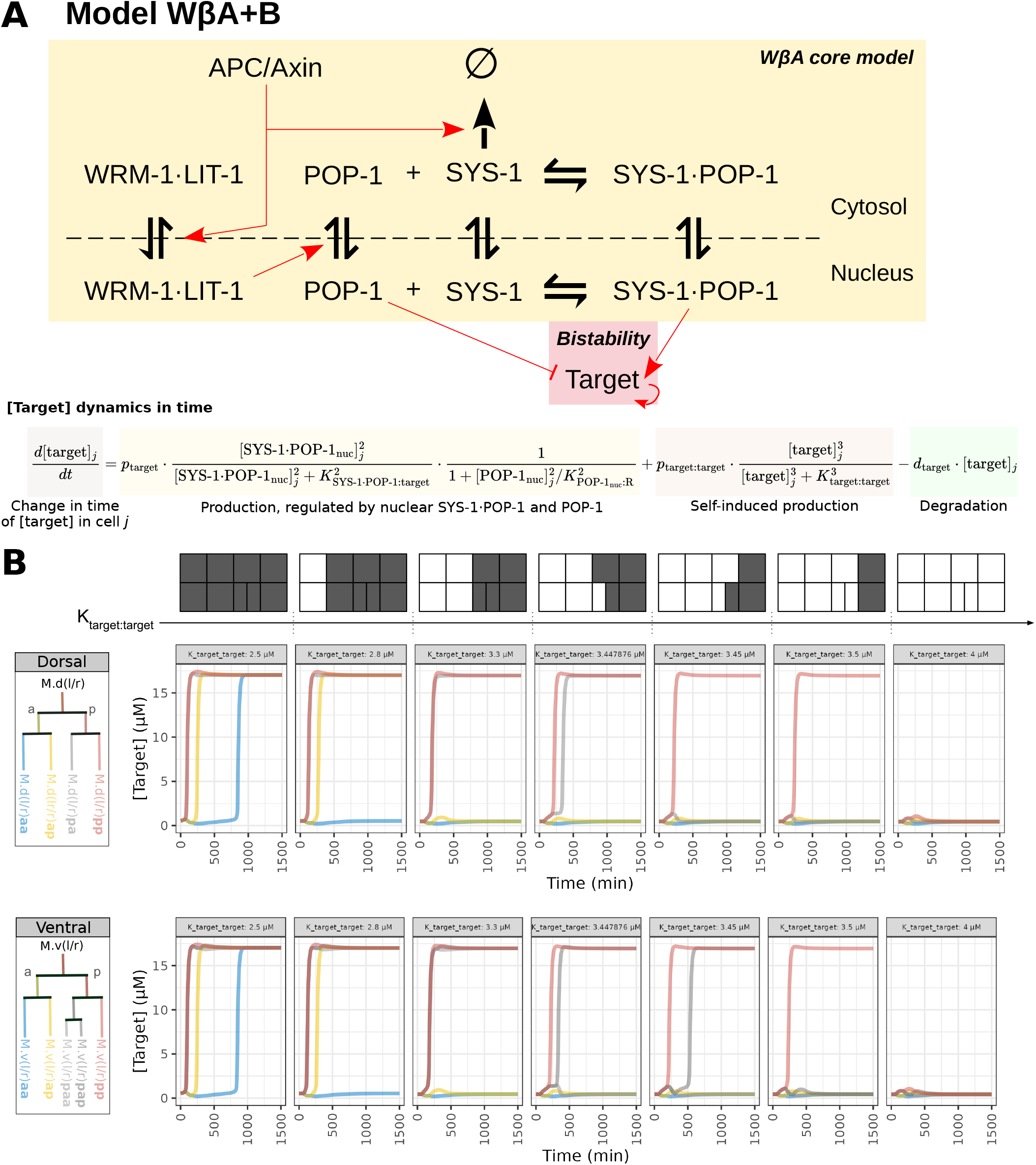
Simulation for a model of the Wnt/*β*-catenin asymmetry pathway with a bistable downstream target (W*β*A+B). (A) Model diagram. (B) Numerical simulations of time dynamics across model components using dorsal and ventral M lineage proliferation schemes for different values of *K*_target:target_ ∈ {2.5, 2.8, 3.3, 3.447876, 3.45, 3.5, 5} *µM*. Solely target concentrations are shown. Steady-state target concentrations are represented as 2-D projections on a grid.

We thus proceeded to characterize Model W*β*A+B. In the most trivial parametrisations, either all cells or none of the cells activate the target gene. With careful selection of the parameters, we can observe a transition sequence connecting both extreme behaviors (Figure 3B). This sequence bears high resemblance to that of the naive model (Figure 1). Similarly, this model has the same problem: None of the intermediates coincide with either *columns {3 and 3A}* or *columns {1, 2, 3B, 4}*. Thus, although this model can produce a symmetry break, it cannot fully explain the fate specification exclusively in *columns 3 and 3A*. To rephrase, Model W*β*A+B behaves “monotonically” in the selected parameter space neighborhood: A higher dose of W*β*A pathway activity leads to higher activation of the downstream target. However, to specifically activate cells M.d(l/r)pa and M.d(l/r)paa in *column 3*, there needs to be a change of directionality of the pathway’s effects. In other words, we expect a posterior activation in the first cell division followed by anterior activation in the second and third cell divisions. We refer to this property as a “non-monotonic dose-response”. We thus conclude that GRNs with this property are required to properly reproduce fate specification in the M lineage.

### The W*β*A pathway’s non-monotonic dose-response in the M lineage can originate from incoherence in the fate specification GRN

We explored several avenues to incorporate a non-monotonic dose-response into our model. Firstly, we considered to trivially replace the Hill-type saturation by a non-monotonic link function. This however does not provide any mechanistic insights into the underlying source of non-monotonicity of the system and thus we discarded this idea.

Secondly, we explored whether stoichiometric effects of SYS-1 and POP-1 abundance on the equilibrium of the SYS-1·POP-1 complex could serve as the source of non-monotonicity. We expected APC/Axin to cause significant shifts in the relative proportions of SYS-1 and POP-1. If SYS-1 is in excess, an increase in POP-1 can drive a substantial increase in the complex formation rate. In this scenario, if the downstream activation mediated by the SYS-1·POP-1 complex surpasses the repression by POP-1, an increase in POP-1 levels can indirectly lead to a higher activation of the downstream target, producing a non-monotonic dose-response. We previously searched for combinations of *p*_POP-1_ and *p*_SYS-1_ that maximize SYS-1·POP-1 asymmetries (**Fig S1-2**). In this search we noticed that for low parameter values of *p*_POP-1_ and *p*_SYS-1_, the [SYS-1·POP-1] complex can indeed behave non-monotonically. Thus, we decided to combine W*β*A in this carefully selected parameter regime with a downstream repressor that is solely regulated by the SYS-1·POP-1 complex and a bistable target gene (hereafter named “Model W*β*A+B+S”, where S stands for stoichiometry).

In Model W*β*A+B+S, small changes early on can have strong effects at later time points and hence, the model is highly sensitive to changes in parameters (**Fig S7**). Interestingly, we obtain a different fate sequence than the W*β*A+B model. Although the W*β*A+B+S model can lead to target activation specifically in *column 3*, it struggles to separate M.v(l/r)paa from M.v(l/r)pap. In fact, for our chosen parametrisation, the target is more easily activated in M.v(l/r)pap than M.d(l/r)pa, whereas the expected activation pattern should be reversed. Therefore, this model too cannot fully explain the fate specification in the M lineage. Moreover, this regime only appears at low SYS-1 and POP-1 copy numbers, which would be expected to be largely susceptible to biological noise. Altogether, it seems unlikely that such stoichiometric effects are the source of non-monotonicity in the anteroposterior specification of the M lineage.

As a final attempt to incorporate a non-monotonic dose-response into our model, we explored adding new network motifs downstream. We hypothesized that incoherent feedforward loops could result in a non-monotonic dose-response. Incoherence (e.g. mixed signals to the downstream target) makes low and high signals converge to the same fate whilst intermediate signals can give access to an alternative fate. Based on this consideration, we propose a new model which we term “Model W*β*A+B+I” (Fig 4A), where “I” stands for incoherence. We succeeded in finding a combination of parameters in which M.d(l/r)pa and M.v(l/r)paa activate the target but not the complementary set of cells (Fig 4B). The robustness of this solution with respect to variations in cell division timing is however quite limited (Fig 4B, right). This is critical, given the experimentally observed intrinsic variability in cell division timing of M lineage cells (standard deviation,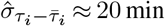; where assuming an underlying normal distribution, we expect ≈95 % of cell divisions to take place in the range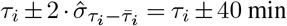; data not shown). Moreover, model W*β*A+B+I is also highly sensitive to minute changes in parameter space (**Fig S8**). Therefore, though the W*β*A+B+I model can reproduce the expected pattern of W*β*A pathway activation, we expect that the regulatory circuit should have additional features that ensure robustness.

**Fig. 4.**
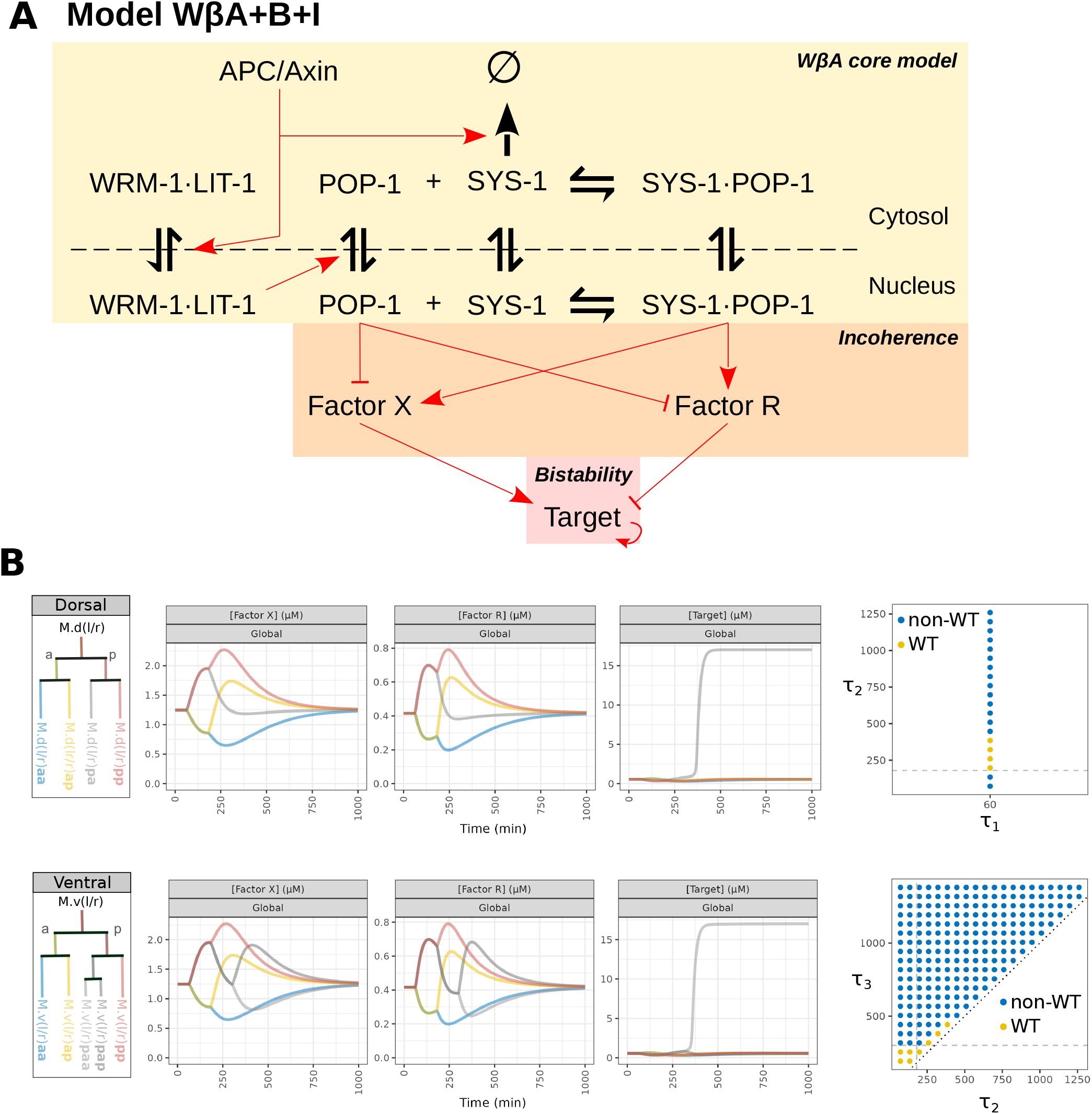
Simulation for a model of the Wnt/*β*-catenin asymmetry pathway with a bistable downstream target and incoherence (W*β*A+B+I). (A) Model diagram. (B) Numerical simulation of time dynamics using the dorsal or the ventral proliferation schemes and timing robustness, where *τ*_1_ = 60 min; *τ*_2_ = 180 min; *τ*_3_ = 300 min, correspond to the different times of anteroposterior cell divisions.

### Memory of the first cell division improves robustness to cell division timing variation

Towards identifying a functional GRN that can also explain the cell division timing robustness in fate specification, we decided to investigate the effects of additional interactions. As recapitulation, due to the monostability of the W*β*A core model, information regarding prior anteroposterior cell divisions tends to be erased in time. In the W*β*A+B+I model, this information is imprinted in the factor [X]. However, with some delay, [X] also eventually returns to a single steady-state and hence “forgets” its prior configurations. We thus sought to investigate whether the timing of the second and third anteroposterior cell divisions could be made more permissive if X was set up to be bistable. This way, X could promote a more robust decision making by establishing its expression in the first cell division and remaining either in the ON or OFF state in successive cell divisions. This consideration was inspired by the translational reporters of genes *let-381* and *unc-130* that turn-on in M.d(l/r)p and M.v(l/r)p, respectively, and remain in an ON state throughout their lineage (44, 45). Similarly to how we implemented bistability of the downstream target in model W*β*A+B, we introduced bistability of X by including a self-activation interaction. We refer to this model as “model W*β*A+B+I+M” (Fig 5A), where M stands for memory (of the first cell division). Introducing bistability of X has important consequences on the analytical solution of the target’s steady state levels, [target]_∞_: there are consequently two stable solutions for X (𝒳_off_, 𝒳_on_) and four stable solutions for the downstream target 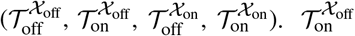 is considerably harder to reach than 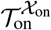; in other words, X in the ON state makes the target ON state more reachable. Since X is expected to switch on at the posterior side of the M lineage, we expect the following: M.(d/v)(l/r)a(a/p)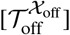; M.d(l/r)pa and M.d(l/r)paa 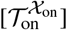; M.d(l/r)pp and M.d(l/r)pap 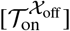. Hence, model W*β*A+B+I+M predicts a higher background concentration of the target in the most posterior side of the lineage. This is notable, because a higher target level implies a higher chance to switch to the target ON state in the presence of biological noise in the posterior *columns 3 and 4*. Despite these differences on the nature of the analytical solution of the GRN’s steady-states, we successfully parameterized model W*β*A+B+I+M to reproduce wildtype fate specification. Significantly, model W*β*A+B+I+M achieved this specification whilst also displaying greater time robustness than in the W*β*A + B + I model that lacked memory of the first cell division (Fig 5B-C, Fig S9).

**Fig. 5.**
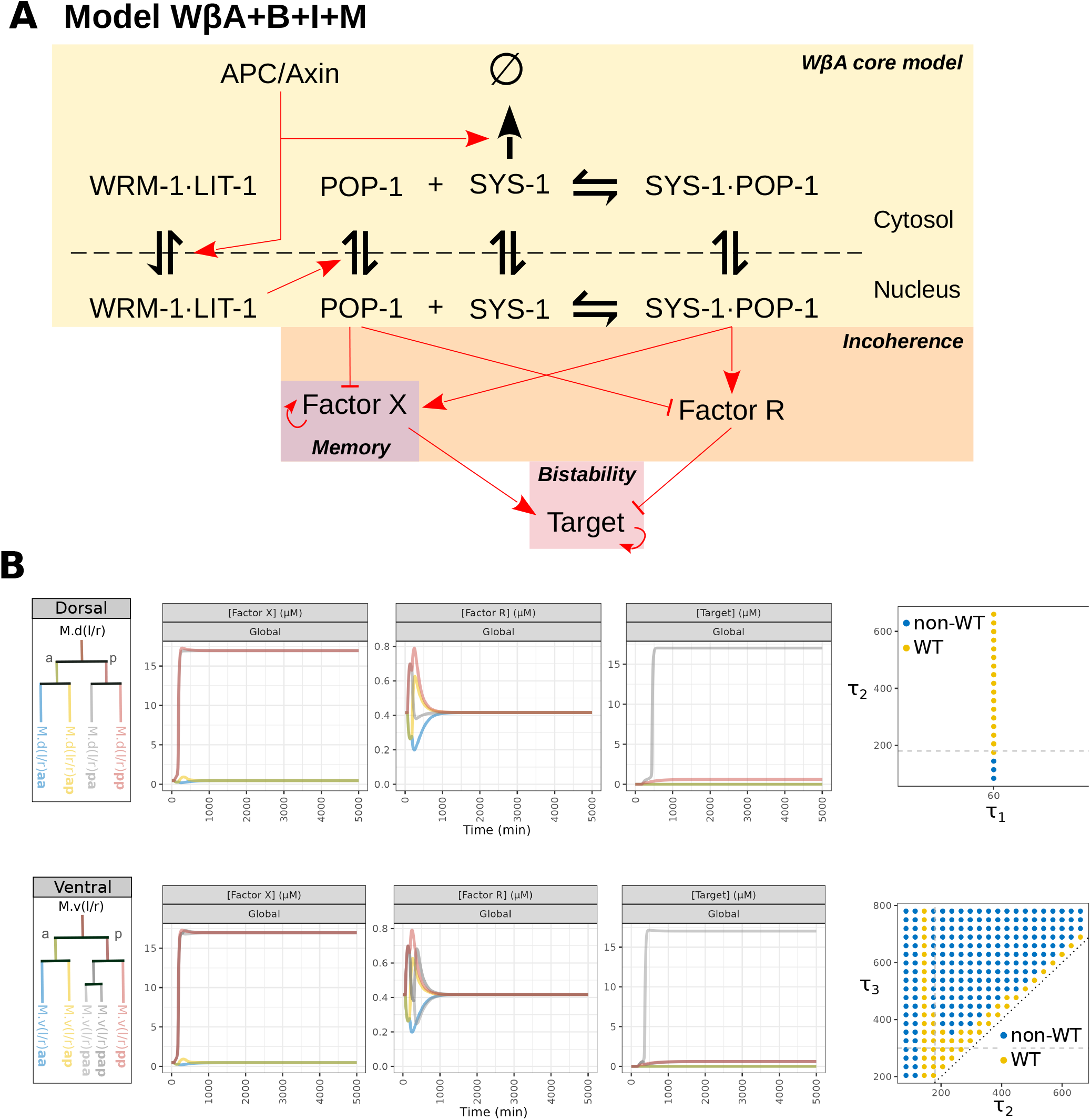
Simulation for a model of the Wnt/*β*-catenin asymmetry pathway with a bistable downstream target, incoherence and memory of the first cell division (W*β*A+B+I+M). (A) Model diagram. (B) Numerical simulation in the dorsal and ventral side proliferation schemes and timing robustness, where *τ*_1_ = 60 min; *τ*_2_ = 180 min; *τ*_3_ = 300 min, correspond to the different times of anteroposterior cell divisions.

Having explored the properties of the W*β*A+B+I+M model in wild type, we next explored whether it could also explain fate specification in mutants of the W*β*A pathway.

In a complete loss-of-function *sys-1(0)* mutant, we expect that [SYS-1·POP-1] = 0. Because the SYS-1·POP-1 complex is strictly required for initial factor X production in the W*β*A+B+I+M model, factor X would remain silenced across the M lineage. Therefore, the W*β*A+B+I+M predicts an anteroposterior symmetric M lineage where no cell activates the downstream target gene. This is however not what is observed in *sys-1(0)* larvae, which exhibit a clear anteroposterior asymmetry (Fig 1B). It is unlikely that other *β*-catenins compensate for SYS-1 in *sys-1(0)* larvae, since *wrm-1* and *hmp-2* have divergent functions, and *bar-1* is thought to be mainly involved in the canonical Wnt pathway (20). In summary, model W*β*A+B+I+M reproduces wild type fate specification whilst achieving greater cell division timing robustness than the W*β*A+B+I model that lacked memory of the first cell division. However, though the W*β*A+B+I+M model exhibits an asymmetry in target activation in the posterior *columns 3 and 4*, it cannot explain the fate specification in W*β*A pathway mutants.

### A model with an additional temporal cue can explain fate specification in wild type and mutants

A model’s ability to predict mutant phenotypes is fundamental to its utility. To this end, we implemented multiple changes to increase the compatibility of model W*β*A+B+I+M with known mutant phenotypes in the W*β*A pathway. Firstly, we removed the activation of SYS-1·POP-1 to X so that the expression of X is not abolished in the *sys-1(0)* mutant condition. Having removed this activation interaction and since we assumed throughout that the system begins in a steady-state and the default state of X is OFF, we now required another activator for X. As a solution, we were inspired by the expression pattern of the protein MLS-2 in the M lineage, which is activated early in the lineage but inactivated before cell fate decisions (46). We thus proposed a hypothetical factor P that is transiently activated during the first anteroposterior cell division, which we introduced in the model as a discontinuous pulse at *τ*_1_. In other words, P serves as a temporal cue of the system. Additionally, factor P ensures that factor X is activated on the lineage’s posterior side during the first anteroposterior cell division but not thereafter. Serendipitously, we also realized that model robustness was improved after removing the repression of R by POP-1. This interaction, a remnant of the prior incoherence model (W*β*A+B+I), contributes to the ectopic activation of the downstream target in *column 1* (cells M.(d/v)(l/r)aa). Thus, its removal lead to more robust target expression exclusively in *column 3*. We refer to this updated model as the “temporal symmetry break (TSB) model” (Fig 6A).

**Fig. 6.**
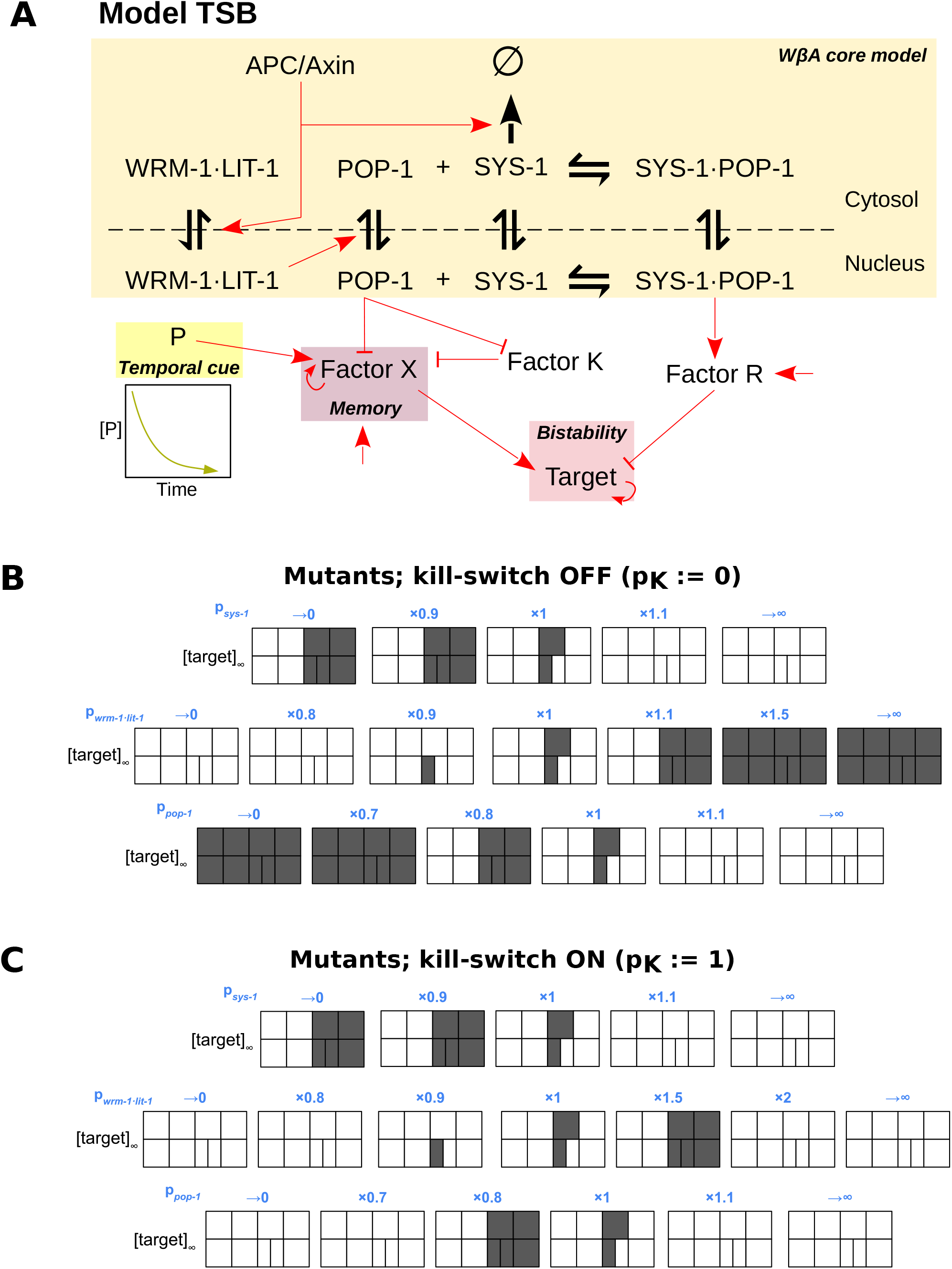
Numerical simulations of the Temporal Symmetry Break (TSB) model. (A) Model diagram. Steady-state concentrations of downstream target at different fold change levels of production rates of SYS-1, POP-1 and LIT-1·WRM-1, *p*_SYS-1_, *p*_POP-1_ and *p*_LIT-1·WRM-1_, relative to reference parametrisation (B) without kill-switch, (C) or with kill-switch. The kill-switch ensures that factor X can be only be activated if in the presence of POP-1.

We thus proceeded to evaluate its properties. The TSB model can explain fate specification in the wild type M lineage (**Fig S10**), with substantial robustness with respect to cell division timing and parameter values (**Fig S11**). In TSB, the entire fate specification process relies on the W*β*A pathway.

The TSB model also correctly recapitulates the fate specification defects in *sys-1(0)* larvae (Fig 6B, *p*_sys-1_ →0). Furthermore, the TSB model predicts that a hypothetical full POP-1 loss-of-function mutant *pop-1(0)*, which compromises both the activating and repressing functions of POP-1, should produce an anteroposterior symmetric M lineage. Theoretically, this symmetry could occur in two configurations: Either the downstream target is expressed in all cells (all-ON) or in none of them (all-OFF). Since POP-1 restricts the activation that P enacts upon X, an absence of POP-1 would cause the activation of X in both M.(d/v)(l/r)a and M.(d/v)(l/r)p. If background levels of R are not enough to repress the downstream target, this would result in an all-ON configuration. To assess whether the all-OFF *pop-1(0)* configuration was also theoretically possible, we introduced a modification on the network, here referred to as a “kill-switch”. The purpose of this kill-switch is to introduce a requirement for residual levels of POP-1 in order to activate X. In practice, we implemented this by introducing a protein K (standing for killswitch), repressed by POP-1 and which itself represses X, but which is normally repressed by POP-1 very effectively 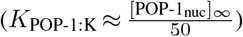. Given these settings, we can switch this interaction on (*p*_*K*_ = 1) or off (*p*_*K*_ = 0) without altering fate specification in the wild type, nor in the experimentally observed mutants for SYS-1 (*sys-1(0)*, equivalent to Fig 6B-C, *p*_sys-1_ →0) and the knockdown of POP-1 (*pop-1(RNAi)*, equivalent to Fig 6B-C, *p*_pop-1_ ×0.8). In contrast, in the total absence of POP-1, K is upregulated and therefore blocks any potential activation of X, resulting in an all-OFF configuration. Different sequence of fates turning on and off can be achieved by tuning *p*_SYS-1_, *p*_WRM-1·LIT-1_ and *p*_POP-1_ with and without the inclusion of K (Fig 6B-C). Therefore, the TSB model is compatible with wild type and mutant fate specification and it predicts an anteroposterior symmetric state in the complete absence of POP-1, which has yet to be experimentally verified.

### A model with an additional spatial cue is also consistent with wild type and mutant fate specification

As an alternative to an additional temporal cue, it is also possible that the W*β*A pathway is not the only anteroposterior cue involved in the fate specification of the M lineage. We aimed to account for this scenario to compare its predictions with those of the TSB model. We envision a posteriorly-enriched morphogen-like substance, M, that diffuses in a gradient along the anteroposterior axis. We modeled this process as 1-dimensional diffusion with a step function initial condition. We consider both a permanent gradient (*D* = 0) and a transient gradient (*D >* 0). The morphogen M drives the expression of a factor X, with an equivalent role to that of the TSB model. Having externalized the symmetry break between M.(d/v)(l/r)a and M.(d/v)(l/r)p, the W*β*A pathway function in this model is to select M.d(l/r)pa and M.d(l/r)paa over M.d(l/r)pp and M.d(l/r)pap via a downstream repressor, R. We refer to this model as the spatial symmetry break (SSB) model (Fig 7). The SSB model explains fate specification in the wild type M lineage (**Fig S12**), and is comparable to the TSB model in terms of robustness (**Fig S13**). Thus, from a theoretical modelling perspective, the SSB is an equally plausible alternative model for fate specification in the M lineage. Notably, we predict that the fate map of a complete null mutant of POP-1 would be able to discriminate between the TSB and SSB models.

**Fig. 7.**
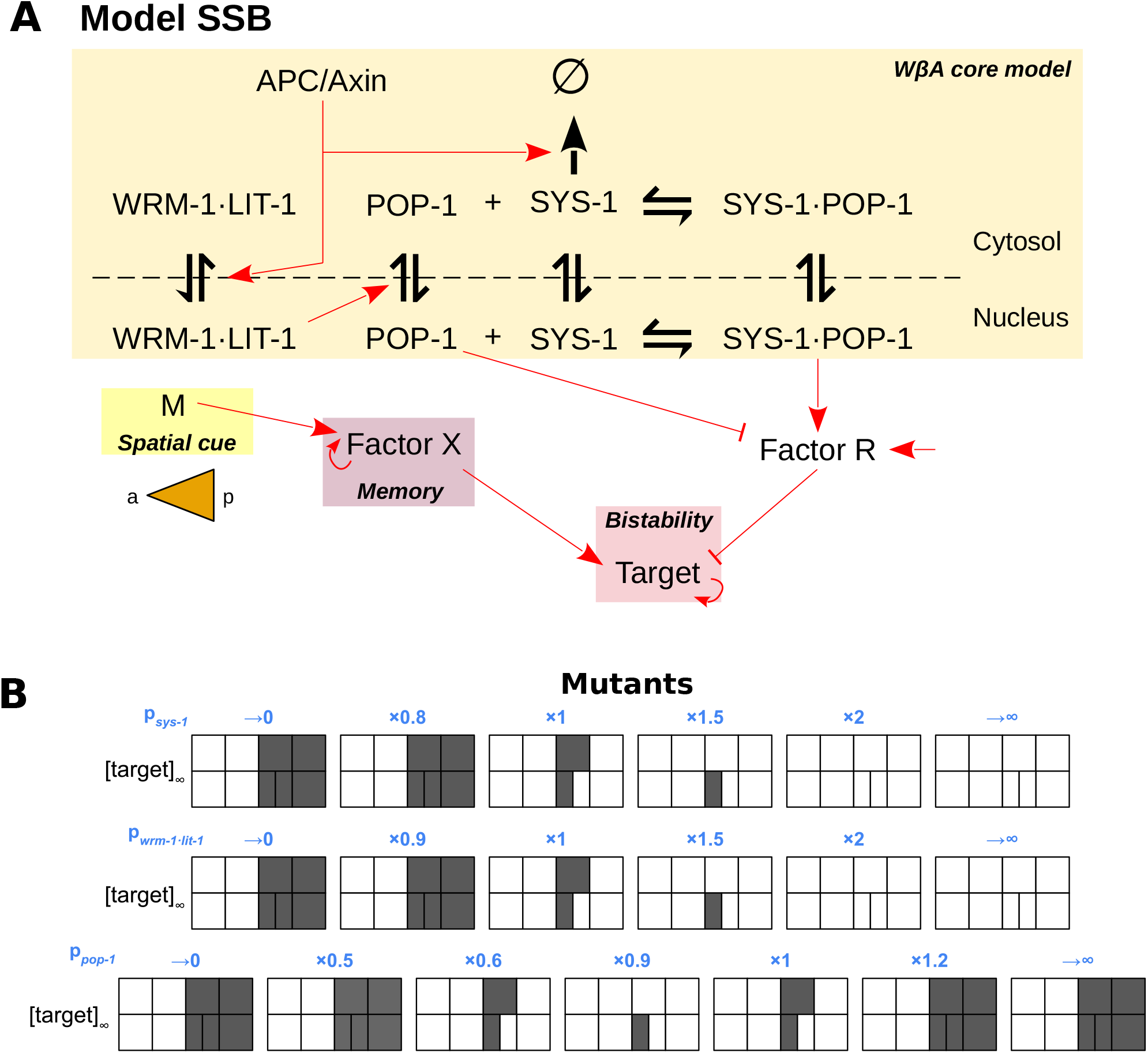
Numerical simulations of the Spatial Symmetry Break (SSB) model. (A) Model diagram. (B) Steady-state concentrations of downstream target at different fold change levels of production rates of SYS-1, POP-1 and LIT-1·WRM-1, *p*_SYS-1_, *p*_POP-1_ and *p*_LIT-1·WRM-1_, relative to reference parametrisation.

## Discussion

The invariance of *C. elegans* cell lineages results from the nematode’s capacity to specify differentiation and proliferation programs with remarkable precision. The molecular basis that underlies such *in vivo* reproducibility has not been properly characterized and could help us better understand how cells are wired to optimally manage biological noise in the context of developmental biology. In this work, we use quantitative modelling to study the GRN involved in fate specification of the postembryonic M lineage in hermaphrodites.

Exploiting the left-right symmetry of the postembryonic M lineage (2), we restricted our analysis to cues in the dorsoventral and anteroposterior developmental axes and introduced a novel and convenient representation for the cellular fate outcomes as 2-D projections. The experimental data on mutants and activity reporters in the LIN-12 Notch pathway are in good alignment with our expectations on the dorso-ventral symmetry break. Ultimately, the foundation of the dorso-ventral axis strictly relies on the ventrally-restricted induction of Notch ligands at the hypodermal cells. How this restriction is achieved is unknown and should be further investigated in the future.

Additionally, we did not address the intriguing observation of LIN-12 that combined full-length and the ICD translational reporters show a degree of anteroposterior asymmetry, whereby M.(d/v)(l/r)a(a/p) express this reporter in a weaker fashion than M.(d/v)(l/r)p(a/p) (11, 47). These observations are unlikely to impact our conclusions since both *lin-12(0)* and *lin-12(gf)* present proper anteroposterior fate specification (Fig 1B). It is unknown to what extent this is a functional feature of the pathway, but it reveals a potential cross-over between the regulation of dorso-ventral and anteroposterior fate specification. In fact, we have not explored the hierarchy of symmetry break decisions in this work. For instance, like our proposed factor X in the TSB model, protein levels for LET-381 and UNC-130 are asymmetrically enriched on the posterior side of the M lineage, but restricted to the dorsal and ventral side, respectively (45, 48). However, whether LET-381 and UNC-130 have similar functional roles to that of X within our modelled GRNs remains uncertain. To address questions regarding hierarchical decision making, future work will attempt to fully integrate dorso-ventral and anteroposterior fate specification in this lineage. We here focused on the anteroposterior symmetry break since our naive model failed to explain how the Wnt/*β*-catenin asymmetry could induce the complete anteroposterior cell fate pattern in the M lineage. Ultimately, this inability results from the asymmetric cell division basis of the W*β*A pathway. Selecting cells that are not in either extreme (M.(d/v)(l/a)aa or M.(d/v)(l/a)pp), requires a change in the underlying regulatory logic as the anteroposterior cell divisions progress. Specifically, to select M.(d/v)(l/r)pa and M.v(l/r)paa, the first division must favor posterior cells whilst the second and third divisions must favor anterior cells. We refer to this phenomenon as a non-monotonic dose-response.

To properly account for the dynamics of different components in time, we used a hybrid continuous-discrete system formalism to combine ODE models of intracellular species with asymmetric cell division. We gradually increased the complexity of our models (W*β*A, W*β*A+B, W*β*A+B+S, W*β*A+B+I, W*β*A+B+I+M, TSB, SSB), harnessing interesting lessons on the underlying logic of fate specification with each new model. Given the high number of parameters, we either focused on i) system properties that were parameter-independent (steady-states, stability) or ii) examined several parameter regimes as case examples. The former required substantial analytical derivations, which are not standard practice for systems this large and are major assets in this work; the latter was approached by tuning parameters such that the kinetic hierarchy of the different subprocesses were plausible and used numerous numerical simulations. In this context, we also highlight the relevance of the observation that POP-1 and SYS-1 translational reporters display anterior and posterior enrichment, respectively. Since the SYS-1·POP-1 complex is expected to fluoresce in both *sys-1::gfp* and *pop-1::gfp* larvae and its constituent monomers have opposite asymmetries, interference can only be avoided if [SYS-1_nuc_](*t*) » [SYS-1·POP-1_nuc_](*t*) «[POP-1_nuc_](*t*). Furthermore, we addressed the question to whether nuclear SYS-1·POP-1 follow the asymmetry of SYS-1, POP-1, or neither. A common preconception in the literature is to assume that SYS-1·POP-1 complex fractions correlate with SYS-1 fractions, since activators (i.e. the SYS-1·POP-1 complex) typically anti-correlate with repressors (POP-1) and that downstream activation is proportional to the nuclear SYS-1/POP-1 ratio. However, increasing the production of either monomer favors complex formation in the form of a saturation curve (**Fig S6**). Thus, this stoichiometric effect can increase downstream activity with an increase in POP-1 under specific circumstances. We used this idea to drive non-monotonicity in the WBA+B+S model, but the resulting dynamics were hypersensitive and failed to distinguish M.v(l/r)paa from M.v(l/r)pap for our chosen parametrisation. Excluding WBA+B+S, all other models seem to support the correlative behavior between SYS-1 and SYS-1·POP-1 fractions. To experimentally verify this, we propose exclusively visualizing the heterodimer form with split fluorophores or with FRET-FLIM imaging.

Under the influence of the Wnt/*β*-catenin pathway, APC/Axin is redistributed asymmetrically among daughter cells dividing in the anteroposterior mitotic axis. However, the consequences of this type of post-translational regulation are expected to be transient in nature unless downstream effects lead to changes in gene regulation. Feedback loops in the W*β*A pathway are not known but could in theory leave an imprint in the form of anteroposterior asymmetries of the components of the destruction complex, experimentally verifiable with translational reporters. Throughout this study though, we assume that APC/Axin asymmetries are transient. Using our core W*β*A model, which integrates known regulatory interactions in this pathway, we showed that downstream components evolve towards a single steady state and that bistability probably resides on networks motifs downstream the W*β*A pathway.

Furthermore, we explored model configurations with increasing complexity to address the limitations of its predecessor: W*β*A+B was unable to generate a non-monotonic response; W*β*A+B+I could properly specify fates but was sensitive to the timing of cell divisions; W*β*A+B+I+M could robustly explain fates in wild type but not in mutants *sys-1(0)* and *pop-1(RNAi)*. Only the final two models, TSB and SSB, could robustly reproduce fate specification in wild type and known W*β*A mutants. TSB uses W*β*A to fully mediate the antero-posterior symmetry break of the lineage, but relies on a temporal cue, P, to limit the time window in which factor X can be turned on. This temporal permissive window effectively inverts the logic between the first and the successive antero-posterior cell divisions. In contrast to TSB, SSB externalizes the decision of the first cell division to another spatial signal, M. Notably, however, the two hypothetical models can be discriminated by a complete loss-of-function of *pop-1*. Thus, The TSB and SSB models are two sides of the same coin; though verifying SSB is much more challenging since it requires identifying an unknown pathway M. Despite numerous previous screening experiments, this pathway could have remained undetected if compromising it leads to earlier embryonic lethality. Indeed, null mutations in *pop-1* itself exhibit such embryonic lethality. To work around this limitation, we propose a cell-specific inducible knock-out of *pop-1* in the M lineage as a highly informative experiment. Removing POP-1 from the network effectively compromises W*β*A pathway transduction. Since the W*β*A pathway is the sole anteroposterior cue in the TSB model, it predicts anteroposterior symmetric fates in two states. If SM and CC fate specification directly or indirectly require POP-1, we expect an ALL-OFF state but an ALL-ON, otherwise. Phenotypically, this might be reflected as randomized fates since the influence of biological noise could potential push the system into either steady-state. This possibility was not incorporated in our models as it would require a different formalism (stochastic differential equations).

Furthermore, our quantitative modelling can help us understand other unrelated mutants. For instance, in *fozi-1(0)* larvae, though fate specification presents inter-individual variation, both dorsal and ventral sublineages generate SMs. However, unlike in *lin-12(gf)* or *sma-9*), SMs are also occasionally formed in the anterior *columns* in cells M.(d/v)(l/r)aa (49). This outcome would be possible in the TSB model if factor X was activated in both M.(d/v)(l/r)a and M.(d/v)(l/r)p and thus, could suggest that FOZI-1 acts as a co-repressor with POP-1 in the anterior inhibition of X. To completely explain the *fozi-1(0)* phenotype, FOZI-1 must also be involved in the dorsoventral symmetry break. Explaining this phenotype with the SSB model would require involving FOZI-1 in the sensing of the hypothetical anteroposterior spatial cue. Such considerations exemplify the utility of our models beyond what they were initially designed for.

On another note, we have so far imposed fixed proliferation patterns in the dorsal and ventral sides. A model that accounts for emergent cell division patterns would have required a substantially more complex modelling approach that integrates cell cycle entry/exit and the interplay between proliferation and differentiation. Such an approach would be required to fully capture the phenotypic alterations in mutants *sys-1(0)* and *pop-1(RNAi)*. In fact, we expect substantial coupling between cell cycle regulation and both dorsoventral and anteroposterior symmetry break. This may not immediately obvious by examining mutants *sem-2(0), ceh-34(RNAi)* and *eya-1(0)* (Fig 1B). Since solely the SM mother (M.v(l/r)pa; *column 3, row 2*) undergoes a third anteroposterior cell division, one could in principle argue that this extra cell division could be just part of the “SM mother” identity. The coupling between the third cell division and SM mother identity is broken in *sem-4(0)* (50) and *fozi-1(0)* mutants (44). In *sem-4(0)*, all M cells differentiate to BWM and both M.d(l/r)pa and M.v(l/r)pa undergo an extra anteroposterior cell division without generating SMs in the process; in *fozi-1(0)*, M.d(l/r)pa becomes SM without an extra cell division whilst M.v(l/r)pa behave as typical SM mother cells. Thus, extra cell divisions can occur without generating SMs and these are not required to generate SMs. Some of the proliferation regulators in the M lineage are known (51). For instance, *mls-2(0)* larvae experience under-proliferation in the M lineage (3). Moreover, other less studied decision-making processes in cell division remain unclear. For instance, much work is needed towards understanding how the sequence of mitotic axes is established (d/v, l/r, a/p [×2 or ×3]) and to explain how mutations in *hlh-8* (52) or *mab-5* (53) lead to alternative axis sequences. Thus, future experimental and theoretical work is needed towards fully understanding the interplay of proliferation and fate diversification in the postembryonic M lineage.

Lastly, we have focused on the sequence of events that occur during the L1 stage. Many aspects remain to be addressed outside this time window. For example, to understand the GRNs involved in the posterior migration the M cell during embryogenesis (54), the anterior migration of SMs during L2 following the EGL-17 FGF gradient originating from the gonad center (55), or the specification of vulval (vm1, vm2) and uterine muscles (um1, um2) in the SM lineage in late L3 and its mirror symmetry (8, 56, 57).

## ACKNOWLEDGEMENTS

We acknowledge Vincent Portegijs (UU), Jonas Mars (Hubrecht), Prof. Jun Liu (Cornell) for critical discussions.

