## Supplementary Methods for "Quantitative modelling of fate specification in the *C. elegans* postembryonic M lineage reveals a missing spatiotemporal signal"

### Supplementary information for quantitative modelling of fate specification in the postembryonic M lineage reveals a missing spatiotemporal signal

Benjamin Planterose Jiménez[1], Alexander R. Blackwell[2], João J. Ramalho[2], Sander van den Heuvel[2], Kirsten ten Tusscher[1], Erika Tsingos[1]


[1] Computational Developmental Biology Group, Utrecht University, Utrecht 3584 CH, The Netherlands
[2] Developmental Biology, Department of Biology, Faculty of Sciences, Utrecht University, Utrecht 3584 CH, The Netherlands

### I. Naive model

The model makes the following assumptions:

- No production nor degradation of APC/Axin
- Cleavage cell divisions
- Constant karyoplasmic ratio
- The activation of downstream Wnt targets, $a^{\text{Wnt}}(t)$, is proportional to $\frac{1}{[\text{APC/Axin}]}$ but also $\frac{[\text{SYS-1}_{\text{nuclear}}]}{[\text{POP-1}_{\text{nuclear}}}$

For each cellular generation $t \in \mathbb{N}$, APC/Axin is distributed among daughter cells with a proportion $\phi$ such that:

$$[\text{APC/Axin}](0) = \frac{n_0^{\text{APC/Axin}}}{V_0} = [\text{APC/Axin}]_0$$

$$\begin{pmatrix} [\text{APC/Axin}]_{\text{a}} \\ [\text{APC/Axin}]_{\text{p}} \end{pmatrix} (1) = [\text{APC/Axin}]_0 \begin{pmatrix} 2 \cdot \phi \\ 2 \cdot (1-\phi) \end{pmatrix}$$

$$\begin{pmatrix} [\text{APC/Axin}]_{\text{aa}} \\ [\text{APC/Axin}]_{\text{ap}} \\ [\text{APC/Axin}]_{\text{pa}} \\ [\text{APC/Axin}]_{\text{pp}} \end{pmatrix} (2) = [\text{APC/Axin}]_0 \cdot \begin{pmatrix} 4 \cdot \phi^2 \\ 4 \cdot \phi \cdot (1-\phi) \\ 4 \cdot \phi \cdot (1-\phi) \\ 4 \cdot (1-\phi)^2 \end{pmatrix}$$

And so on. Based on our assumptions, we can write:

$$a^{\text{Wnt}}(t) = k_1 \cdot R(t) = \frac{k_2}{[\text{APC/Axin}](t)} \tag{1}$$

where $R(t) := \frac{[\text{SYS-1}_{\text{nuclear}}](t)}{[\text{POP-1}_{\text{nuclear}}](t)}$ and $k_1$ and $k_2$ are the corresponding proportionality constants. We thus can write:

$$R(2) = \begin{pmatrix} R_{\text{aa}} \\ R_{\text{ap}} \\ R_{\text{pa}} \\ R_{\text{pp}} \end{pmatrix} (2) = \frac{k_2}{k_1 \cdot [\text{APC/Axin}]_0} \cdot \begin{pmatrix} \frac{1}{4 \cdot \phi^2} \\ \frac{1}{4 \cdot \phi \cdot (1-\phi)} \\ \frac{1}{4 \cdot \phi \cdot (1-\phi)} \\ \frac{1}{4 \cdot (1-\phi)^2} \end{pmatrix}$$

Prior work estimated $\hat{R}(2)$ in the ABpl lineage [6]:

$$\hat{R}(2) = \begin{pmatrix} 0.2 \\ 3.6 \\ 1.1 \\ 9.6 \end{pmatrix}$$

With this data, we can fit $\phi$ by optimizing the following objective function:

$$\hat{\phi} = \arg\max_{\phi} \mathrm{cor}\left(\hat{R}(2), R(2|\phi)\right)$$

where cor is the sample estimate of the Pearson correlation. In the R-programming environment, optimizing with `stats::optim` (method: BFGS) gave the value $\hat{\phi} = 0.7712768 \approx 0.8$. Finally, we estimate $a^{\text{Wnt}}$ relative to $k_2/[\text{APC/Axin}]_0$ in the M lineage for the dorsal and ventral side as:

$$\frac{1}{k_2/[\text{APC/Axin}]_0} \cdot \begin{pmatrix} a_{\text{a}}^{\text{Wnt}} \\ a_{\text{p}}^{\text{Wnt}} \end{pmatrix}(1) = \begin{pmatrix} \frac{1}{2\cdot\hat{\phi}} \\ \frac{1}{2\cdot(1-\hat{\phi})} \end{pmatrix} \approx \begin{pmatrix} 0.62 \\ 2.50 \end{pmatrix}$$

$$\frac{1}{k_2/[\text{APC/Axin}]_0} \cdot \begin{pmatrix} a_{\text{aa}}^{\text{Wnt}} \\ a_{\text{ap}}^{\text{Wnt}} \\ a_{\text{pa}}^{\text{Wnt}} \\ a_{\text{pp}}^{\text{Wnt}} \end{pmatrix}(2) = \begin{pmatrix} \frac{1}{4\cdot\hat{\phi}^2} \\ \frac{1}{4\cdot\hat{\phi}\cdot(1-\hat{\phi})} \\ \frac{1}{4\cdot\hat{\phi}\cdot(1-\hat{\phi})} \\ \frac{1}{4\cdot(1-\hat{\phi})^2} \end{pmatrix} \approx \begin{pmatrix} 0.39 \\ 1.56 \\ 1.56 \\ 6.25 \end{pmatrix}$$

$$\frac{1}{k_2/[\text{APC/Axin}]_0} \cdot \begin{pmatrix} a_{\text{aa}}^{\text{Wnt}} \\ a_{\text{ap}}^{\text{Wnt}} \\ a_{\text{paa}}^{\text{Wnt}} \\ a_{\text{pap}}^{\text{Wnt}} \\ a_{\text{pp}}^{\text{Wnt}} \end{pmatrix}(3) = \begin{pmatrix} \frac{1}{4\cdot\hat{\phi}^2} \\ \frac{1}{4\cdot\hat{\phi}\cdot(1-\hat{\phi})} \\ \frac{1}{8\cdot\hat{\phi}^2\cdot(1-\hat{\phi})} \\ \frac{1}{8\cdot\hat{\phi}\cdot(1-\hat{\phi})^2} \\ \frac{1}{4\cdot(1-\hat{\phi})^2} \end{pmatrix} \approx \begin{pmatrix} 0.39 \\ 1.56 \\ 0.98 \\ 3.91 \\ 6.25 \end{pmatrix}$$

Since

$$[\text{APC/Axin}] = \frac{k_2}{a^{\text{Wnt}}}$$

We define

$$\mathcal{A}^{\text{Wnt}} := \frac{a^{\text{Wnt}}}{[\text{APC/Axin}]_0} = \frac{[\text{APC/Axin}]_0}{[\text{APC/Axin}](t)}$$

$$\mathcal{F}_\beta = \mathbb{1}_{\mathcal{A}^{\text{Wnt}} > \mathcal{A}^{\text{Wnt}}_{\text{critical}}}; \quad \mathcal{F}_\alpha = 1 - \mathcal{F}_\beta$$

#### II. General parameters

##### A. Cell volume of the M cell

We use three different approximations to estimate the volume of the M cell:

*1) Direct measurement from Nomarski interference contrast:* We used the scaled Nomarski micrograph of an M cell in a young L1 hermaphrodite (Fig 6A in [5]). We used the ImageJ's elliptical selection tool to measure the major and minor diameter. Cell were approximately spherical so we averaged out both major and minor diameters obtaining:

$$\hat{R}_{\text{cell}}^{\text{M}} = 2.19\ \mu\text{m}$$

which corresponds to:

$$\hat{V}_{\text{cell}}^{\text{M}} = \frac{4}{3} \cdot \pi \cdot \hat{R}_{\text{M}}^{3} = 43.71\ \mu\text{m}^3$$

*2) Using cell volumes at early embryonic development:* The M cell is generated in early development (MS.apaapp), where MS is P0.paa. X. Kuang *et al* [3] have made precise measurements of cell volumes in early embryogenesis using 3D time-lapse confocal microscopy, including $\hat{V}_{\text{cell}}^{\text{MS}} = 2389.470\ \mu\text{m}^3$. Assuming that $\phi_{\text{cell}}$ remains close to $1/2$ in the MS lineage and since the M cell is MS.apaapp, we propose the following approximation:

$$\hat{V}_{\text{cell}}^{\text{M}} = \hat{V}_{\text{cell}}^{\text{MS}} \cdot \left(\frac{1}{2}\right)^{6} = 37.3\ \mu\text{m}^3$$

that, assuming spherical geometry, corresponds to a radius of:

$$\hat{R}_{\text{cell}}^{\text{M}} = \sqrt[3]{\frac{3 \cdot \hat{V}_{\text{cell}}^{\text{M}}}{4 \cdot \pi}} = 2.1\ \mu\text{m}$$

*3) Using the radius of a worm as a reference:* The M cell approximately occupies $\frac{1}{3}$ of a worm's width (early hermaphrodite L1 *C. elegans*; strain N2). In other words: $D_{\text{worm}} \approx 6 \cdot R_{\text{cell}}^{\text{M}}$. Early hermaphodite L1 *C. elegans*; strain N2 have a diameter of around 13 $\mu$m [1]. Thus:

$$\hat{R}_{\text{cell}}^{\text{M}} = \frac{\hat{D}_{\text{worm}}}{6} = 2.17\ \mu\text{m}$$

which, assuming spherical geometry, corresponds to a volume of:

$$\hat{V}_{\text{cell}}^{\text{M}} = \frac{4}{3} \cdot \pi \cdot \hat{R}_{\text{M}}^{3} = 42.60\ \mu\text{m}^3$$

*4) Combining estimates:* Since for our aim, we do not require a precise measurement but simply a value in the same order of magnitude, we propose to set $\hat{V}_{\text{M}} := 40\ \mu\text{m}^3$ and $\hat{R}_{\text{M}} := \sqrt[3]{\frac{30}{\pi}}\ \mu\text{m} \approx 2.12\ \mu\text{m}$.

##### B. Nuclear volume

*1) Direct measurement from Nomarski interference contrast:* As for II-A1, we used the scaled Nomarski micrograph of an M cell in a young L1 hermaphrodite (Fig 6A in [5]) to estimate nuclear volume. The major and minor diameters we slightly different (1.48 $\mu$m Vs 1.32 $\mu$m), but we decided to ignore these differences for the sake of simplicity ($\approx$ 1.41 $\mu$m), resulting in:

$$\hat{V}^{\mathrm{M}}_{\mathrm{nuc}} = \frac{4}{3} \cdot \pi \cdot (\hat{R}^{\mathrm{M}}_{\mathrm{nuc}})^3 = 11.64\ \mu\mathrm{m}^3$$

which corresponds to a karyoplasmic ratio of $\frac{\hat{V}^{\mathrm{M}}_{\mathrm{nuc}}}{\hat{V}^{\mathrm{M}}_{\mathrm{cell}}} \approx 0.27$.

##### C. Cellular and nuclear volume of M.(d/v)(l/r)

In this work, we focus on the anterior-posterior cell divisions. At this point, two cell divisions have occurred already (d/v, l/r), thus, assuming cleavage-like cell divisions:

$$V^{\mathrm{cell}}_0 := \hat{V}^{\mathrm{M}}_{\mathrm{cell}} \cdot (1/2)^2 = 10\ \mu\mathrm{m}^3$$

And assuming the same karyoplasmic ratio as for the M cell:

$$V^{\mathrm{nuc}}_0 := V^{\mathrm{cell}}_0 \cdot \left( \frac{\hat{V}^{\mathrm{M}}_{\mathrm{nuc}}}{\hat{V}^{\mathrm{M}}_{\mathrm{cell}}} \right) = 2.7\ \mu\mathrm{m}^3$$

$$V^{\mathrm{cyt}}_0 := V^{\mathrm{cell}}_0 - V^{\mathrm{nuc}}_0 = 7.3\ \mu\mathrm{m}^3$$

which corresponds to:

$$R^{\mathrm{nuc}}_0 := \sqrt[3]{\frac{3 \cdot V^{\mathrm{nuc}}_0}{4 \cdot \pi}} \approx 0.86\ \mu\mathrm{m}$$

$$\hat{A}^{\mathrm{nuc}}_0 = 4 \cdot \pi \cdot (\hat{R}^{\mathrm{M}}_{\mathrm{nuc}})^2 \approx 9.38\ \mu\mathrm{m}^2$$

##### D. Cell division timing

Cell divisions begin at 6.5 hours after hatching and they are approximately spaced every 2 hours [5]:

| Mitotic Axis | Which cells divide? | Time of Cell division, 20°C [Time since hatching (h)] |
|---|---|---|
| (d/v) | M | 6.5 |
| (l/r) | M.(d/v) | 8.5 |
| (a/p) | M.(d/v)(l/r) | 10.5 |
| (a/p) | M.(d/v)(l/r)(a/p) | 12.5 |
| (a/p) | M.v(l/r)pa | 14.5 |

Since in this work we solely model the anterior-posterior cell divisions, we set the reference 1 hour after the (l/r) cell division (or 9.5 hours after hatching) which results in:

| Mitotic axis | Which cells divide? | Time of Cell division, 20°C [Time since hatching (min)] |
|---|---|---|
| (a/p) | M.(d/v)(l/r) | 60 |
| (a/p) | M.(d/v)(l/r)(a/p) | 180 |
| (a/p) | M.v(l/r)pa | 300 |

#### III. WNT/$\beta$-CATENIN ASYMMETRY PATHWAY CORE MODEL [W$\beta$A]

The model makes the following assumptions:

- Continuous time
- APC/Axin constant production and first-order mass-action law kinetics for degradation
- Cleavage cell divisions with constant karyoplasmic ratio

##### A. *Differential equations*

Let $i \in \{0, 1, \ldots, N_{\text{division}}\}$, where $N_{\text{division}}$ is the maximum number of cell divisions experienced across the cells included in the lineage, let $\tau_i$ denote the time of cell division $i$, sorted such that $\tau_{i+1} > \tau_i$ and $\tau_0 := 0$ and let $j$ be an index that runs across the total number of cellular product at time-infinity (i.e. total number of paths). For clarity, $j \in \{\text{aa}, \text{ap}, \text{pa}, \text{pp}\}$ for the dorsal side and $j \in \{\text{aa}, \text{ap}, \text{paa}, \text{pap}, \text{pp}\}$ for the ventral side. Additionally, let us define the status of the cell $j$ at time $t$, $s_j(t)$ as:

$$\text{if } t \in [\tau_k, \tau_{k+1}) : s_j(t) := \begin{cases} \{n\} & \text{if cell } j \text{ is not a cellular product of a cell division occurring at } \tau_k \\ \{a\} & \text{if cell } j \text{ is the anterior cellular product of a cell division occurring at } \tau_k \\ \{p\} & \text{if cell } j \text{ is the posterior cellular product of a cell division occurring at } \tau_k \end{cases}$$

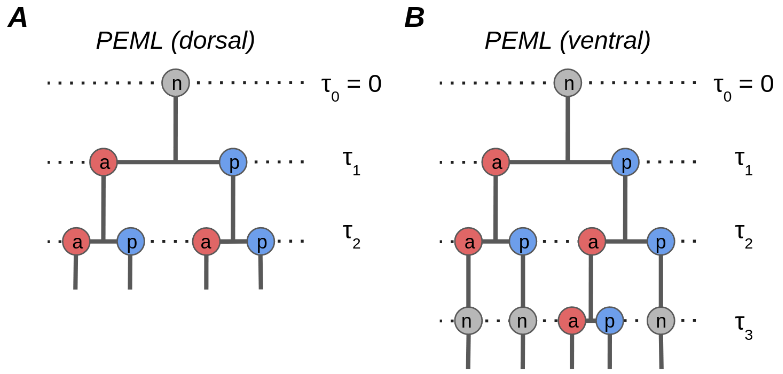


**Fig. SM1:** Definition of $s_j(t)$ for the dorsal and ventral cell proliferation scheme in the postembryonic M lineage (PEML). (A) There are a total of 4 paths and 2 cell divisions. (B) There are a total of 5 paths and 3 cell divisions.

We propose the following system of differential equations (eq 13-26), defined for each path $j$:

$$\frac{d(V_{\text{cell}})_j}{dt} = -(V_{\text{cell}})_j/2 \cdot \left( \sum_{i=1}^{N_{\text{division}}} \delta(t - \tau_i) \right) \cdot \mathbb{1}_{\{a,p\}}(s_j(t)) \tag{2}$$

where $\mathbb{1}_A(x)$ denotes an indicator function, equal to 1 when $x \in A$ but equal to 0 otherwise and $\delta(x)$ is the Dirac delta function.

$$\frac{d[\text{APC}\cdot\text{Axin}]_j}{dt} = p_{\text{APC}\cdot\text{Axin}} - d_{\text{APC}\cdot\text{Axin}} \cdot [\text{APC}\cdot\text{Axin}]_j + \\ - [\text{APC}\cdot\text{Axin}]_j \cdot \left[1 - 2\cdot\phi\cdot\mathbb{1}_{\{a\}}\left(s_j(t)\right) - 2\cdot(1-\phi)\cdot\mathbb{1}_{\{p\}}\left(s_j(t)\right)\right] \cdot \left(\sum_{i=1}^{N_{\text{division}}} \delta(t-\tau_i)\right) \cdot \mathbb{1}_{\{a,p\}}\left(s_j(t)\right) \tag{3}$$

where $\phi \in [0.5, 1]$ is the APC/Axin partition enrichment coefficient for the anterior sister cell product.

$$\frac{d[\text{LIT-1}\cdot\text{WRM-1}_{\text{cyt}}]_j}{dt} = p_{\text{LIT-1}\cdot\text{WRM-1}} - d_{\text{LIT-1}\cdot\text{WRM-1}} \cdot [\text{LIT-1}\cdot\text{WRM-1}_{\text{cyt}}]_j + \\ \frac{(A_{\text{nuc}})_j}{(V_{\text{cell}})_j - (V_{\text{nuc}})_j} \cdot \left(K^{\text{nuc:cyt}}_{\text{LIT-1}\cdot\text{WRM-1}} \cdot [\text{LIT-1}\cdot\text{WRM-1}_{\text{nuc}}]_j \cdot \frac{[\text{APC}\cdot\text{Axin}]_j^2}{[\text{APC}\cdot\text{Axin}]_j^2 + K^2_{\text{APC}\cdot\text{Axin:LIT-1}\cdot\text{WRM-1}}} - K^{\text{cyt:nuc}}_{\text{LIT-1}\cdot\text{WRM-1}} \cdot [\text{LIT-1}\cdot\text{WRM-1}_{\text{cyt}}]_j\right) \tag{4}$$

where $(V_{\text{nuc}})_j := 0.27 \cdot (V_{\text{cell}})_j$ (constant karyoplasmic ratio) and $(A_{\text{nuc}})_j = 4\cdot\pi\cdot\left(\sqrt[3]{\frac{3\cdot(V_{\text{nuc}})_j}{4\cdot\pi}}\right)^2$.

$$\frac{d[\text{LIT-1}\cdot\text{WRM-1}_{\text{nuc}}]_j}{dt} = \frac{(A_{\text{nuc}})_j}{(V_{\text{nuc}})_j} \cdot \left(K^{\text{cyt:nuc}}_{\text{LIT-1}\cdot\text{WRM-1}} \cdot [\text{LIT-1}\cdot\text{WRM-1}_{\text{cyt}}]_j - K^{\text{nuc:cyt}}_{\text{LIT-1}\cdot\text{WRM-1}} \cdot [\text{LIT-1}\cdot\text{WRM-1}_{\text{nuc}}]_j \cdot \frac{[\text{APC}\cdot\text{Axin}]_j^2}{[\text{APC}\cdot\text{Axin}]_j^2 + K^2_{\text{APC}\cdot\text{Axin:LIT-1}\cdot\text{WRM-1}}}\right) \tag{5}$$

$$\frac{d[\text{SYS-1}_{\text{cyt}}]_j}{dt} = p_{\text{SYS-1}} - d_{\text{SYS-1}} \cdot [\text{SYS-1}_{\text{cyt}}]_j - d_{\text{APC}\cdot\text{Axin:SYS-1}} \cdot [\text{SYS-1}_{\text{cyt}}]_j \cdot \frac{[\text{APC}\cdot\text{Axin}]_j^2}{[\text{APC}\cdot\text{Axin}]_j^2 + K^2_{\text{APC}\cdot\text{Axin:SYS-1}}} + \\ - K^{\text{on}}_{\text{SYS-1}\cdot\text{POP-1}} \cdot [\text{SYS-1}_{\text{cyt}}]_j \cdot [\text{POP-1}_{\text{cyt}}]_j + K^{\text{off}}_{\text{SYS-1}\cdot\text{POP-1}} \cdot [\text{SYS-1}\cdot\text{POP-1}_{\text{cyt}}]_j + \frac{(A_{\text{nuc}})_j}{(V_{\text{cell}})_j - (V_{\text{nuc}})_j} \cdot \left(K^{\text{nuc:cyt}}_{\text{SYS-1}} \cdot [\text{SYS-1}_{\text{nuc}}]_j - K^{\text{cyt:nuc}}_{\text{SYS-1}} \cdot [\text{SYS-1}_{\text{cyt}}]_j\right) \tag{6}$$

$$\frac{d[\text{SYS-1}_{\text{nuc}}]_j}{dt} = -K^{\text{on}}_{\text{SYS-1}\cdot\text{POP-1}} \cdot [\text{SYS-1}_{\text{nuc}}]_j \cdot [\text{POP-1}_{\text{nuc}}]_j + K^{\text{off}}_{\text{SYS-1}\cdot\text{POP-1}} \cdot [\text{SYS-1}\cdot\text{POP-1}_{\text{nuc}}]_j + \\ \frac{(A_{\text{nuc}})_j}{(V_{\text{nuc}})_j} \cdot \left(K^{\text{cyt:nuc}}_{\text{SYS-1}} \cdot [\text{SYS-1}_{\text{cyt}}]_j - K^{\text{nuc:cyt}}_{\text{SYS-1}} \cdot [\text{SYS-1}_{\text{nuc}}]_j\right) \tag{7}$$

$$\frac{d[\text{POP-1}_{\text{cyt}}]_j}{dt} = p_{\text{POP-1}} - d_{\text{POP-1}} \cdot [\text{POP-1}_{\text{cyt}}]_j - K^{\text{on}}_{\text{SYS-1}\cdot\text{POP-1}} \cdot [\text{SYS-1}_{\text{cyt}}]_j \cdot [\text{POP-1}_{\text{cyt}}]_j + K^{\text{off}}_{\text{SYS-1}\cdot\text{POP-1}} \cdot [\text{SYS-1}\cdot\text{POP-1}_{\text{cyt}}]_j + \\ \frac{(A_{\text{nuc}})_j}{(V_{\text{cell}})_j - (V_{\text{nuc}})_j} \cdot \left(K^{\text{nuc:cyt}}_{\text{POP-1}} \cdot [\text{POP-1}_{\text{nuc}}]_j \cdot \frac{[\text{LIT-1}\cdot\text{WRM-1}_{\text{nuc}}]_j^2}{[\text{LIT-1}\cdot\text{WRM-1}_{\text{nuc}}]_j^2 + K^2_{\text{LIT-1}\cdot\text{WRM-1:POP-1}}} - K^{\text{cyt:nuc}}_{\text{POP-1}} \cdot [\text{POP-1}_{\text{cyt}}]_j\right) \tag{8}$$

$$\frac{d[\text{POP-1}_{\text{nuc}}]_j}{dt} = -K^{\text{on}}_{\text{SYS-1}\cdot\text{POP-1}} \cdot [\text{SYS-1}_{\text{nuc}}]_j \cdot [\text{POP-1}_{\text{nuc}}]_j + K^{\text{off}}_{\text{SYS-1}\cdot\text{POP-1}} \cdot [\text{SYS-1}\cdot\text{POP-1}_{\text{nuc}}]_j + \\ \frac{(A_{\text{nuc}})_j}{(V_{\text{nuc}})_j} \cdot \left(K^{\text{cyt:nuc}}_{\text{POP-1}} \cdot [\text{POP-1}_{\text{cyt}}]_j - K^{\text{nuc:cyt}}_{\text{POP-1}} \cdot [\text{POP-1}_{\text{nuc}}]_j \cdot \frac{[\text{LIT-1}\cdot\text{WRM-1}_{\text{nuc}}]_j^2}{[\text{LIT-1}\cdot\text{WRM-1}_{\text{nuc}}]_j^2 + K^2_{\text{LIT-1}\cdot\text{WRM-1:POP-1}}}\right) \tag{9}$$

$$\frac{d[\text{SYS-1}\cdot\text{POP-1}_{\text{cyt}}]_j}{dt} = K^{\text{on}}_{\text{SYS-1}\cdot\text{POP-1}} \cdot [\text{SYS-1}_{\text{cyt}}]_j \cdot [\text{POP-1}_{\text{cyt}}]_j - K^{\text{off}}_{\text{SYS-1}\cdot\text{POP-1}} \cdot [\text{SYS-1}\cdot\text{POP-1}_{\text{cyt}}]_j - d_{\text{SYS-1}\cdot\text{POP-1}} \cdot [\text{SYS-1}\cdot\text{POP-1}_{\text{cyt}}]_j + \frac{(A_{\text{nuc}})_j}{(V_{\text{cell}})_j - (V_{\text{nuc}})_j} \cdot \left(K^{\text{nuc:cyt}}_{\text{SYS-1}\cdot\text{POP-1}} \cdot [\text{SYS-1}\cdot\text{POP-1}_{\text{nuc}}]_j - K^{\text{cyt:nuc}}_{\text{SYS-1}\cdot\text{POP-1}} \cdot [\text{SYS-1}\cdot\text{POP-1}_{\text{cyt}}]_j\right) \tag{10}$$

$$\frac{d[\text{SYS-1}\cdot\text{POP-1}_{\text{nuc}}]_j}{dt} = K^{\text{on}}_{\text{SYS-1}\cdot\text{POP-1}} \cdot [\text{SYS-1}_{\text{nuc}}]_j \cdot [\text{POP-1}_{\text{nuc}}]_j - K^{\text{off}}_{\text{SYS-1}\cdot\text{POP-1}} \cdot [\text{SYS-1}\cdot\text{POP-1}_{\text{nuc}}]_j + \frac{(A_{\text{nuc}})_j}{(V_{\text{nuc}})_j} \cdot \left(K^{\text{cyt:nuc}}_{\text{SYS-1}\cdot\text{POP-1}} \cdot [\text{SYS-1}\cdot\text{POP-1}_{\text{cyt}}]_j - K^{\text{nuc:cyt}}_{\text{SYS-1}\cdot\text{POP-1}} \cdot [\text{SYS-1}\cdot\text{POP-1}_{\text{nuc}}]_j\right) \tag{11}$$

The nuclear:cytosolic import/export was set-up according to thermodynamically-consistent modelling standards [2], but assuming that the kinetic constants have been corrected for the fraction of nuclear pore area to total nucleus area, which we assume to be invariant upon cell division.

##### *B. Steady-states*

$$(V_{\text{cell}})_{j,\infty} = V_0^{\text{cell}} \cdot 2^{-\int_{t=0}^{\infty}\left(\sum_{i=1}^{N_{\text{division}}} \delta(t-\tau_i)\right)\cdot \mathbb{1}_{\{a,p\}}(s_j(t))\cdot dt} \tag{12}$$

$$(V_{\text{nuc}})_{j,\infty} = 0.27 \cdot (V_{\text{cell}})_{j,\infty} \tag{13}$$

$$(A_{\text{nuc}})_{j,\infty} = 4 \cdot \pi \cdot \left(\sqrt[3]{\frac{3 \cdot (V_{\text{nuc}})_{j,\infty}}{4 \cdot \pi}}\right)^2 \tag{14}$$

$$[\text{APC}\cdot\text{Axin}]_\infty = \frac{p_{\text{APC}\cdot\text{Axin}}}{d_{\text{APC}\cdot\text{Axin}}} \tag{15}$$

$$[\text{LIT-1}\cdot\text{WRM-1}_{\text{cyt}}]_\infty = \frac{p_{\text{LIT-1}\cdot\text{WRM-1}}}{d_{\text{LIT-1}\cdot\text{WRM-1}}} \tag{16}$$

$$[\text{LIT-1}\cdot\text{WRM-1}_{\text{nuc}}]_\infty = \frac{K^{\text{cyt:nuc}}_{\text{LIT-1}\cdot\text{WRM-1}} \cdot [\text{LIT-1}\cdot\text{WRM-1}_{\text{cyt}}]_\infty \cdot \left([\text{APC}\cdot\text{Axin}]^2_\infty + K^2_{\text{APC}\cdot\text{Axin:LIT-1}\cdot\text{WRM-1}}\right)}{K^{\text{nuc:cyt}}_{\text{LIT-1}\cdot\text{WRM-1}} \cdot [\text{APC}\cdot\text{Axin}]^2_\infty} \tag{17}$$

For the remaining components, we first propose the following balance equations:

$$V_{\text{cyto}} \cdot \left(\frac{d[\text{SYS-1}_{\text{cyt}}]_j}{dt} + \frac{d[\text{SYS-1}\cdot\text{POP-1}_{\text{cyt}}]_j}{dt}\right) + V_{\text{nuc}} \cdot \left(\frac{d[\text{SYS-1}_{\text{nuc}}]_j}{dt} + \frac{d[\text{SYS-1}\cdot\text{POP-1}_{\text{nuc}}]_j}{dt}\right) = 0$$

$$V_{\text{cyto}} \cdot \left( \frac{d[\text{POP-1}_{\text{cyt}}]_j}{dt} + \frac{d[\text{SYS-1}{\cdot}\text{POP-1}_{\text{cyt}}]_j}{dt} \right) + V_{\text{nuc}} \cdot \left( \frac{d[\text{POP-1}_{\text{nuc}}]_j}{dt} + \frac{d[\text{SYS-1}{\cdot}\text{POP-1}_{\text{nuc}}]_j}{dt} \right) = 0$$

which result in

$$V_{\text{cyto}} \cdot (p_{\text{SYS-1}} - \alpha \cdot [\text{SYS-1}_{\text{cyt}}]_\infty - d_{\text{SYS-1}\cdot\text{POP-1}} \cdot [\text{SYS-1}{\cdot}\text{POP-1}_{\text{cyt}}]_\infty) = 0 \tag{18}$$

where $\alpha := d_{\text{SYS-1}} + d_{\text{APC}\cdot\text{Axin:SYS-1}} \cdot \frac{[\text{APC}\cdot\text{Axin}]_j^2}{[\text{APC}\cdot\text{Axin}]_j^2 + K^2_{\text{APC}\cdot\text{Axin:SYS-1}}}$, and

$$V_{\text{cyto}} \cdot (p_{\text{POP-1}} - d_{\text{POP-1}} \cdot [\text{POP}_{\text{cyt}}]_\infty - d_{\text{SYS-1}\cdot\text{POP-1}} \cdot [\text{SYS-1}{\cdot}\text{POP-1}_{\text{cyt}}]_\infty) = 0 \tag{19}$$

Using the balance results, we can sequentially solve for the steady-states of the remaining components; defining $x := [\text{SYS-1}_{\text{cyt}}]_\infty$, $y := [\text{POP-1}_{\text{cyt}}]_\infty$, $z := [\text{SYS-1}{\cdot}\text{POP-1}_{\text{cyt}}]_\infty$, $a := [\text{SYS-1}_{\text{nuc}}]_\infty$, $b := [\text{POP-1}_{\text{nuc}}]_\infty$, $c := [\text{SYS-1}{\cdot}\text{POP-1}_{\text{nuc}}]_\infty$, we use the following sequence to solve the steady-states with the help of symbolic calculus:

- Use Eq. 18 to write $z = f(y)$
- Use above and Eq. 19 to write $x = f(y)$
- substitute all of the above on Eq. 6 to write $a = f(y)$
- substitute all of the above on Eq. 8 to write $b = f(y)$
- substitute all of the above on Eq. 10 to write $c = f(y)$
- substitute all of the above on Eq. 11 to end up with a polynomial as a function of $y$.

This procedure results in the following polynomial:

$$a_4 \cdot y^4 + a_3 \cdot y^3 + a_2 \cdot y^2 + a_1 \cdot y + a_0 = 0 \tag{20}$$

where:

$$a_4 = \frac{{K^{\text{on}}_{\text{SYS-1}\cdot\text{POP-1}}}^3 \cdot V^2_{\text{cyt}} \cdot d^2_{\text{POP-1}}}{A^2_{\text{nuc}} \cdot K^{\text{nuc:cyt}}_{\text{SYS-1}} \cdot \alpha^2 \cdot \beta}$$

where $\beta := K^{\text{nuc:cyt}}_{\text{POP-1}} \cdot \frac{[\text{LIT-1}\cdot\text{WRM-1}_{\text{nuc}}]^2_\infty}{[\text{LIT-1}\cdot\text{WRM-1}_{\text{nuc}}]^2_\infty + K^2_{\text{LIT-1}\cdot\text{WRM-1:POP-1}}}$, $V_{\text{cyt}} := (V_{\text{cell}})_{j,\infty} - (V_{\text{nuc}})_{j,\infty}$ and $A_{\text{nuc}} = (A_{\text{nuc}})_{j,\infty}$

$$a_3 = \frac{{K^{\text{on}}_{\text{SYS-1}\cdot\text{POP-1}}}^2 V_{\text{cyt}} d_{\text{POP-1}}}{A^2_{\text{nuc}} K^{\text{nuc:cyt}}_{\text{SYS-1}} \alpha^2 \beta d_{\text{SYS-1}\cdot\text{POP-1}}} [A_{\text{nuc}} K^{\text{cyt:nuc}}_{\text{POP-1}} \alpha d_{\text{SYS-1}\cdot\text{POP-1}} + A_{\text{nuc}} K^{\text{cyt:nuc}}_{\text{SYS-1}} d_{\text{POP-1}} d_{\text{SYS-1}\cdot\text{POP-1}} + 2K^{\text{off}}_{\text{SYS-1}\cdot\text{POP-1}} V_{\text{cyt}} \alpha d_{\text{POP-1}} - 2K^{\text{on}}_{\text{SYS-1}\cdot\text{POP-1}} V_{\text{cyt}} d_{\text{SYS-1}\cdot\text{POP-1}} p_{\text{POP-1}} + 2K^{\text{on}}_{\text{SYS-1}\cdot\text{POP-1}} V_{\text{cyt}} d_{\text{SYS-1}\cdot\text{POP-1}} p_{\text{SYS-1}} + 2V_{\text{cyt}} \alpha d_{\text{POP-1}} d_{\text{SYS-1}\cdot\text{POP-1}}]$$

$$a_2 = \frac{K^{\text{POP-1}}_{\text{cyt:nuc}} K^{\text{SYS-1}}_{\text{cyt:nuc}} K^{\text{on}}_{\text{SYS-1·POP-1}} d_{\text{POP-1}}}{K^{\text{SYS-1}}_{\text{nuc:cyt}} \alpha\beta} + \frac{K^{\text{on}}_{\text{SYS-1·POP-1}} V_{\text{cyto}} d_{\text{POP-1}}}{V_{\text{nuc}}\alpha} + \frac{K^{\text{POP-1}}_{\text{cyt:nuc}} K^{\text{off}}_{\text{SYS-1·POP-1}} K^{\text{on}}_{\text{SYS-1·POP-1}} V_{\text{cyto}} d_{\text{POP-1}}}{A_{\text{nuc}} K^{\text{SYS-1}}_{\text{nuc:cyt}} \beta d_{\text{SYS-1·POP-1}}} - \frac{K^{\text{POP-1}}_{\text{cyt:nuc}} {K^{\text{on}}_{\text{SYS-1·POP-1}}}^2 V_{\text{cyto}} p_{\text{POP-1}}}{A_{\text{nuc}} K^{\text{SYS-1}}_{\text{nuc:cyt}} \alpha\beta} + \frac{K^{\text{POP-1}}_{\text{cyt:nuc}} {K^{\text{on}}_{\text{SYS-1·POP-1}}}^2 V_{\text{cyto}} p_{\text{SYS-1}}}{A_{\text{nuc}} K^{\text{SYS-1}}_{\text{nuc:cyt}} \alpha\beta} +$$
$$\frac{K^{\text{POP-1}}_{\text{cyt:nuc}} K^{\text{on}}_{\text{SYS-1·POP-1}} V_{\text{cyto}} d_{\text{POP-1}}}{A_{\text{nuc}} K^{\text{SYS-1}}_{\text{nuc:cyt}} \beta} + \frac{K^{\text{SYS-1}}_{\text{cyt:nuc}} K^{\text{off}}_{\text{SYS-1·POP-1}} K^{\text{on}}_{\text{SYS-1·POP-1}} V_{\text{cyto}} d^2_{\text{POP-1}}}{A_{\text{nuc}} K^{\text{SYS-1}}_{\text{nuc:cyt}} \alpha\beta d_{\text{SYS-1·POP-1}}} - \frac{2 K^{\text{SYS-1}}_{\text{cyt:nuc}} {K^{\text{on}}_{\text{SYS-1·POP-1}}}^2 V_{\text{cyto}} d_{\text{POP-1}} p_{\text{POP-1}}}{A_{\text{nuc}} K^{\text{SYS-1}}_{\text{nuc:cyt}} \alpha^2\beta} +$$
$$\frac{2 K^{\text{SYS-1}}_{\text{cyt:nuc}} {K^{\text{on}}_{\text{SYS-1·POP-1}}}^2 V_{\text{cyto}} d_{\text{POP-1}} p_{\text{SYS-1}}}{A_{\text{nuc}} K^{\text{SYS-1}}_{\text{nuc:cyt}} \alpha^2\beta} + \frac{K^{\text{SYS-1}}_{\text{cyt:nuc}} K^{\text{on}}_{\text{SYS-1·POP-1}} V_{\text{cyto}} d^2_{\text{POP-1}}}{A_{\text{nuc}} K^{\text{SYS-1}}_{\text{nuc:cyt}} \alpha\beta} + \frac{K^{\text{off}}_{\text{SYS-1·POP-1}} K^{\text{on}}_{\text{SYS-1·POP-1}} V_{\text{cyto}} d_{\text{POP-1}}}{A_{\text{nuc}} K^{\text{SYS-1·POP-1}}_{\text{nuc:cyt}} \alpha} + \frac{{K^{\text{off}}_{\text{SYS-1·POP-1}}}^2 K^{\text{on}}_{\text{SYS-1·POP-1}} V^2_{\text{cyto}} d^2_{\text{POP-1}}}{A^2_{\text{nuc}} K^{\text{SYS-1}}_{\text{nuc:cyt}} \beta d^2_{\text{SYS-1·POP-1}}} +$$
$$- \frac{4 K^{\text{off}}_{\text{SYS-1·POP-1}} {K^{\text{on}}_{\text{SYS-1·POP-1}}}^2 V^2_{\text{cyto}} d_{\text{POP-1}} p_{\text{POP-1}}}{A^2_{\text{nuc}} K^{\text{SYS-1}}_{\text{nuc:cyt}} \alpha\beta d_{\text{SYS-1·POP-1}}} + \frac{2 K^{\text{off}}_{\text{SYS-1·POP-1}} {K^{\text{on}}_{\text{SYS-1·POP-1}}}^2 V^2_{\text{cyto}} d_{\text{POP-1}} p_{\text{SYS-1}}}{A^2_{\text{nuc}} K^{\text{SYS-1}}_{\text{nuc:cyt}} \alpha\beta d_{\text{SYS-1·POP-1}}} + \frac{2 K^{\text{off}}_{\text{SYS-1·POP-1}} K^{\text{on}}_{\text{SYS-1·POP-1}} V^2_{\text{cyto}} d^2_{\text{POP-1}}}{A^2_{\text{nuc}} K^{\text{SYS-1}}_{\text{nuc:cyt}} \beta d_{\text{SYS-1·POP-1}}} + \frac{{K^{\text{on}}_{\text{SYS-1·POP-1}}}^3 V^2_{\text{cyto}} p^2_{\text{POP-1}}}{A^2_{\text{nuc}} K^{\text{SYS-1}}_{\text{nuc:cyt}} \alpha^2\beta} +$$
$$- \frac{2 {K^{\text{on}}_{\text{SYS-1·POP-1}}}^3 V^2_{\text{cyto}} p_{\text{POP-1}} p_{\text{SYS-1}}}{A^2_{\text{nuc}} K^{\text{SYS-1}}_{\text{nuc:cyt}} \alpha^2\beta} + \frac{{K^{\text{on}}_{\text{SYS-1·POP-1}}}^3 V^2_{\text{cyto}} p^2_{\text{SYS-1}}}{A^2_{\text{nuc}} K^{\text{SYS-1}}_{\text{nuc:cyt}} \alpha^2\beta} - \frac{4 {K^{\text{on}}_{\text{SYS-1·POP-1}}}^2 V^2_{\text{cyto}} d_{\text{POP-1}} p_{\text{POP-1}}}{A^2_{\text{nuc}} K^{\text{SYS-1}}_{\text{nuc:cyt}} \alpha\beta} + \frac{2 {K^{\text{on}}_{\text{SYS-1·POP-1}}}^2 V^2_{\text{cyto}} d_{\text{POP-1}} p_{\text{SYS-1}}}{A^2_{\text{nuc}} K^{\text{SYS-1}}_{\text{nuc:cyt}} \alpha\beta} + \frac{K^{\text{on}}_{\text{SYS-1·POP-1}} V^2_{\text{cyto}} d^2_{\text{POP-1}}}{A^2_{\text{nuc}} K^{\text{SYS-1}}_{\text{nuc:cyt}} \beta}$$

where $V_{\text{nuc}} = (V_{\text{nuc}})_{j,\infty}$ and $A_{\text{nuc}} = (A_{\text{nuc}})_{j,\infty}$.

$$a_1 = -\frac{K^{\text{POP-1}}_{\text{cyt:nuc}} K^{\text{SYS-1}}_{\text{cyt:nuc}} K^{\text{on}}_{\text{SYS-1·POP-1}} p_{\text{POP-1}}}{K^{\text{SYS-1}}_{\text{nuc:cyt}} \alpha\beta} + \frac{K^{\text{POP-1}}_{\text{cyt:nuc}} K^{\text{SYS-1}}_{\text{cyt:nuc}} K^{\text{on}}_{\text{SYS-1·POP-1}} p_{\text{SYS-1}}}{K^{\text{SYS-1}}_{\text{nuc:cyt}} \alpha\beta} + \frac{K^{\text{SYS-1·POP-1}}_{\text{cyt:nuc}} K^{\text{off}}_{\text{SYS-1·POP-1}} d_{\text{POP-1}}}{K^{\text{SYS-1·POP-1}}_{\text{nuc:cyt}} d_{\text{SYS-1·POP-1}}} + \frac{K^{\text{off}}_{\text{SYS-1·POP-1}} V_{\text{cyto}} d_{\text{POP-1}}}{V_{\text{nuc}} d_{\text{SYS-1·POP-1}}} - \frac{K^{\text{on}}_{\text{SYS-1·POP-1}} V_{\text{cyto}} p_{\text{POP-1}}}{V_{\text{nuc}}\alpha} +$$
$$\frac{K^{\text{on}}_{\text{SYS-1·POP-1}} V_{\text{cyto}} p_{\text{SYS-1}}}{V_{\text{nuc}}\alpha} + \frac{V_{\text{cyto}} d_{\text{POP-1}}}{V_{\text{nuc}}} - \frac{K^{\text{POP-1}}_{\text{cyt:nuc}} K^{\text{off}}_{\text{SYS-1·POP-1}} K^{\text{on}}_{\text{SYS-1·POP-1}} V_{\text{cyto}} p_{\text{POP-1}}}{A_{\text{nuc}} K^{\text{SYS-1}}_{\text{nuc:cyt}} \beta d_{\text{SYS-1·POP-1}}} - \frac{K^{\text{POP-1}}_{\text{cyt:nuc}} K^{\text{on}}_{\text{SYS-1·POP-1}} V_{\text{cyto}} p_{\text{POP-1}}}{A_{\text{nuc}} K^{\text{SYS-1}}_{\text{nuc:cyt}} \beta} - \frac{2 K^{\text{SYS-1}}_{\text{cyt:nuc}} K^{\text{off}}_{\text{SYS-1·POP-1}} K^{\text{on}}_{\text{SYS-1·POP-1}} V_{\text{cyto}} d_{\text{POP-1}} p_{\text{POP-1}}}{A_{\text{nuc}} K^{\text{SYS-1}}_{\text{nuc:cyt}} \alpha\beta d_{\text{SYS-1·POP-1}}} +$$
$$\frac{K^{\text{SYS-1}}_{\text{cyt:nuc}} K^{\text{off}}_{\text{SYS-1·POP-1}} K^{\text{on}}_{\text{SYS-1·POP-1}} V_{\text{cyto}} d_{\text{POP-1}} p_{\text{SYS-1}}}{A_{\text{nuc}} K^{\text{SYS-1}}_{\text{nuc:cyt}} \alpha\beta d_{\text{SYS-1·POP-1}}} + \frac{K^{\text{SYS-1}}_{\text{cyt:nuc}} {K^{\text{on}}_{\text{SYS-1·POP-1}}}^2 V_{\text{cyto}} p^2_{\text{POP-1}}}{A_{\text{nuc}} K^{\text{SYS-1}}_{\text{nuc:cyt}} \alpha^2\beta} - \frac{2 K^{\text{SYS-1}}_{\text{cyt:nuc}} {K^{\text{on}}_{\text{SYS-1·POP-1}}}^2 V_{\text{cyto}} p_{\text{POP-1}} p_{\text{SYS-1}}}{A_{\text{nuc}} K^{\text{SYS-1}}_{\text{nuc:cyt}} \alpha^2\beta} + \frac{K^{\text{SYS-1}}_{\text{cyt:nuc}} {K^{\text{on}}_{\text{SYS-1·POP-1}}}^2 V_{\text{cyto}} p^2_{\text{SYS-1}}}{A_{\text{nuc}} K^{\text{SYS-1}}_{\text{nuc:cyt}} \alpha^2\beta} +$$
$$- \frac{2 K^{\text{SYS-1}}_{\text{cyt:nuc}} K^{\text{on}}_{\text{SYS-1·POP-1}} V_{\text{cyto}} d_{\text{POP-1}} p_{\text{POP-1}}}{A_{\text{nuc}} K^{\text{SYS-1}}_{\text{nuc:cyt}} \alpha\beta} + \frac{K^{\text{SYS-1}}_{\text{cyt:nuc}} K^{\text{on}}_{\text{SYS-1·POP-1}} V_{\text{cyto}} d_{\text{POP-1}} p_{\text{SYS-1}}}{A_{\text{nuc}} K^{\text{SYS-1}}_{\text{nuc:cyt}} \alpha\beta} + \frac{{K^{\text{off}}_{\text{SYS-1·POP-1}}}^2 V_{\text{cyto}} d_{\text{POP-1}}}{A_{\text{nuc}} K^{\text{SYS-1·POP-1}}_{\text{nuc:cyt}} d_{\text{SYS-1·POP-1}}} - \frac{K^{\text{off}}_{\text{SYS-1·POP-1}} K^{\text{on}}_{\text{SYS-1·POP-1}} V_{\text{cyto}} p_{\text{POP-1}}}{A_{\text{nuc}} K^{\text{SYS-1·POP-1}}_{\text{nuc:cyt}} \alpha} +$$
$$\frac{K^{\text{off}}_{\text{SYS-1·POP-1}} K^{\text{on}}_{\text{SYS-1·POP-1}} V_{\text{cyto}} p_{\text{SYS-1}}}{A_{\text{nuc}} K^{\text{SYS-1·POP-1}}_{\text{nuc:cyt}} \alpha} + \frac{K^{\text{off}}_{\text{SYS-1·POP-1}} V_{\text{cyto}} d_{\text{POP-1}}}{A_{\text{nuc}} K^{\text{SYS-1·POP-1}}_{\text{nuc:cyt}}} - \frac{2 {K^{\text{off}}_{\text{SYS-1·POP-1}}}^2 K^{\text{on}}_{\text{SYS-1·POP-1}} V^2_{\text{cyto}} d_{\text{POP-1}} p_{\text{POP-1}}}{A^2_{\text{nuc}} K^{\text{SYS-1}}_{\text{nuc:cyt}} \beta d^2_{\text{SYS-1·POP-1}}} + \frac{2 K^{\text{off}}_{\text{SYS-1·POP-1}} {K^{\text{on}}_{\text{SYS-1·POP-1}}}^2 V^2_{\text{cyto}} p^2_{\text{POP-1}}}{A^2_{\text{nuc}} K^{\text{SYS-1}}_{\text{nuc:cyt}} \alpha\beta d_{\text{SYS-1·POP-1}}} +$$
$$- \frac{2 K^{\text{off}}_{\text{SYS-1·POP-1}} {K^{\text{on}}_{\text{SYS-1·POP-1}}}^2 V^2_{\text{cyto}} p_{\text{POP-1}} p_{\text{SYS-1}}}{A^2_{\text{nuc}} K^{\text{SYS-1}}_{\text{nuc:cyt}} \alpha\beta d_{\text{SYS-1·POP-1}}} - \frac{4 K^{\text{off}}_{\text{SYS-1·POP-1}} K^{\text{on}}_{\text{SYS-1·POP-1}} V^2_{\text{cyto}} d_{\text{POP-1}} p_{\text{POP-1}}}{A^2_{\text{nuc}} K^{\text{SYS-1}}_{\text{nuc:cyt}} \beta d_{\text{SYS-1·POP-1}}} + \frac{2 {K^{\text{on}}_{\text{SYS-1·POP-1}}}^2 V^2_{\text{cyto}} p^2_{\text{POP-1}}}{A^2_{\text{nuc}} K^{\text{SYS-1}}_{\text{nuc:cyt}} \alpha\beta} - \frac{2 {K^{\text{on}}_{\text{SYS-1·POP-1}}}^2 V^2_{\text{cyto}} p_{\text{POP-1}} p_{\text{SYS-1}}}{A^2_{\text{nuc}} K^{\text{SYS-1}}_{\text{nuc:cyt}} \alpha\beta} +$$
$$- \frac{2 K^{\text{on}}_{\text{SYS-1·POP-1}} V^2_{\text{cyto}} d_{\text{POP-1}} p_{\text{POP-1}}}{A^2_{\text{nuc}} K^{\text{SYS-1}}_{\text{nuc:cyt}} \beta}$$

$$a_0 = -\frac{K^{\text{cyt:nuc}}_{\text{SYS-1·POP-1}} K^{\text{off}}_{\text{SYS-1·POP-1}} p_{\text{POP-1}}}{K^{\text{nuc:cyt}}_{\text{SYS-1·POP-1}} d_{\text{SYS-1·POP-1}}} - \frac{K^{\text{off}}_{\text{SYS-1·POP-1}} V_{\text{cyt}} p_{\text{POP-1}}}{V_{\text{nuc}} d_{\text{SYS-1·POP-1}}} - \frac{V_{\text{cyt}} p_{\text{POP-1}}}{V_{\text{nuc}}} + \frac{K^{\text{cyt:nuc}}_{\text{SYS-1}} K^{\text{off}}_{\text{SYS-1·POP-1}} K^{\text{on}}_{\text{SYS-1·POP-1}} V_{\text{cyt}} p^2_{\text{POP-1}}}{A_{\text{nuc}} K^{\text{nuc:cyt}}_{\text{SYS-1}} \alpha\beta d_{\text{SYS-1·POP-1}}} - \frac{K^{\text{cyt:nuc}}_{\text{SYS-1}} K^{\text{off}}_{\text{SYS-1·POP-1}} K^{\text{on}}_{\text{SYS-1·POP-1}} V_{\text{cyt}} p_{\text{POP-1}} p_{\text{SYS-1}}}{A_{\text{nuc}} K^{\text{nuc:cyt}}_{\text{SYS-1}} \alpha\beta d_{\text{SYS-1·POP-1}}} +$$
$$\frac{K^{\text{cyt:nuc}}_{\text{SYS-1}} K^{\text{on}}_{\text{SYS-1·POP-1}} V_{\text{cyt}} p^2_{\text{POP-1}}}{A_{\text{nuc}} K^{\text{nuc:cyt}}_{\text{SYS-1}} \alpha\beta} - \frac{K^{\text{cyt:nuc}}_{\text{SYS-1}} K^{\text{on}}_{\text{SYS-1·POP-1}} V_{\text{cyto}} p_{\text{POP-1}} p_{\text{SYS-1}}}{A_{\text{nuc}} K^{\text{SYS-1}}_{\text{nuc:cyt}} \alpha\beta} - \frac{{K^{\text{off}}_{\text{SYS-1·POP-1}}}^2 V_{\text{cyt}} p_{\text{POP-1}}}{A_{\text{nuc}} K^{\text{SYS-1·POP-1}}_{\text{nuc:cyt}} d_{\text{SYS-1·POP-1}}} - \frac{K^{\text{off}}_{\text{SYS-1·POP-1}} V_{\text{cyt}} p_{\text{POP-1}}}{A_{\text{nuc}} K^{\text{SYS-1·POP-1}}_{\text{nuc:cyt}}} + \frac{{K^{\text{off}}_{\text{SYS-1·POP-1}}}^2 K^{\text{on}}_{\text{SYS-1·POP-1}} V^2_{\text{cyt}} p^2_{\text{POP-1}}}{A^2_{\text{nuc}} K^{\text{SYS-1}}_{\text{nuc:cyt}} \beta d^2_{\text{SYS-1·POP-1}}} +$$
$$\frac{2 K^{\text{off}}_{\text{SYS-1·POP-1}} K^{\text{on}}_{\text{SYS-1·POP-1}} V^2_{\text{cyt}} p^2_{\text{POP-1}}}{A^2_{\text{nuc}} K^{\text{SYS-1}}_{\text{nuc:cyt}} \beta d_{\text{SYS-1·POP-1}}} + \frac{K^{\text{on}}_{\text{SYS-1·POP-1}} V^2_{\text{cyt}} p^2_{\text{POP-1}}}{A^2_{\text{nuc}} K^{\text{SYS-1}}_{\text{nuc:cyt}} \beta}$$

For each value of $z$, we can compute all the other components $x := [\text{SYS-1}_{\text{cyt}}]_\infty$, $z := [\text{SYS-1·POP-1}_{\text{cyt}}]_\infty$, $a := [\text{SYS-1}_{\text{nuc}}]_\infty$, $b := [\text{POP-1}_{\text{nuc}}]_\infty$, $x := [\text{SYS-1·POP-1}_{\text{nuc}}]_\infty$, as

$$x(y) = \frac{d_{\text{POP1}} y - p_{\text{POP1}} + p_{\text{SYS1}}}{\alpha}$$

$$z(y) = \frac{-d_{\text{POP1}} y + p_{\text{POP1}}}{d_{\text{SYS-1·POP-1}}}$$

$$a(y) = \frac{1}{A_{\text{nuc}} K^{\text{SYS-1}}_{\text{nuc:cyt}} \alpha d_{\text{SYS-1·POP-1}}} \cdot (A_{\text{nuc}} K^{\text{SYS-1}}_{\text{cyt:nuc}} d_{\text{POP1}} d_{\text{SYS-1·POP-1}} y - A_{\text{nuc}} K^{\text{SYS-1}}_{\text{cyt:nuc}} d_{\text{SYS-1·POP-1}} p_{\text{POP1}} + A_{\text{nuc}} K^{\text{SYS-1}}_{\text{cyt:nuc}} d_{\text{SYS-1·POP-1}} p_{\text{SYS1}} + K^{\text{off}}_{\text{SYS-1·POP-1}} V_{\text{cyt}} \alpha d_{\text{POP1}} y$$
$$- K^{\text{off}}_{\text{SYS-1·POP-1}} V_{\text{cyt}} \alpha p_{\text{POP1}} + K^{\text{on}}_{\text{SYS-1·POP-1}} V_{\text{cyt}} d_{\text{POP1}} d_{\text{SYS-1·POP-1}} y^2 - K^{\text{on}}_{\text{SYS-1·POP-1}} V_{\text{cyt}} d_{\text{SYS-1·POP-1}} p_{\text{POP1}} y + K^{\text{on}}_{\text{SYS-1·POP-1}} V_{\text{cyt}} d_{\text{SYS-1·POP-1}} p_{\text{SYS1}} y + V_{\text{cyt}} \alpha d_{\text{POP1}} d_{\text{SYS-1·POP-1}} y - V_{\text{cyt}} \alpha d_{\text{SYS-1·POP-1}} p_{\text{POP1}})$$

$$b(y) = \frac{1}{A_{\text{nuc}}\alpha\beta d_{\text{SYS-1}\cdot\text{POP-1}}} \cdot \Big(A_{\text{nuc}}K^{\text{POP-1}}_{\text{cyt:nuc}}\alpha d_{\text{SYS-1}\cdot\text{POP-1}}y + K^{\text{off}}_{\text{SYS-1}\cdot\text{POP-1}}V_{\text{cyt}}\alpha d_{\text{POP1}}y - K^{\text{off}}_{\text{SYS-1}\cdot\text{POP-1}}V_{\text{cyt}}\alpha p_{\text{POP1}} + K^{\text{on}}_{\text{SYS-1}\cdot\text{POP-1}}V_{\text{cyt}}d_{\text{POP1}}d_{\text{SYS-1}\cdot\text{POP-1}}y^2$$
$$- K^{\text{on}}_{\text{SYS-1}\cdot\text{POP-1}}V_{\text{cyt}}d_{\text{SYS-1}\cdot\text{POP-1}}p_{\text{POP1}}y + K^{\text{on}}_{\text{SYS-1}\cdot\text{POP-1}}V_{\text{cyt}}d_{\text{SYS-1}\cdot\text{POP-1}}p_{\text{SYS1}}y + V_{\text{cyt}}\alpha d_{\text{POP1}}d_{\text{SYS-1}\cdot\text{POP-1}}y - V_{\text{cyt}}\alpha d_{\text{SYS-1}\cdot\text{POP-1}}p_{\text{POP1}}\Big)$$

$$c(y) = \frac{1}{A_{\text{nuc}}K^{\text{SYS-1}\cdot\text{POP-1}}_{\text{nuc:cyt}}\alpha d_{\text{SYS-1}\cdot\text{POP-1}}} \cdot \Big(-A_{\text{nuc}}K^{\text{SYS-1}\cdot\text{POP-1}}_{\text{cyt:nuc}}\alpha d_{\text{POP1}}y + A_{\text{nuc}}K^{\text{SYS-1}\cdot\text{POP-1}}_{\text{cyt:nuc}}\alpha p_{\text{POP1}} - K^{\text{off}}_{\text{SYS-1}\cdot\text{POP-1}}V_{\text{cyt}}\alpha d_{\text{POP1}}y + K^{\text{off}}_{\text{SYS-1}\cdot\text{POP-1}}V_{\text{cyt}}\alpha p_{\text{POP1}}$$
$$- K^{\text{on}}_{\text{SYS-1}\cdot\text{POP-1}}V_{\text{cyt}}d_{\text{POP1}}d_{\text{SYS-1}\cdot\text{POP-1}}y^2 + K^{\text{on}}_{\text{SYS-1}\cdot\text{POP-1}}V_{\text{cyt}}d_{\text{SYS-1}\cdot\text{POP-1}}p_{\text{POP1}}y - K^{\text{on}}_{\text{SYS-1}\cdot\text{POP-1}}V_{\text{cyt}}d_{\text{SYS-1}\cdot\text{POP-1}}p_{\text{SYS1}}y - V_{\text{cyt}}\alpha d_{\text{POP1}}d_{\text{SYS-1}\cdot\text{POP-1}}y + V_{\text{cyt}}\alpha d_{\text{SYS-1}\cdot\text{POP-1}}p_{\text{POP1}}\Big)$$

Thus, there are four fixed points in $\mathbb{C}$. The complex conjugate root theorem states that provided a polynomial with real coefficients, if $z$ is a root such that $\text{Im}(z) \neq 0$, then its complex conjugate $z^*$ will also be a root. Thus, without taking the constraints of the problem into account, there could be zero, two or four roots in $\mathbb{R}$. To see whether more than one fixed point in $\mathbb{R}^+$ is possible in practice, we sampled random parameters, drawn from $U(m_k, M_k)$ ($n_{\text{sampling}} = 100000$), where $m_k := 0.001$ and $M_k := 10$ for production [$p$'s] and degradation rates [$d$'s] or $m_k := 0.1$ and $M_k := 100$ for import/export, $K_{\text{on}}/K_{\text{off}}$ or Michaelis-Menten constants [$K$'s]. We then evaluated the steady-states with the analytical formula to study the nature of the different roots. The obtained counts of the randomly-sampled polynomials are shown in the table below:

| Root type | $\mathbb{C}[\Re(z) < 0] : 2;\ \mathbb{R}^- : 1;\ \mathbb{R}^+ : 1$ | $\mathbb{R}^- : 3;\ \mathbb{R}^+ : 1$ | |
|---|---|---|---|
| Number of observations | 55765 | 44235 | 100000 |

There seems to be strong statistical evidence that solely one fixed point is possible in $R^+$ in the explored neighbourhood of hyperspace. Mathematically proving this result is beyond the scope of this work.

#### IV. Model parameter fitting

##### A. Half-life of different biological subprocesses

To help us decide on appropriate values to use for each parameter, we made a set of toy models that focus on individual biological subprocesses in isolation. This way we can derive a measure of the half-life.

*1) Protein degradation:*

$$\dot{x} = p - d \cdot x$$

The analytical solution to this differential equation is:

$$x(t) = x_0 \cdot e^{-d \cdot t} + \frac{p}{d} \cdot (1 - e^{-d \cdot t})$$

$$t_{½}^{\text{degradation}} = \frac{\log(2)}{d} \tag{21}$$

*2) Nuclear/cytosolic shuttling:*

$$\begin{pmatrix} \dot{x}_{\text{cyt}} \\ \dot{x}_{\text{nuc}} \end{pmatrix} = \begin{pmatrix} -\alpha \cdot x_{\text{cyt}} + \beta \cdot x_{\text{nuc}} \\ \gamma \cdot \alpha \cdot x_{\text{cyt}} - \gamma \cdot \beta \cdot x_{\text{nuc}} \end{pmatrix}$$

where $\alpha := \frac{A_{\text{nuc}}}{V_{\text{cyt}}} \cdot K_{\text{cyt:nuc}}$, $\beta := \frac{A_{\text{nuc}}}{V_{\text{cyt}}} \cdot K_{\text{nuc:cyt}}$ and $\gamma = \frac{V_{\text{cyt}}}{V_{\text{nuc}}}$. We assume that these import/export constants have been corrected for the ratio of nuclear pore area to total nucleus area. The analytical solution to the system of equations is:

$$\begin{pmatrix} x_{\text{cyt}}(t) \\ x_{\text{nuc}}(t) \end{pmatrix} = \begin{pmatrix} x_0 \\ y_0 \end{pmatrix} \cdot e^{-(\alpha+\beta\cdot\gamma)\cdot t} + \begin{pmatrix} \frac{\gamma \cdot x_0 + y_0}{\gamma + \alpha/\beta} \\ \alpha/\beta \cdot \frac{\gamma \cdot x_0 + y_0}{\gamma + \alpha/\beta} \end{pmatrix} \cdot (1 - e^{-(\alpha+\beta\cdot\gamma)\cdot t})$$

Thus:

$$t_{½}^{\text{cyt:nuc}} = \frac{\log(2)}{\alpha + \beta \cdot \gamma} = \frac{\log(2)}{\frac{A_{\text{nuc}}}{V_{\text{cyt}}} \cdot K_{\text{cyt:nuc}} + \frac{A_{\text{nuc}}}{V_{\text{nuc}}} \cdot K_{\text{nuc:cyt}}} \tag{22}$$

*3) Complex formation:*

$$\frac{d[\text{s}]}{dt} = -K_{\text{on}} \cdot [\text{s}] \cdot [\text{p}] + K_{\text{off}} \cdot [\text{s·p}]$$
$$\frac{d[\text{p}]}{dt} = -K_{\text{on}} \cdot [\text{s}] \cdot [\text{p}] + K_{\text{off}} \cdot [\text{s·p}]$$
$$\frac{d[\text{s·p}]}{dt} = K_{\text{on}} \cdot [\text{s}] \cdot [\text{p}] - K_{\text{off}} \cdot [\text{s·p}]$$

Since $\frac{d[\text{s}]}{dt} - \frac{d[\text{p}]}{dt} = 0$, then $[\text{s}] - [\text{p}] = s_0 - p_0 := \alpha$; since $\frac{d[\text{p}]}{dt} + \frac{d[\text{s·p}]}{dt} = 0$, then $[\text{p}] + [\text{s·p}] = p_0 + c_0 := \beta$:

$$\frac{d[\text{s}\cdot\text{p}]}{dt} = K_{\text{on}}\alpha\beta + K_{\text{on}}\beta^2 - (K_{\text{off}} + K_{\text{on}}\alpha + 2K_{\text{on}}\beta - K_{\text{on}}[\text{s}\cdot\text{p}])[\text{s}\cdot\text{p}]$$

We propose the following first order approximation:

$$\frac{d[\text{s}\cdot\text{p}]}{dt} = K_{\text{on}}\alpha\beta + K_{\text{on}}\beta^2 - (K_{\text{off}} + K_{\text{on}}\alpha + 2K_{\text{on}}\beta - K_{\text{on}}c_0)[\text{s}\cdot\text{p}]$$

and thus:

$$t_{½}^{\text{on/off}} \approx \frac{\log(2)}{K_{\text{off}} + K_{\text{on}}\alpha + 2K_{\text{on}}\beta - K_{\text{on}}c_0} = \frac{\log(2)}{K_{\text{off}} + K_{\text{on}} \cdot (s_0 + p_0 + c_0)} \approx \frac{\log(2)}{K_{\text{off}} + K_{\text{on}}} \tag{23}$$

where we assume that $s_0 + p_0 + c_0$ is in the order of 1.

##### *B. Selection of parameters*

For the cumulative model, we assume that:

$$t_{½}^{\text{APC/Axin}} \approx \Delta t_{\text{division}}^{\text{M}} = 120 \text{ min} \tag{24}$$

and thus:

$$d_{\text{APC/Axin}} := \frac{\log(2)}{\Delta t_{\text{division}}} \approx 0.006 \text{ min}^{-1}$$

For the non-cumulative model, we assume that:

$$t_{½}^{\text{APC/Axin}} \approx \Delta t_{\text{division}}^{\text{embryonic}} = 15 \text{ min} \tag{25}$$

and thus:

$$d_{\text{APC/Axin}} := \frac{\log(2)}{\Delta t_{\text{division}}} \approx 0.046 \text{ min}^{-1}$$

We arbitrarily set $p_{\text{APC/Axin}} := \frac{10}{d_{\text{APC/Axin}}}$ $\mu\text{M} \cdot \text{min}^{-1}$ so that $[\text{APC/Axin}]_\infty := 10$ $\mu$M, which corresponds to ~44,000 molecules in the cytoplasmic volume of 7.3 $\mu\text{m}^3$. To maximize dose-response, we center APC/Axin regulation to its steady-state concentration by setting $K_{\text{APC/Axin:LIT-1}\cdot\text{WRM-1}} = K_{\text{APC/Axin:SYS-1}} := 10$ $\mu$M.

We also assume that $t_{½}^{\text{LIT-1}\cdot\text{WRM-1}} = t_{½}^{\text{SYS-1}} = t_{½}^{\text{POP-1}} = t_{½}^{\text{SYS-1}\cdot\text{POP-1}} \approx \frac{\Delta t_{\text{division}}^{\text{M}}}{10}$ and thus:

$$d_{\text{LIT-1}\cdot\text{WRM-1}} = d_{\text{SYS-1}} = d_{\text{POP-1}} = d_{\text{SYS-1}\cdot\text{POP-1}} := 0.06 \text{ min}^{-1}$$

We also assume that $d_{\text{APC/Axin:SYS-1}} = 10 \cdot d_{\text{SYS-1}} := 0.6 \text{ min}^{-1}$. We set $\phi := 0.8$, which is what we used in the naive model.

Nuclear/cytosol shuttling time dynamics must be set to be much faster than cell division timing. We propose the following criterion:

$$t_{½}^{\text{cyt:nuc}} := \frac{\Delta t_{\text{division}}}{100} \tag{26}$$

Since the following proteins are mostly nuclear, we also propose that:

$$\frac{K_{\text{SYS-1}}^{\text{cyt:nuc}}}{K_{\text{SYS-1}}^{\text{nuc:cyt}}} := 5$$
$$\frac{K_{\text{SYS-1·POP-1}}^{\text{cyt:nuc}}}{K_{\text{SYS-1·POP-1}}^{\text{nuc:cyt}}} := 5$$

which results in:

$$K_{\text{SYS-1}}^{\text{nuc:cyt}} = K_{\text{SYS-1·POP-1}}^{\text{nuc:cyt}} := 0.06\ \mu\text{m} \cdot \text{min}^{-1}$$
$$K_{\text{SYS-1}}^{\text{cyt:nuc}} = K_{\text{SYS-1·POP-1}}^{\text{cyt:nuc}} := 0.30\ \mu\text{m} \cdot \text{min}^{-1}$$

To fit experimental observations, a single cell division and its consequent depletion in APC/Axin should be enough to make LIT-1·WRM-1 posteriorly nuclear and anteriorly cytosolic.

After one multiplicative discontinuity, anterior and posterior cells end up with [APC/Axin] of $2 \cdot \phi \cdot [\text{APC/Axin}]_\infty = 2 \cdot \phi \cdot K_{\text{APC/Axin:LIT-1·WRM-1}}$ and $2 \cdot (1-\phi) \cdot [\text{APC/Axin}]_\infty = 2 \cdot (1-\phi) \cdot K_{\text{APC/Axin:LIT-1·WRM-1}}$, respectively. Focusing on the regulation of APC/Axin on LIT-1·WRM-1:

| [APC·Axin] | $K_{\text{APC·Axin:LIT-1·WRM-1}}$ | $2 \cdot \phi \cdot K_{\text{APC·Axin:LIT-1·WRM-1}}$ | $2 \cdot (1-\phi) \cdot K_{\text{APC·Axin:LIT-1·WRM-1}}$ |
|---|---|---|---|
| $\frac{[\text{APC·Axin}]^2}{[\text{APC·Axin}]^2 + K_{\text{APC·Axin:LIT-1·WRM-1}}^2}$ | $\frac{1}{2}$ | $\frac{4 \cdot \phi^2}{4 \cdot \phi^2 + 1}$ | $\frac{4 \cdot (1-\phi)^2}{4 \cdot (1-\phi)^2 + 1}$ |

And thus:

| [APC·Axin] | $K_{\text{APC·Axin:LIT-1·WRM-1}}$ | $2 \cdot \phi \cdot K_{\text{APC·Axin:LIT-1·WRM-1}}$ | $2 \cdot (1-\phi) \cdot K_{\text{APC·Axin:LIT-1·WRM-1}}$ |
|---|---|---|---|
| Effective cyt:nuc | $K_{\text{LIT-1·WRM-1}}^{\text{cyt:nuc}}$ | $K_{\text{LIT-1·WRM-1}}^{\text{cyt:nuc}}$ | $K_{\text{LIT-1·WRM-1}}^{\text{cyt:nuc}}$ |
| Effective nuc:cyt | $K_{\text{LIT-1·WRM-1}}^{\text{nuc:cyt}} \cdot \frac{1}{2}$ | $K_{\text{LIT-1·WRM-1}}^{\text{nuc:cyt}} \cdot \frac{4 \cdot \phi^2}{4 \cdot \phi^2 + 1}$ | $K_{\text{LIT-1·WRM-1}}^{\text{nuc:cyt}} \cdot \frac{4 \cdot (1-\phi)^2}{4 \cdot (1-\phi)^2 + 1}$ |

To set the default steady-state as equally cytosolic and nuclear:

$$\frac{K_{\text{LIT-1·WRM-1}}^{\text{cyt:nuc}}}{K_{\text{LIT-1·WRM-1}}^{\text{nuc:cyt}}} := \frac{1}{2}$$

such that:

| [APC·Axin] | $K_{\text{APC·Axin:LIT-1·WRM-1}}$ | $2 \cdot \phi \cdot K_{\text{APC·Axin:LIT-1·WRM-1}}$ | $2 \cdot (1-\phi) \cdot K_{\text{APC·Axin:LIT-1·WRM-1}}$ |
|---|---|---|---|
| $\frac{\text{Effective cyt:nuc}}{\text{Effective nuc:cyt}}$ | 1 | $\frac{1}{2} \cdot \frac{4 \cdot \phi^2 + 1}{4 \cdot \phi^2} \approx 0.70$ | $\frac{1}{2} \cdot \frac{4 \cdot (1-\phi)^2 + 1}{4 \cdot (1-\phi)^2} \approx 3.57$ |

This results in:

$$K^{\text{cyt:nuc}}_{\text{LIT-1·WRM-1}} := 0.096\ \mu\text{m} \cdot \text{min}^{-1}$$

$$K^{\text{nuc:cyt}}_{\text{LIT-1·WRM-1}} := 0.19\ \mu\text{m} \cdot \text{min}^{-1}$$

For convenience, we set $[\text{LIT-1·WRM-1}_{\text{nuc}}]_\infty := 10\ \mu\text{M}$, which means that:

$$[\text{LIT-1·WRM-1}_{\text{nuc}}]_\infty := 10 = \frac{K^{\text{cyt:nuc}}_{\text{LIT-1·WRM-1}} \cdot \frac{p_{\text{LIT-1·WRM-1}}}{d_{\text{LIT-1·WRM-1}}} \cdot \left([\text{APC·Axin}]^2_\infty + K^2_{\text{APC·Axin:LIT-1·WRM-1}}\right)}{K^{\text{nuc:cyt}}_{\text{LIT-1·WRM-1}} \cdot [\text{APC·Axin}]^2_\infty}$$

And thus $p_{\text{LIT-1·WRM-1}} = 0.6\ \mu\text{M}\cdot\text{min}^{-1}$. For maximum dose-response, we set $K_{\text{LIT-1·WRM-1:POP-1}} := 10\ \mu\text{M}$.

Focusing on the regulation of LIT-1·WRM-1 on POP-1, we can make the POP-1's default state as equally cytosolic and nuclear by setting $\frac{K^{\text{cyt:nuc}}_{\text{POP-1}}}{K^{\text{nuc:cyt}}_{\text{POP-1}}} := \frac{1}{2}$ such that:

$$\frac{K^{\text{cyt:nuc}}_{\text{POP-1}}}{K^{\text{nuc:cyt}}_{\text{POP-1}} \frac{[\text{LIT-1·WRM-1}_{\text{nuc}}]^2_\infty}{[\text{LIT-1·WRM-1}_{\text{nuc}}]^2_\infty + K^2_{\text{LIT-1·WRM-1:POP-1}}}} := 1$$

And thus:

$$K^{\text{cyt:nuc}}_{\text{POP-1}} := 0.096\ \mu\text{m} \cdot \text{min}^{-1}$$

$$K^{\text{nuc:cyt}}_{\text{POP-1}} := 0.19\ \mu\text{m} \cdot \text{min}^{-1}$$

Complex formation time dynamics must be set to be much faster than cell division timing. Assuming that complex formation is favored by a factor of 5:

$$\frac{K^{\text{on}}_{\text{SYS-1·POP-1}}}{K^{\text{off}}_{\text{SYS-1·POP-1}}} := 5$$

And that:

$$t^{\text{on/off}}_{1/2} \approx \frac{\log(2)}{K_{\text{off}} + K_{\text{on}}} := \frac{\Delta t_{\text{division}}}{500} \tag{27}$$

Then:

$$K^{\text{on}}_{\text{SYS-1}\cdot\text{POP-1}} := 2.4$$
$$K^{\text{off}}_{\text{SYS-1}\cdot\text{POP-1}} := 0.48$$

Lastly, to fit $p_{\text{SYS-1}}$ and $p_{\text{POP-1}}$, we numerically tested a wide range of combinations. Large values of one or the other, or both, completely compromised asymmetries in total POP-1 and total SYS-1 (Fig S2). We aimed to ensure that the selected values accurately captured our expectations on the total nuclear SYS-1 and POP-1 based on translational reporters, that cannot distinguish the hetero-dimer from the monomer. We selected a combination that maximized SYS-1·POP-1 asymmetry defined as:

$$\mathbb{L}(p_{\text{POP-1}}, p_{\text{SYS-1}}) = \sum_{i\in\{\text{aa,ap,pa,pp}\}} \log\left(\frac{[\text{SYS-1}\cdot\text{POP-1}]_i(t = 280 \mid p_{\text{POP-1}}, p_{\text{SYS-1}})}{[\text{SYS-1}\cdot\text{POP-1}]_{\text{aa}}(t = 280 \mid p_{\text{POP-1}}, p_{\text{SYS-1}})}\right)$$

$$\underset{p_{\text{POP-1}},p_{\text{SYS-1}}}{\arg\max}\ \mathbb{L}(p_{\text{POP-1}}, p_{\text{SYS-1}}) = \{0.046, 0.001\}$$

| Parameter | Cumulative model | Non-cumulative model | Units |
|---|---|---|---|
| $\frac{V_{\text{nuc}}}{V_{\text{cell}}}$ | 0.27 | 0.27 | |
| $\phi$ | 0.8 | 0.8 | |
| $p_{\text{APC/Axin}}$ | 0.06 | 0.46 | $\mu\text{M} \cdot \text{min}^{-1}$ |
| $p_{\text{LIT-1·WRM-1}}$ | 0.60 | 0.60 | $\mu\text{M} \cdot \text{min}^{-1}$ |
| $p_{\text{SYS-1}}$ | 0.001 | 0.001 | $\mu\text{M} \cdot \text{min}^{-1}$ |
| $p_{\text{POP-1}}$ | 0.046 | 0.046 | $\mu\text{M} \cdot \text{min}^{-1}$ |
| $d_{\text{APC/Axin}}$ | 0.006 | 0.046 | $\text{min}^{-1}$ |
| $d_{\text{LIT-1·WRM-1}}$ | 0.06 | 0.06 | $\text{min}^{-1}$ |
| $d_{\text{SYS-1}}$ | 0.06 | 0.06 | $\text{min}^{-1}$ |
| $d_{\text{POP-1}}$ | 0.06 | 0.06 | $\text{min}^{-1}$ |
| $d_{\text{SYS-1·POP-1}}$ | 0.06 | 0.06 | $\text{min}^{-1}$ |
| $d_{\text{APC/Axin:SYS-1}}$ | 0.6 | 0.6 | $\text{min}^{-1}$ |
| $K^{\text{LIT-1·WRM-1}}_{\text{nuc:cyt}}$ | 0.19 | 0.19 | $\mu\text{m} \cdot \text{min}^{-1}$ |
| $K^{\text{LIT-1·WRM-1}}_{\text{cyt:nuc}}$ | 0.096 | 0.096 | $\mu\text{m} \cdot \text{min}^{-1}$ |
| $K^{\text{SYS-1}}_{\text{nuc:cyt}}$ | 0.06 | 0.06 | $\mu\text{m} \cdot \text{min}^{-1}$ |
| $K^{\text{SYS-1}}_{\text{cyt:nuc}}$ | 0.30 | 0.30 | $\mu\text{m} \cdot \text{min}^{-1}$ |
| $K^{\text{SYS-1·POP-1}}_{\text{nuc:cyt}}$ | 0.06 | 0.06 | $\mu\text{m} \cdot \text{min}^{-1}$ |
| $K^{\text{SYS-1·POP-1}}_{\text{cyt:nuc}}$ | 0.30 | 0.30 | $\mu\text{m} \cdot \text{min}^{-1}$ |
| $K^{\text{POP-1}}_{\text{nuc:cyt}}$ | 0.19 | 0.19 | $\mu\text{m} \cdot \text{min}^{-1}$ |
| $K^{\text{POP-1}}_{\text{cyt:nuc}}$ | 0.096 | 0.096 | $\mu\text{m} \cdot \text{min}^{-1}$ |
| $K^{\text{on}}_{\text{SYS-1·POP-1}}$ | 2.4 | 2.4 | $\mu\text{M}^{-1} \cdot \text{min}^{-1}$ |
| $K^{\text{off}}_{\text{SYS-1·POP-1}}$ | 0.48 | 0.48 | $\text{min}^{-1}$ |
| $K_{\text{APC/Axin:LIT-1·WRM-1}}$ | 10 | 10 | $\mu\text{M}$ |
| $K_{\text{APC/Axin:SYS-1}}$ | 10 | 10 | $\mu\text{M}$ |
| $K_{\text{LIT-1·WRM-1:POP-1}}$ | 10 | 10 | $\mu\text{M}$ |

We set initial conditions equal to steady-states, which are the same for both the cumulative and non-cumulative models:

| Variable | Value | Units |
|---|---|---|
| $V_0^{\text{cell}}$ | 10 | $\mu$m$^3$ |
| $[\text{APC/Axin}]_0$ | 10 | $\mu$M |
| $[\text{LIT-1}\cdot\text{WRM-1}_{\text{cyt}}]_0$ | 10 | $\mu$M |
| $[\text{LIT-1}\cdot\text{WRM-1}_{\text{nuc}}]_0$ | 10.11 | $\mu$M |
| $[\text{SYS-1}_{\text{cyt}}]_0$ | 0.0018 | $\mu$M |
| $[\text{SYS-1}_{\text{nuc}}]_0$ | 0.0082 | $\mu$M |
| $[\text{POP-1}_{\text{cyt}}]_0$ | 0.761 | $\mu$M |
| $[\text{POP-1}_{\text{nuc}}]_0$ | 0.760 | $\mu$M |
| $[\text{SYS-1}\cdot\text{POP-1}_{\text{cyt}}]_0$ | 0.006 | $\mu$M |
| $[\text{SYS-1}\cdot\text{POP-1}_{\text{nuc}}]_0$ | 0.031 | $\mu$M |

#### V. WNT/$\beta$-CATENIN ASYMMETRY PATHWAY + BISTABLE MOTIFS [W$\beta$A+B]

##### A. Differential equations

$$\frac{d[\text{target}]_j}{dt} = p_{\text{target}} \cdot \frac{[\text{SYS-1·POP-1}_{\text{nuc}}]_j^2}{[\text{SYS-1·POP-1}_{\text{nuc}}]_j^2 + K^2_{\text{SYS-1·POP-1:target}}} \cdot \frac{1}{1 + [\text{POP-1}_{\text{nuc}}]_j^2 / K^2_{\text{POP-1:target}}} + p_{\text{target:target}} \cdot \frac{[\text{target}]_j^3}{[\text{target}]_j^3 + K^3_{\text{target:target}}} - d_{\text{target}} \cdot [\text{target}]_j \quad (28)$$

##### B. Steady States

Defining $p_0 := p_{\text{target}} \cdot \frac{[\text{SYS-1·POP-1}_{\text{nuc}}]_\infty^2}{[\text{SYS-1·POP-1}_{\text{nuc}}]_\infty^2 + K^2_{\text{SYS-1·POP-1:target}}} \cdot \frac{1}{1+[\text{POP-1}_{\text{nuc}}]_\infty^2 / K^2_{\text{POP-1:target}}}$ and $\mathcal{T} := [\text{target}]_\infty$, we end up with the following polynomial equation:

$$\mathcal{T}^4 + \frac{-(p_0 + p_{\text{target:target}})}{d_{\text{target}}} \cdot \mathcal{T}^3 + K^3_{\text{target:target}} \cdot \mathcal{T} - \frac{p_0 \cdot K^3_{\text{target:target}}}{d_{\text{target}}} = 0 \quad (29)$$

##### C. Parameter fitting

We set $K_{\text{SYS-1·POP-1:target}} := [\text{SYS-1·POP-1}_{\text{nuc}}]_\infty \approx 0.031\ \mu\text{M}$ and $K_{\text{POP-1:target}} := [\text{POP-1}_{\text{nuc}}]_\infty \approx 0.76\ \mu\text{M}$. As before, we set $d_{\text{target}} := \frac{\log(2)}{\Delta t^{\text{M}}_{\text{division}}/10} \approx 0.06\ \text{min}^{-1}$. The remaining parameters were fitted empirically by trial and error via numerical simulations. We chose different $K_{\text{target:target}}$ to explore different regimes. For $K_{\text{target:target}} \in \{2.5, 4\}$, $\mathcal{T}$ is monostable across cells, whilst for $K_{\text{target:target}} \in \{2.8, 3.3, 3.447876, 3.45, 3.5\}$ $\mathcal{T}$ is bistable with different cells turning from the OFF to the ON state. In this region, there are three solutions for $\mathcal{T} \in \mathbb{R}^+$: $\mathcal{T}_1 < \mathcal{T}_2 < \mathcal{T}_3$; $\mathcal{T}_2$ is unstable whilst $\mathcal{T}_1$ and $\mathcal{T}_3$ are stable fixed points.

| Parameter | Value | Units |
|---|---|---|
| $p_{\text{target}}$ | 0.1 | $\mu$M · min$^{-1}$ |
| $p_{\text{target:target}}$ | 1 | $\mu$M · min$^{-1}$ |
| $d_{\text{target}}$ | 0.06 | min$^{-1}$ |
| $K_{\text{SYS-1·POP-1:target}}$ | 0.031 | $\mu$M |
| $K_{\text{POP-1:target}}$ | 0.76 | $\mu$M |
| $K_{\text{target:target}}$ | {2.5, 2.8, 3.3, 3.447876, 3.45, 3.5, 4} | $\mu$M |

We set the initial condition of the downstream target as $\min(\mathcal{T}_1, \mathcal{T}_3)$ (initiate on the off state), except for $K_{\text{target:target}} = 2.5$ (always ON), which we initiated with the OFF state of $K_{\text{target:target}} = 3.3$:

| Variable | Value | Units |
|---|---|---|
| $[\text{target}]_0$ | {0.53, 0.53, 0.46, 0.45, 0.45, 0.45, 0.44} | $\mu$M |

### VI. Incoherence by SYS-1 and POP-1 stoichiometry-alone [W$\beta$A+B+I']

#### A. Differential equations

$$\frac{d[\mathrm{R}]_j}{dt} = \frac{p_{\mathrm{R}} \cdot [\text{SYS-1·POP-1}_{\mathrm{nuc}}]_j^2}{[\text{SYS-1·POP-1}_{\mathrm{nuc}}]_j^2 + K_{\text{SYS-1·POP-1:R}}^2} - d_{\mathrm{R}} \cdot [\mathrm{R}]_j \tag{30}$$

$$\frac{d[\mathrm{target}]_j}{dt} = \frac{p_{\mathrm{target}}}{1 + [\mathrm{R}]_j^2 / K_{\mathrm{R:target}}^2} + p_{\mathrm{target:target}} \cdot \frac{[\mathrm{target}]_j^3}{[\mathrm{target}]_j^3 + K_{\mathrm{target:target}}^3} - d_{\mathrm{target}} \cdot [\mathrm{target}]_j \tag{31}$$

#### B. Steady States

$$[\mathrm{R}]_\infty = \frac{p_{\mathrm{R}} \cdot [\text{SYS-1·POP-1}_{\mathrm{nuc}}]_\infty^2}{d_{\mathrm{R}} \cdot ([\text{SYS-1·POP-1}_{\mathrm{nuc}}]_\infty^2 + K_{\text{SYS-1·POP-1:R}}^2)} \tag{32}$$

Defining $p_0 := \frac{p_{\mathrm{target}}}{1+[\mathrm{R}]_\infty^2/K_{\mathrm{R:target}}^2}$ and $\mathcal{T} := [\mathrm{target}]_\infty$, we end up with the following polynomial equation:

$$\mathcal{T}^4 + \frac{-(p_0 + p_{\mathrm{target:target}})}{d_{\mathrm{target}}} \cdot \mathcal{T}^3 + K_{\mathrm{target:target}}^3 \cdot \mathcal{T} - \frac{p_0 \cdot K_{\mathrm{target:target}}^3}{d_{\mathrm{target}}} = 0 \tag{33}$$

#### C. Parameter fitting

We set $K_{\text{SYS-1·POP-1:R}} := [\text{SYS-1·POP-1}_{\mathrm{nuc}}]_\infty \approx 0.031\ \mu\mathrm{M}$. As before, we set $d_{\mathrm{target}} := \frac{\log(2)}{\Delta t_{\mathrm{division}}^{\mathrm{M}}/10} \approx 0.06\ \mathrm{min}^{-1}$. The remaining parameters were fitted empirically by trial and error via numerical simulations.

| Parameter | Value | Units |
|---|---|---|
| $p_{\text{POP-1}}$ | 0.001 | $\mu\mathrm{M} \cdot \mathrm{min}^{-1}$ |
| $p_{\mathrm{R}}$ | 0.6 | $\mu\mathrm{M} \cdot \mathrm{min}^{-1}$ |
| $p_{\mathrm{target}}$ | 0.1 | $\mu\mathrm{M} \cdot \mathrm{min}^{-1}$ |
| $p_{\mathrm{target:target}}$ | 1 | $\mu\mathrm{M} \cdot \mathrm{min}^{-1}$ |
| $d_{\mathrm{R}}$ | 0.06 | $\mathrm{min}^{-1}$ |
| $d_{\mathrm{target}}$ | 0.06 | $\mathrm{min}^{-1}$ |
| $K_{\text{SYS-1·POP-1:R}}$ | 0.031 | $\mu\mathrm{M}$ |
| $K_{\mathrm{R:target}}$ | 0.013 | $\mu\mathrm{M}$ |
| $K_{\mathrm{target:target}}$ | {4.36131, 4.3615, 4.362, 4.375, 4.38} | $\mu\mathrm{M}$ |

| Variable | Value | Units |
|---|---|---|
| $[\text{POP-1}_{\text{nuc}}]_0$ | 0.016 | $\mu$M |
| $[\text{POP-1}_{\text{cyt}}]_0$ | 0.016 | $\mu$M |
| $[\text{SYS-1}_{\text{nuc}}]_0$ | 0.014 | $\mu$M |
| $[\text{SYS-1}_{\text{cyt}}]_0$ | 0.0027 | $\mu$M |
| $[\text{SYS-1}\cdot\text{POP-1}_{\text{nuc}}]_0$ | 0.0011 | $\mu$M |
| $[\text{SYS-1}\cdot\text{POP-1}_{\text{cyt}}]_0$ | 0.00021 | $\mu$M |
| $[\text{target}]_0$ | $\{1.23,\ 1.23,\ 1.23,\ 1.21,\ 1.20\}$ | $\mu$M |

#### VII. WNT/$\beta$-CATENIN ASYMMETRY PATHWAY + BISTABLE MOTIFS + INCOHERENCE [W$\beta$A+B+I]

##### A. Differential Equations

$$\frac{d[\mathrm{R}]_j}{dt} = \frac{p_\mathrm{R}\cdot[\text{SYS-1}\cdot\text{POP-1}_\mathrm{nuc}]_j^2}{[\text{SYS-1}\cdot\text{POP-1}_\mathrm{nuc}]_j^2 + K^2_{\text{SYS-1}\cdot\text{POP-1:R}}}\cdot\frac{1}{1+[\text{POP-1}_\mathrm{nuc}]_j^2/K^2_{\text{POP-1:R}}} - d_\mathrm{R}\cdot[\mathrm{R}]_j \quad (34)$$

$$\frac{d[\mathrm{X}]_j}{dt} = \frac{p_\mathrm{X}\cdot[\text{SYS-1}\cdot\text{POP-1}_\mathrm{nuc}]_j^2}{[\text{SYS-1}\cdot\text{POP-1}_\mathrm{nuc}]_j^2 + K^2_{\text{SYS-1}\cdot\text{POP-1:X}}}\cdot\frac{1}{1+[\text{POP-1}_\mathrm{nuc}]_j^2/K^2_{\text{POP-1:X}}} - d_\mathrm{X}\cdot[\mathrm{X}]_j \quad (35)$$

$$\frac{d[\mathrm{target}]_j}{dt} = \frac{p_\mathrm{target}\cdot[\mathrm{X}]_j^2}{[\mathrm{X}]_j^2 + K^2_\mathrm{X:target}}\cdot\frac{1}{1+[\mathrm{R}]_j^2/K^2_\mathrm{R:target}} + p_\mathrm{target:target}\cdot\frac{[\mathrm{target}]_j^3}{[\mathrm{target}]_j^3 + K^3_\mathrm{target:target}} - d_\mathrm{target}\cdot[\mathrm{target}]_j \quad (36)$$

##### B. Steady-states

$$[\mathrm{R}]_\infty = \frac{p_\mathrm{R}\cdot[\text{SYS-1}\cdot\text{POP-1}_\mathrm{nuc}]_\infty^2}{d_\mathrm{R}\cdot[\text{SYS-1}\cdot\text{POP-1}_\mathrm{nuc}]_\infty^2 + K^2_{\text{SYS-1}\cdot\text{POP-1:R}}}\cdot\frac{1}{1+[\text{POP-1}_\mathrm{nuc}]_\infty^2/K^2_{\text{POP-1:R}}} \quad (37)$$

$$[\mathrm{X}]_\infty = \frac{p_\mathrm{X}\cdot[\text{SYS-1}\cdot\text{POP-1}_\mathrm{nuc}]_\infty^2}{d_\mathrm{X}\cdot[\text{SYS-1}\cdot\text{POP-1}_\mathrm{nuc}]_\infty^2 + K^2_{\text{SYS-1}\cdot\text{POP-1:X}}}\cdot\frac{1}{1+[\text{POP-1}_\mathrm{nuc}]_\infty^2/K^2_{\text{POP-1:X}}} \quad (38)$$

Defining $p_0 := p_\mathrm{target}\cdot\frac{[\mathrm{X}]_\infty^2}{[\mathrm{X}]_\infty^2+K^2_\mathrm{X:target}}\cdot\frac{1}{1+[\mathrm{R}]_\infty^2/K^2_\mathrm{R:target}}$ and $\mathcal{T} := [\mathrm{target}]_\infty$, we end up with the following polynomial:

$$\mathcal{T}^4 + \frac{-(p_0 + p_\mathrm{target:target})}{d_\mathrm{target}}\cdot\mathcal{T}^3 + K^3_\mathrm{target:target}\cdot\mathcal{T} - \frac{p_0\cdot K^3_\mathrm{target:target}}{d_\mathrm{target}} = 0 \quad (39)$$

In some region of parameter space, there can be three solutions for $\mathcal{T}\in\mathbb{R}^+$: $\mathcal{T}_1 < \mathcal{T}_2 < \mathcal{T}_3$; $\mathcal{T}_2$ is unstable whilst $\mathcal{T}_1$ and $\mathcal{T}_3$ are stable fixed points.

##### C. Parameter fitting

We set $K_{\text{SYS-1}\cdot\text{POP-1:R}} = K_{\text{SYS-1}\cdot\text{POP-1:X}} := [\text{SYS-1}\cdot\text{POP-1}_\mathrm{nuc}]_\infty \approx 0.031$ $\mu$M and $K_\text{POP-1:R} = K_\text{POP-1:X} := [\text{POP-1}_\mathrm{nuc}]_\infty \approx 0.76$ $\mu$M. As for model W$\beta$A+B, we set $d_\mathrm{target} := 0.06$ min$^{-1}$, $p_\mathrm{target} := 0.1$ and $p_\mathrm{target:target} := 1$. We arbitrarily set $p_\mathrm{R} = p_\mathrm{X} := 0.1$ and set $K_\mathrm{X:target} := [\mathrm{X}]_\infty$ and $K_\mathrm{R:target} := [\mathrm{R}]_\infty$. This results in a symmetric incoherence scheme. We realized that setting $d_\mathrm{X} = d_\mathrm{R} := 0.06$ min$^{-1}$ resulted in very poor asymmetries. We achieved high asymmetries by making the X branch slower by setting $d_\mathrm{X} := 0.02$ min$^{-1}$ whilst $d_\mathrm{R} := 0.06$ min$^{-1}$ (inverting this scheme results in selecting M.d(l/r)ap instead of M.v(l/r)pa and M.v(l/r)paa). Lastly, $K_\mathrm{target:target}$ was fitted empirically by trial and error via numerical simulations.

| Parameter | Value | Units |
|---|---|---|
| $p_{\text{R}}$ | 0.1 | $\mu$M · min$^{-1}$ |
| $p_{\text{X}}$ | 0.1 | $\mu$M · min$^{-1}$ |
| $p_{\text{target}}$ | 0.1 | $\mu$M · min$^{-1}$ |
| $p_{\text{target:target}}$ | 1 | $\mu$M · min$^{-1}$ |
| $d_{\text{R}}$ | 0.06 | min$^{-1}$ |
| $d_{\text{X}}$ | 0.02 | min$^{-1}$ |
| $d_{\text{target}}$ | 0.06 | min$^{-1}$ |
| $K_{\text{SYS-1·POP-1:R}}$ | 0.031 | $\mu$M |
| $K_{\text{SYS-1·POP-1:X}}$ | 0.031 | $\mu$M |
| $K_{\text{POP-1:R}}$ | 0.76 | $\mu$M |
| $K_{\text{POP-1:X}}$ | 0.76 | $\mu$M |
| $K_{\text{R:target}}$ | 0.42 | $\mu$M |
| $K_{\text{X:target}}$ | 1.25 | $\mu$M |
| $K_{\text{target:target}}$ | 2.75 | $\mu$M |

We set the initial conditions as steady-state concentrations; for the downstream target we used $\min(\mathcal{T}_1, \mathcal{T}_3)$ (initiate on the off state):

| Variable | Value | Units |
|---|---|---|
| [R]$_0$ | 0.42 | $\mu$M |
| [X]$_0$ | 1.25 | $\mu$M |
| [target]$_0$ | 0.42 | $\mu$M |

#### VIII. Wnt/$\beta$-catenin asymmetry pathway + downstream bistable motif + incoherence + memory of first cell division [W$\beta$A+B+I+M]

##### A. Differential equations

$$\frac{d[\mathrm{R}]_j}{dt} = \frac{p_\mathrm{R}\cdot[\text{SYS-1·POP-1}_\mathrm{nuc}]_j^2}{[\text{SYS-1·POP-1}_\mathrm{nuc}]_j^2 + K^2_{\text{SYS-1·POP-1:target}}}\cdot\frac{1}{1+[\text{POP-1}_\mathrm{nuc}]_j^2/K^2_{\text{POP-1:R}}} - d_\mathrm{R}\cdot[\mathrm{R}]_j \quad (40)$$

$$\frac{d[\mathrm{X}]_j}{dt} = \frac{p_\mathrm{X}\cdot[\text{SYS-1·POP-1}_\mathrm{nuc}]_j^2}{[\text{SYS-1·POP-1}_\mathrm{nuc}]_j^2 + K^2_{\text{SYS-1·POP-1:X}}}\cdot\frac{1}{1+[\text{POP-1}_\mathrm{nuc}]_j^2/K^2_{\text{POP-1:X}}} + p_\mathrm{X:X}\cdot\frac{[\mathrm{X}]_j^3}{[\mathrm{X}]_j^3+K^3_\mathrm{X:X}} - d_\mathrm{X}\cdot[\mathrm{X}]_j \quad (41)$$

$$\frac{d[\mathrm{target}]_j}{dt} = \frac{p_\mathrm{target}\cdot[\mathrm{X}]_j^2}{[\mathrm{X}]_j^2+K^2_\mathrm{X:target}}\cdot\frac{1}{1+[\mathrm{R}]_j^2/K^2_\mathrm{R:target}} + p_\mathrm{target:target}\cdot\frac{[\mathrm{target}]_j^3}{[\mathrm{target}]_j^3+K^3_\mathrm{target:target}} - d_\mathrm{target}\cdot[\mathrm{target}]_j \quad (42)$$

##### B. Steady-states

$$[\mathrm{R}]_\infty = \frac{p_\mathrm{R}\cdot[\text{SYS-1·POP-1}_\mathrm{nuc}]_\infty^2}{d_\mathrm{R}\cdot[\text{SYS-1·POP-1}_\mathrm{nuc}]_\infty^2 + K^2_{\text{SYS-1·POP-1:R}}}\cdot\frac{1}{1+[\text{POP-1}_\mathrm{nuc}]_\infty^2/K^2_{\text{POP-1:R}}} \quad (43)$$

Defining $p_0^\mathrm{X} := \frac{p_\mathrm{X}\cdot[\text{SYS-1·POP-1}_\mathrm{nuc}]_\infty^2}{[\text{SYS-1·POP-1}_\mathrm{nuc}]_\infty^2+K^2_{\text{SYS-1·POP-1:X}}}\cdot\frac{1}{1+[\text{POP-1}_\mathrm{nuc}]_\infty^2/K^2_{\text{POP-1:X}}}$ and $\mathcal{X} := [\mathrm{X}]_\infty$, we end up with the following polynomial:

$$\mathcal{X}^4 + \frac{-(p_0^\mathrm{X}+p_\mathrm{X:X})}{d_\mathrm{X}}\cdot\mathcal{X}^3 + K^3_\mathrm{X:X}\cdot\mathcal{X} - \frac{p_0^\mathrm{X}\cdot K^3_\mathrm{X:X}}{d_\mathrm{X}} = 0 \quad (44)$$

Defining $p_0^\mathrm{target} := p_\mathrm{target}\cdot\frac{[\mathrm{X}]_\infty^2}{[\mathrm{X}]_\infty^2+K^2_\mathrm{X:target}}\cdot\frac{1}{1+[\mathrm{R}]_\infty^2/K^2_\mathrm{R:target}}$ and $\mathcal{T} := [\mathrm{target}]_\infty$, we end up with the following polynomial:

$$\mathcal{T}^4 + \frac{-(p_0^\mathrm{target}+p_\mathrm{target:target})}{d_\mathrm{target}}\cdot\mathcal{T}^3 + K^3_\mathrm{target:target}\cdot\mathcal{T} - \frac{p_0^\mathrm{target}\cdot K^3_\mathrm{target:target}}{d_\mathrm{target}} = 0 \quad (45)$$

Thus, there are a total of 8 solutions in $\mathbb{C}$. In the region of interest, $\mathcal{X}$ has three solutions in $\mathbb{R}^+$ (two stable, one unstable) and for each stable solution of $\mathcal{X}$ there are three solutions in $\mathbb{R}^+$ for $\mathcal{T}$ (two stable, one unstable), then, the entire system will have a maximum of four stable fixed points in $\mathbb{R}^+$. We can enumerate them as:

$$\begin{pmatrix}\mathcal{X}\\ \mathcal{T}\end{pmatrix} \in \left\{\begin{pmatrix}\mathcal{X}_1^*\\ \mathcal{T}_{11}^*\end{pmatrix}, \begin{pmatrix}\mathcal{X}_1^*\\ \mathcal{T}_{12}^*\end{pmatrix}, \begin{pmatrix}\mathcal{X}_2^*\\ \mathcal{T}_{21}^*\end{pmatrix}, \begin{pmatrix}\mathcal{X}_2^*\\ \mathcal{T}_{22}^*\end{pmatrix}\right\} \quad (46)$$

If we assume that $\mathcal{T}_{11}^*, \mathcal{T}_{12}^*, \mathcal{T}_{21}^* \approx 0^+$ and that $\mathcal{T}_{22}^* \gg 0$, we can say that there are two effective states (target-off and target-on).

##### C. Parameter fitting

Make X remember the first cell division.

| Parameter | Value | Units |
|---|---|---|
| $p_{\text{R}}$ | 0.1 | $\mu$M $\cdot$ min$^{-1}$ |
| $p_{\text{X}}$ | 0.1 | $\mu$M $\cdot$ min$^{-1}$ |
| $p_{\text{target}}$ | 0.1 | $\mu$M $\cdot$ min$^{-1}$ |
| $p_{\text{target:target}}$ | 1 | $\mu$M $\cdot$ min$^{-1}$ |
| $p_{\text{X:X}}$ | 1 | $\mu$M $\cdot$ min$^{-1}$ |
| $d_{\text{R}}$ | 0.06 | min$^{-1}$ |
| $d_{\text{X}}$ | 0.06 | min$^{-1}$ |
| $d_{\text{target}}$ | 0.06 | min$^{-1}$ |
| $K_{\text{SYS-1}\cdot\text{POP-1:R}}$ | 0.031 | $\mu$M |
| $K_{\text{SYS-1}\cdot\text{POP-1:X}}$ | 0.031 | $\mu$M |
| $K_{\text{POP-1:R}}$ | 0.76 | $\mu$M |
| $K_{\text{POP-1:X}}$ | 0.76 | $\mu$M |
| $K_{\text{X:target}}$ | 16.96 | $\mu$M |
| $K_{\text{R:target}}$ | 0.42 | $\mu$M |
| $K_{\text{target:target}}$ | 2.7 | $\mu$M |
| $K_{\text{X:X}}$ | 3.3 | $\mu$M |

| Variable | Value | Units |
|---|---|---|
| $[\text{R}]_0$ | 0.42 | $\mu$M |
| $[\text{X}]_0$ | 0.46 | $\mu$M |
| $[\text{target}]_0$ | 0.0006 | $\mu$M |

##### D. Mutants

*1) sys-1(0):* If $p_{\text{SYS-1}} = 0$,

$$[\text{POP-1}_{\text{nuc}}]_0 = [\text{POP-1}_{\text{nuc}}]_\infty > 0$$
$$[\text{SYS-1}\cdot\text{POP-1}_{\text{nuc}}]_0 = [\text{SYS-1}\cdot\text{POP-1}_{\text{nuc}}]_\infty = 0$$
$$[\text{R}]_0 = [\text{R}]_\infty = 0$$
$$[\text{X}]_0 = [\text{X}]_\infty = 0$$
$$[\text{target}]_0 = \min([\text{target}]_\infty) = 0$$

*2) pop-1(0):* If $p_{\text{POP-1}} = 0$,

$$[\text{POP-1}_{\text{nuc}}]_0 = [\text{POP-1}_{\text{nuc}}]_\infty = 0$$
$$[\text{SYS-1}{\cdot}\text{POP-1}_{\text{nuc}}]_0 = [\text{SYS-1}{\cdot}\text{POP-1}_{\text{nuc}}]_\infty = 0$$
$$[\text{R}]_0 = [\text{R}]_\infty = 0$$
$$[\text{X}]_0 = [\text{X}]_\infty = 0$$
$$[\text{target}]_0 = \min([\text{target}]_\infty) = 0$$

### IX. TEMPORAL SYMMETRY BREAK MODEL [TSB]

#### A. Differential equations

$$\frac{d[\mathrm{P}]_j}{dt} = p_P \cdot \delta(t - \tau_1) - d_\mathrm{P} \cdot [\mathrm{P}]_j \quad (47)$$

$$\frac{d[\mathrm{R}]_j}{dt} = p_0^\mathrm{R} + \frac{p_\mathrm{R} \cdot [\text{SYS-1}\cdot\text{POP-1}_\mathrm{nuc}]_j^2}{[\text{SYS-1}\cdot\text{POP-1}_\mathrm{nuc}]_j^2 + K^2_{\text{SYS-1}\cdot\text{POP-1:target}}} - d_\mathrm{R} \cdot [\mathrm{R}]_j \quad (48)$$

$$\frac{d[\mathrm{K}]_j}{dt} = \frac{p_\mathrm{K}}{1 + [\text{POP-1}_\mathrm{nuc}]_j^2 / K^2_{\text{POP-1:K}}} - d_\mathrm{K} \cdot [\mathrm{K}]_j \quad (49)$$

$$\frac{d[\mathrm{X}]_j}{dt} = p_0^\mathrm{X} + p_\mathrm{X} \cdot [\mathrm{P}]_j \cdot \frac{1}{1 + [\text{POP-1}_\mathrm{nuc}]_j^2 / K^2_{\text{POP-1:X}}} \cdot \frac{1}{1 + [\mathrm{K}]_j^2 / K_\mathrm{K:X}} + \frac{p_\mathrm{X:X} \cdot [\mathrm{X}]_j^3}{[\mathrm{X}]_j^3 + K^3_\mathrm{X:X}} - d_\mathrm{X} \cdot [\mathrm{X}]_j \quad (50)$$

$$\frac{d[\mathrm{target}]_j}{dt} = \frac{p_\mathrm{target} \cdot [\mathrm{X}]_j^2}{[\mathrm{X}]_j^2 + K^2_\mathrm{X:target}} \cdot \frac{1}{1 + [\mathrm{R}]_j^2 / K^2_\mathrm{R:target}} + p_\mathrm{target:target} \cdot \frac{[\mathrm{target}]_j^3}{[\mathrm{target}]_j^3 + K^3_\mathrm{target:target}} - d_\mathrm{target} \cdot [\mathrm{target}]_j \quad (51)$$

#### B. Steady-states

$$[\mathrm{P}]_\infty = 0 \quad (52)$$

$$[\mathrm{R}]_\infty = \frac{p_\mathrm{R} \cdot [\text{SYS-1}\cdot\text{POP-1}_\mathrm{nuc}]_\infty^2}{d_\mathrm{R} \cdot ([\text{SYS-1}\cdot\text{POP-1}_\mathrm{nuc}]_\infty^2 + K^2_{\text{SYS-1}\cdot\text{POP-1:R}})} \quad (53)$$

$$[\mathrm{K}]_\infty = \frac{p_\mathrm{K}}{d_\mathrm{K} \cdot (1 + [\text{POP-1}_\mathrm{nuc}]_\infty^2 / K^2_{\text{POP-1:K}})} \quad (54)$$

The steady states for X, $\mathcal{X}$, are the solutions to the following polynomial equation:

$$\mathcal{X}^4 + \frac{-(p_0^\mathrm{X} + p_\mathrm{X:X})}{d_\mathrm{X}} \cdot \mathcal{X}^3 + K^3_\mathrm{X:X} \cdot \mathcal{X} - \frac{p_0^\mathrm{X} \cdot K^3_\mathrm{X:X}}{d_\mathrm{X}} = 0 \quad (55)$$

Defining $p_0^\mathrm{target} := p_\mathrm{target} \cdot \frac{[\mathrm{X}]_\infty^2}{[\mathrm{X}]_\infty^2 + K^2_\mathrm{X:target}} \cdot \frac{1}{1+[\mathrm{R}]_\infty^2 / K^2_\mathrm{R:target}}$ and $\mathcal{T} := [\mathrm{target}]_\infty$, we end up with the following polynomial:

$$\mathcal{T}^4 + \frac{-(p_0^\mathrm{target} + p_\mathrm{target:target})}{d_\mathrm{target}} \cdot \mathcal{T}^3 + K^3_\mathrm{target:target} \cdot \mathcal{T} - \frac{p_0^\mathrm{target} \cdot K^3_\mathrm{target:target}}{d_\mathrm{target}} = 0 \quad (56)$$

#### C. Parameter fitting

We set $K_\text{POP-1:K} = \frac{[\text{POP-1}_\mathrm{nuc}]_\infty}{25}$, so that the dynamics of K are not relevant for wildtype steady-state concentrations of POP-1; the killswitch can only kick-in when POP-1 concentrations in the order of $\frac{[\text{POP-1}_\mathrm{nuc}]_\infty}{25}$. We kept all parameters with the same value as before except for $K_\mathrm{X:target}$, $K_\mathrm{target:target}$ and $K_\mathrm{X:X}$, we you fitted numerically via trial and error:

| Parameter | Value | Units |
|---|---|---|
| $p_{\mathrm{P}}$ | 1 | $\mu\mathrm{M}\cdot\mathrm{min}^{-1}$ |
| $p_{\mathrm{K}}$ | 1 or 0 | $\mu\mathrm{M}\cdot\mathrm{min}^{-1}$ |
| $p_{\mathrm{R}}$ | 0.1 | $\mu\mathrm{M}\cdot\mathrm{min}^{-1}$ |
| $p_{\mathrm{X}}$ | 0.1 | $\mu\mathrm{M}\cdot\mathrm{min}^{-1}$ |
| $p_0^{\mathrm{X}}$ | 0.01 | $\mu\mathrm{M}\cdot\mathrm{min}^{-1}$ |
| $p_0^{\mathrm{R}}$ | 0.01 | $\mu\mathrm{M}\cdot\mathrm{min}^{-1}$ |
| $p_{\mathrm{target}}$ | 0.1 | $\mu\mathrm{M}\cdot\mathrm{min}^{-1}$ |
| $p_{\mathrm{target:target}}$ | 1 | $\mu\mathrm{M}\cdot\mathrm{min}^{-1}$ |
| $p_{\mathrm{X:X}}$ | 1 | $\mu\mathrm{M}\cdot\mathrm{min}^{-1}$ |
| $d_{\mathrm{R}}$ | 0.06 | $\mathrm{min}^{-1}$ |
| $d_{\mathrm{X}}$ | 0.06 | $\mathrm{min}^{-1}$ |
| $d_{\mathrm{target}}$ | 0.06 | $\mathrm{min}^{-1}$ |
| $d_{\mathrm{P}}$ | 0.06 | $\mathrm{min}^{-1}$ |
| $d_{\mathrm{K}}$ | 0.06 | $\mathrm{min}^{-1}$ |
| $K_{\mathrm{SYS\text{-}1\cdot POP\text{-}1:R}}$ | 0.031 | $\mu\mathrm{M}$ |
| $K_{\mathrm{POP\text{-}1:X}}$ | 0.76 | $\mu\mathrm{M}$ |
| $K_{\mathrm{X:target}}$ | 3 | $\mu\mathrm{M}$ |
| $K_{\mathrm{R:target}}$ | 0.42 | $\mu\mathrm{M}$ |
| $K_{\mathrm{target:target}}$ | 1.875 | $\mu\mathrm{M}$ |
| $K_{\mathrm{X:X}}$ | 2.3 | $\mu\mathrm{M}$ |
| $K_{\mathrm{POP\text{-}1:K}}$ | 0.03 | $\mu\mathrm{M}$ |
| $K_{\mathrm{K:X}}$ | 0.1 | $\mu\mathrm{M}$ |

For models with ($p_K = 1$) or without kill-switch ($p_K = 0$):

| Variable | Value ($p_K = 1$) | Value ($p_K = 0$) | Units |
|---|---|---|---|
| $[\mathrm{R}]_0$ | 0.998 | 0.998 | $\mu\mathrm{M}$ |
| $[\mathrm{X}]_0$ | 0.174 | 0.174 | $\mu\mathrm{M}$ |
| $[\mathrm{K}]_0$ | 0.026 | 0 | $\mu\mathrm{M}$ |
| $[\mathrm{target}]_0$ | 0.001 | 0.001 | $\mu\mathrm{M}$ |

#### X. SPATIAL SYMMETRY BREAK MODEL [SSB]

##### A. Assumptions

- Morphogen is present in a increasing gradient along the anterior-posterior axis ($x$)
- Either: gradient vanishes at $t \to \infty$ (if $D > 0$) or gradient in the shape of a step function as in a French flag model (if $D = 0$)
- Cells are point-like towards sensing the morphogen
- The gradient and the cells are uncoupled (cells do not affect the gradient).
- Cellular positions are fixed in time for each cellular identity, though cell divisions alter their position along the gradient; e.g. $x_{\text{M.d(l/r)a}} < x_{\text{M.d(l/r)}} < x_{\text{M.d(l/r)p}}$
- Perfect dorso-ventral alignment; e.g. $x_{\text{M.d(l/r)pa}} = x_{\text{M.v(l/r)pa}}$
- Cells in the M lineage remain in contact after cell divisions.

##### B. Differential equations

We envision a morphogen, $M$, diffusing in one dimension (corresponding to the anterior-posterior axis), whose dynamics in time evolve following the partial differential equation:

$$\frac{\partial M(x,t)}{\partial t} = D \cdot \frac{\partial^2 M(x,t)}{\partial x^2}; \quad M(x,0) = H(x) := \frac{1}{2} + \frac{1}{2} \cdot \text{sign}(x) \tag{57}$$

where $H(x)$ is the Heaviside step function, $\text{sign}(x)$ the sign function, and we assume that the morphogen concentration has been normalized by maximum concentration, $M(x \to +\infty, t)$, and thus, $M(x,t) \in [0,1]$ and is adimensional. This equation can be solved analytically as:

$$M(x,t) = \frac{1}{2} + \frac{1}{2} \cdot \text{erf}\left(\frac{x}{\sqrt{4 \cdot D \cdot t}}\right) \tag{58}$$

where $\text{erf}(x)$ is the error function. We set the maximum asymmetry at the time of the first anterior-posterior cell division, $\tau_1$, by introducing a time delay into the prior expression as:

$$M(x,t|\tau_1) = \begin{cases} t \leq \tau_1 & H(x) \\ t > \tau_1 & \frac{1}{2} + \frac{1}{2} \cdot \text{erf}\left(\frac{x}{\sqrt{4 \cdot D \cdot (t-\tau_1)}}\right) \end{cases} \tag{59}$$

Every division reduces cellular radius by a factor of $\sqrt[3]{2}$:

$$R(k) = R_0 \cdot \left(\frac{1}{\sqrt[3]{2}}\right)^K \tag{60}$$

We can assign each cell across the M lineage to a corresponding x coordinates as:

| Cell | $x_j$ | Value | Units |
|---|---|---|---|
| M.(d/v)(l/r) | $0$ | 0 | $\mu$m |
| M.(d/v)(l/r)a | $-R(1)$ | -1.06 | $\mu$m |
| M.(d/v)(l/r)p | $R(1)$ | 1.06 | $\mu$m |
| M.(d/v)(l/r)aa | $-3 \cdot R(2)$ | -2.53 | $\mu$m |
| M.(d/v)(l/r)ap | $-R(2)$ | -0.84 | $\mu$m |
| M.(d/v)(l/r)pa | $R(2)$ | 0.84 | $\mu$m |
| M.(d/v)(l/r)pp | $3 \cdot R(2)$ $[g(t) = 2]$ or $4 \cdot R(3) + R(2)$ $[g(t) = 3]$ | 2.53 or 3.51 | $\mu$m |
| M.(d/v)(l/r)paa | $R(3)$ | 0.67 | $\mu$m |
| M.(d/v)(l/r)pap | $3 \cdot R(3)$ | 2.00 | $\mu$m |

where $g \in \{0, 1, 2, 3\}$ is the number of anterior-posterior cell divisions defined as:

$$g(t) := \sum_{i=1}^{N_{\text{division}}} \mathbb{1}_{\{t>\tau_i\}} \tag{61}$$

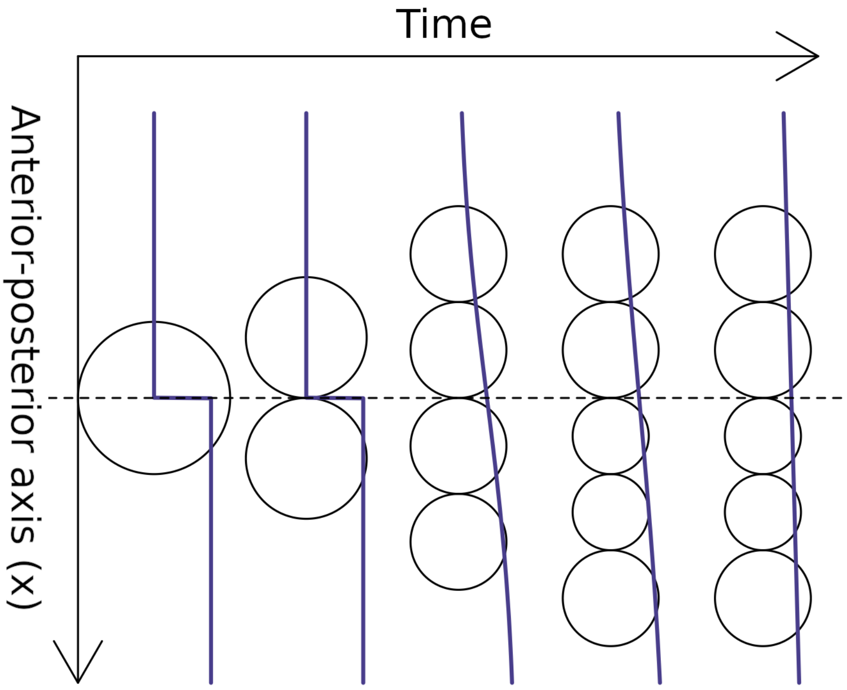


**Fig. SM2:** Visual representation of the set-up

We propose the following model:

$$\frac{d[\mathrm{R}]_j}{dt} = p_0^{\mathrm{R}} + \frac{p_{\mathrm{R}} \cdot [\text{SYS-1}\cdot\text{POP-1}_{\mathrm{nuc}}]_j^2}{[\text{SYS-1}\cdot\text{POP-1}_{\mathrm{nuc}}]_j^2 + K^2_{\text{SYS-1}\cdot\text{POP-1:target}}} \cdot \frac{1}{1 + [\text{POP-1}_{\mathrm{nuc}}]_j^2 / K^2_{\text{POP-1:R}}} - d_{\mathrm{R}} \cdot [\mathrm{R}]_j \quad (62)$$

$$\frac{d[\mathrm{X}]_j}{dt} = p_{\mathrm{X}} \cdot M(x, t|\tau_1) + p_{\mathrm{X:X}} \cdot \frac{[\mathrm{X}]_j^3}{[\mathrm{X}]_j^3 + K^3_{\mathrm{X:X}}} - d_{\mathrm{X}} \cdot [\mathrm{X}]_j \quad (63)$$

$$\frac{d[\text{target}]_j}{dt} = \frac{p_{\text{target}} \cdot [\mathrm{X}]_j^2}{[\mathrm{X}]_j^2 + K^2_{\text{X:target}}} \cdot \frac{1}{1 + [\mathrm{R}]_j^2 / K^2_{\text{R:target}}} + p_{\text{target:target}} \cdot \frac{[\text{target}]_j^3}{[\text{target}]_j^3 + K^3_{\text{target:target}}} - d_{\text{target}} \cdot [\text{target}]_j \quad (64)$$

We add background expression of the repressor R, $p_0^{\mathrm{R}} > 0$, to ensure that $[\mathrm{R}]_\infty > 0$ even if $[\text{SYS-1}\cdot\text{POP-1}_{\mathrm{nuc}}]_\infty = 0$.

#### *C. Steady-states*

$$[\mathrm{R}]_\infty = \frac{p_{\mathrm{R}} \cdot [\text{SYS-1}\cdot\text{POP-1}_{\mathrm{nuc}}]_\infty^2}{d_{\mathrm{R}} \cdot [\text{SYS-1}\cdot\text{POP-1}_{\mathrm{nuc}}]_\infty^2 + K^2_{\text{SYS-1}\cdot\text{POP-1:R}}} \cdot \frac{1}{1 + [\text{POP-1}_{\mathrm{nuc}}]_\infty^2 / K^2_{\text{POP-1:R}}} \quad (65)$$

Defining $p_0^{\mathrm{X}} := \frac{p_{\mathrm{X}}}{2}$ and $\mathcal{X} := [\mathrm{X}]_\infty$, we end up with the following polynomial:

$$\mathcal{X}^4 + \frac{-(p_0^{\mathrm{X}} + p_{\mathrm{X:X}})}{d_{\mathrm{X}}} \cdot \mathcal{X}^3 + K^3_{\mathrm{X:X}} \cdot \mathcal{X} - \frac{p_0^{\mathrm{X}} \cdot K^3_{\mathrm{X:X}}}{d_{\mathrm{X}}} = 0 \quad (66)$$

Defining $p_0^{\text{target}} := p_{\text{target}} \cdot \frac{[\mathrm{X}]_\infty^2}{[\mathrm{X}]_\infty^2 + K^2_{\text{X:target}}} \cdot \frac{1}{1+[\mathrm{R}]_\infty^2/K^2_{\text{R:target}}}$ and $\mathcal{T} := [\text{target}]_\infty$, we end up with the following polynomial:

$$\mathcal{T}^4 + \frac{-(p_0^{\text{target}} + p_{\text{target:target}})}{d_{\text{target}}} \cdot \mathcal{T}^3 + K^3_{\text{target:target}} \cdot \mathcal{T} - \frac{p_0^{\text{target}} \cdot K^3_{\text{target:target}}}{d_{\text{target}}} = 0 \quad (67)$$

Thus, there are a total of 8 solutions in $\mathbb{C}$. In the region of interest, $\mathcal{X}$ has three solutions in $\mathbb{R}^+$ (two stable, one unstable) and for each stable solution of $\mathcal{X}$ there are three solutions in $\mathbb{R}^+$ for $\mathcal{T}$ (two stable, one unstable), then, the entire system will have a maximum of four stable fixed points in $\mathbb{R}^+$. We can enumerate them as:

$$\begin{pmatrix} \mathcal{X} \\ \mathcal{T} \end{pmatrix} \in \left\{ \begin{pmatrix} \mathcal{X}_1^* \\ \mathcal{T}_{11}^* \end{pmatrix}, \begin{pmatrix} \mathcal{X}_1^* \\ \mathcal{T}_{12}^* \end{pmatrix}, \begin{pmatrix} \mathcal{X}_2^* \\ \mathcal{T}_{21}^* \end{pmatrix}, \begin{pmatrix} \mathcal{X}_2^* \\ \mathcal{T}_{22}^* \end{pmatrix} \right\} \quad (68)$$

If we assume that $\mathcal{T}_{11}^*, \mathcal{T}_{12}^*, \mathcal{T}_{21}^* \approx 0^+$ and that $\mathcal{T}_{22}^* \gg 0$, we can say that there are two effective states (target-off and target-on).

#### *D. Parameter fitting*

We can compute the halflife of the dissipation of the gradient between positions $-x$ and $x$ as:

$$\Delta M(x, t_{½}) = M(x, t_{½}) - M(-x, t_{½}) = \operatorname{erf}\left(\frac{x}{\sqrt{4 \cdot D \cdot t_{½}}}\right) = \frac{1}{2} \quad (69)$$

and thus:

$$t_{½}(x) = \frac{1}{4 \cdot D} \cdot \left( \frac{x}{\mathrm{erf}^{-1}(1/2)} \right)^2 \tag{70}$$

We propose the following criterion for $D$: $x := 5 \cdot R_M$, $t_{½} := \Delta t^{\mathrm{M}}_{\mathrm{division}}$ and thus

$$D := \frac{1}{4 \cdot \Delta t^{\mathrm{M}}_{\mathrm{division}}} \cdot \left( \frac{5 \cdot R_M}{\mathrm{erf}^{-1}(1/2)} \right)^2 \approx 1.03\ \mu\mathrm{m}^2 \cdot \mathrm{min}^{-1} = 0.017\ \mu\mathrm{m}^2 \cdot \mathrm{s}^{-1} \tag{71}$$

in the same order of magnitude as mKikGR-Wnt8 in *Xenopus* ($D_a \approx 0.042\ \mu\mathrm{m}^2 \cdot \mathrm{s}^{-1}$ [4]). To make the gradient permanent and step-like, we simply set $D := 0$.

| Parameter | Value | Units |
|---|---|---|
| $p_{\mathrm{R}}$ | 0.1 | $\mu$M $\cdot$ min$^{-1}$ |
| $p_0^{\mathrm{R}}$ | 0.01 | $\mu$M $\cdot$ min$^{-1}$ |
| $p_{\mathrm{X}}$ | 0.05 | $\mu$M $\cdot$ min$^{-1}$ |
| $p_{\mathrm{X:X}}$ | 1 | $\mu$M $\cdot$ min$^{-1}$ |
| $p_{\mathrm{target}}$ | 0.1 | $\mu$M $\cdot$ min$^{-1}$ |
| $p_{\mathrm{target:target}}$ | 1 | $\mu$M $\cdot$ min$^{-1}$ |
| $d_{\mathrm{R}}$ | 0.06 | min$^{-1}$ |
| $d_{\mathrm{X}}$ | 0.06 | min$^{-1}$ |
| $d_{\mathrm{target}}$ | 0.06 | min$^{-1}$ |
| $K_{\mathrm{SYS\text{-}1 \cdot POP\text{-}1:R}}$ | 0.031 | $\mu$M |
| $K_{\mathrm{POP\text{-}1:R}}$ | 0.76 | $\mu$M |
| $K_{\mathrm{X:target}}$ | 17.02 | $\mu$M |
| $K_{\mathrm{R:target}}$ | 0.42 | $\mu$M |
| $K_{\mathrm{target:target}}$ | 2.1 | $\mu$M |
| $K_{\mathrm{X:X}}$ | 2.7 | $\mu$M |
| $D$ | 1.03 (transient gradient) or 0 (permanent gradient) | $\mu$m$^2$ $\cdot$ min$^{-1}$ |

| Variable | Value | Units |
|---|---|---|
| $[\mathrm{R}]_0$ | 0.582 | $\mu$M |
| $[\mathrm{X}]_0$ | 0.582 | $\mu$M |
| $[\mathrm{target}]_0$ | 0.001 | $\mu$M |
