## Supplementary Figures for "Quantitative modelling of fate specification in the *C. elegans* postembryonic M lineage reveals a missing spatiotemporal signal"

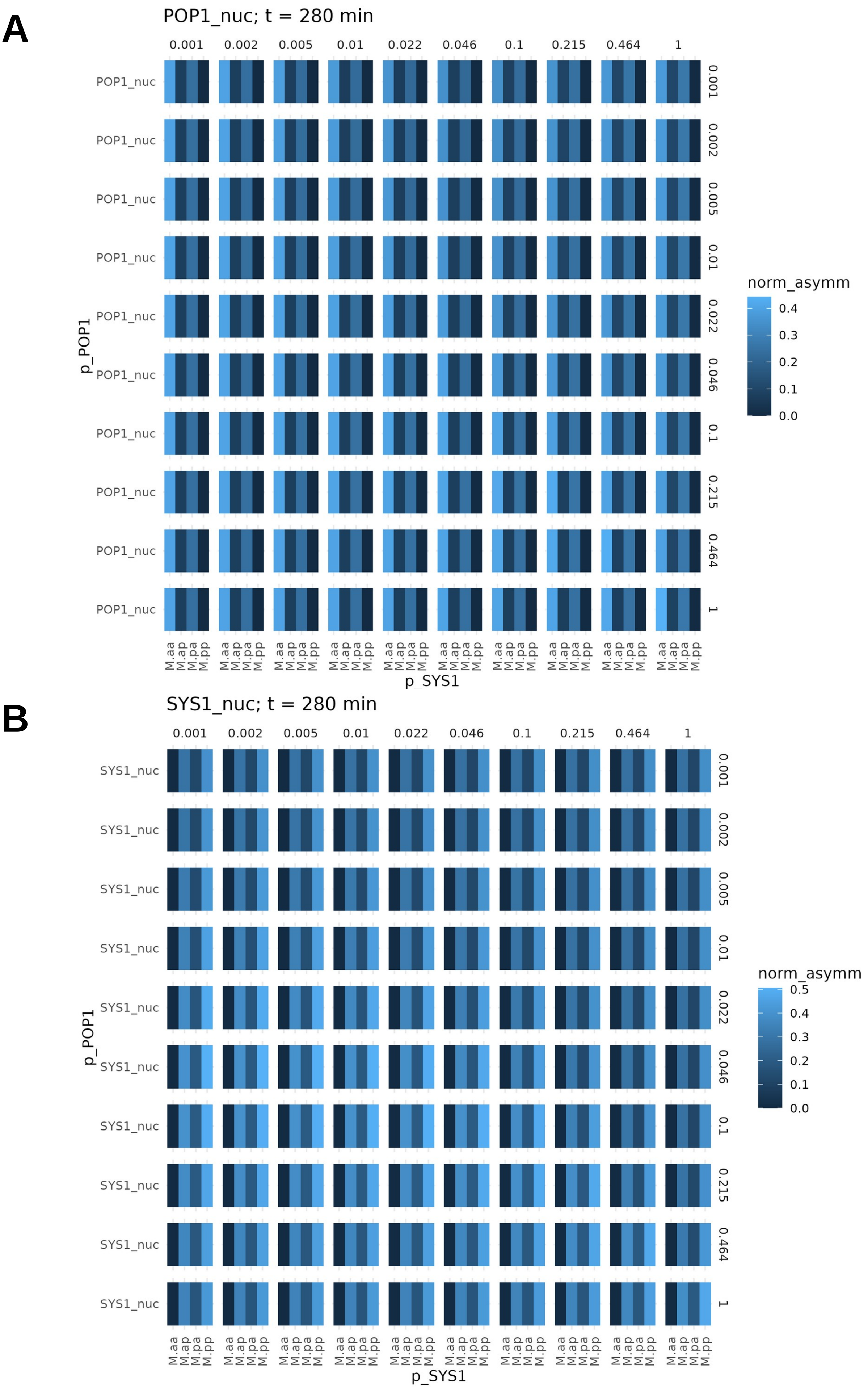


**Figure S1**. Relative asymmetries at t = 280 min as a function of $p_{POP1}$ and $p_{SYS1}$. (A) Asymmetry in nuclear POP-1 (relative to pp). (B) Asymmetry in nuclear SYS-1 (relative to aa).

**A**

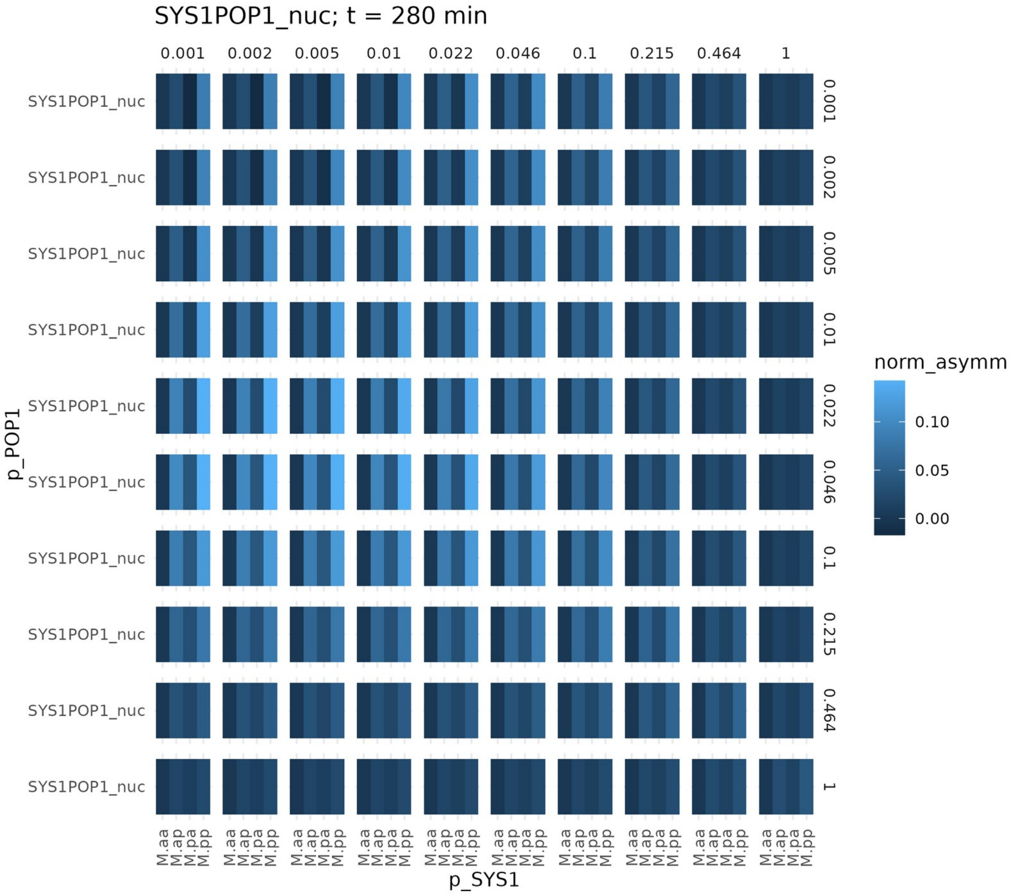


**B**

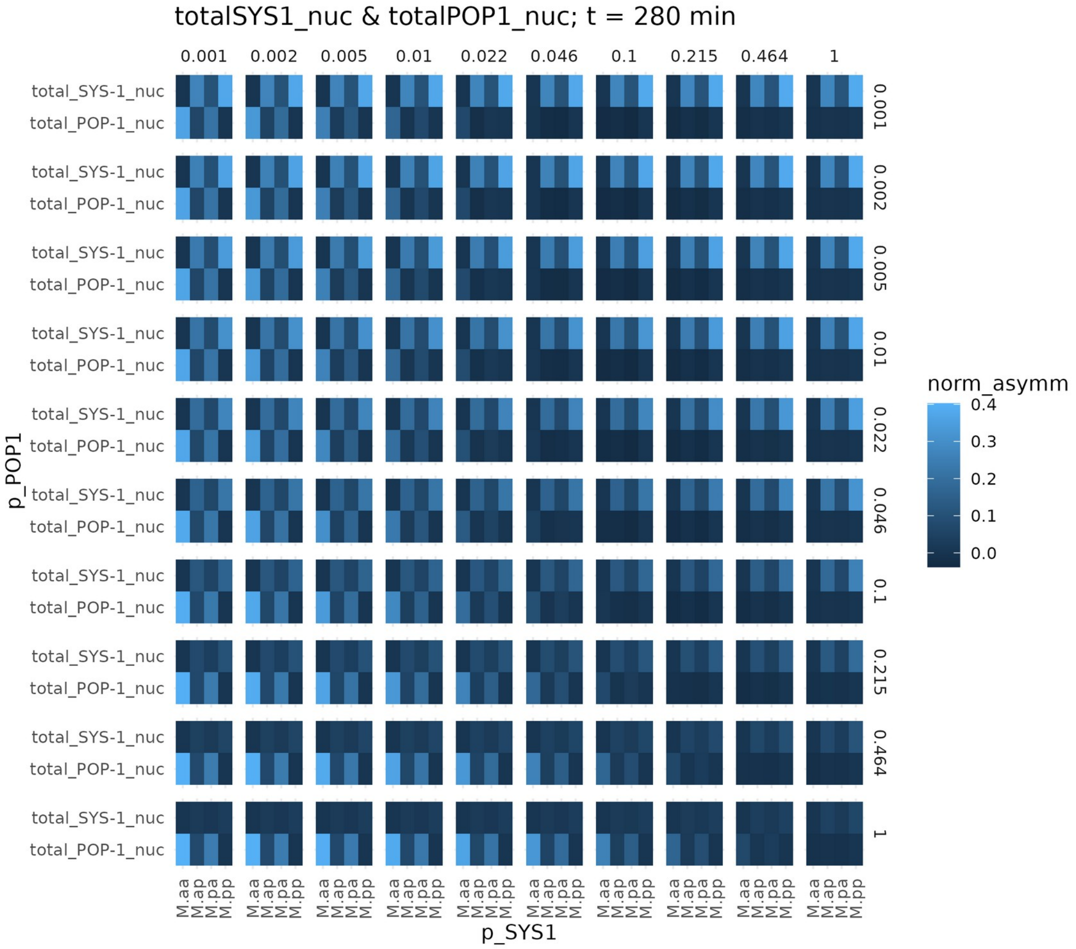


**Figure S2**. Relative asymmetries at t = 280 min as a function of $p_{POP1}$ and $p_{SYS1}$. (A) Asymmetry in nuclear SYS-1·POP-1 (relative to aa). (B) Asymmetry in total nuclear SYS-1 (relative to aa) and total POP-1 (relative to pp).

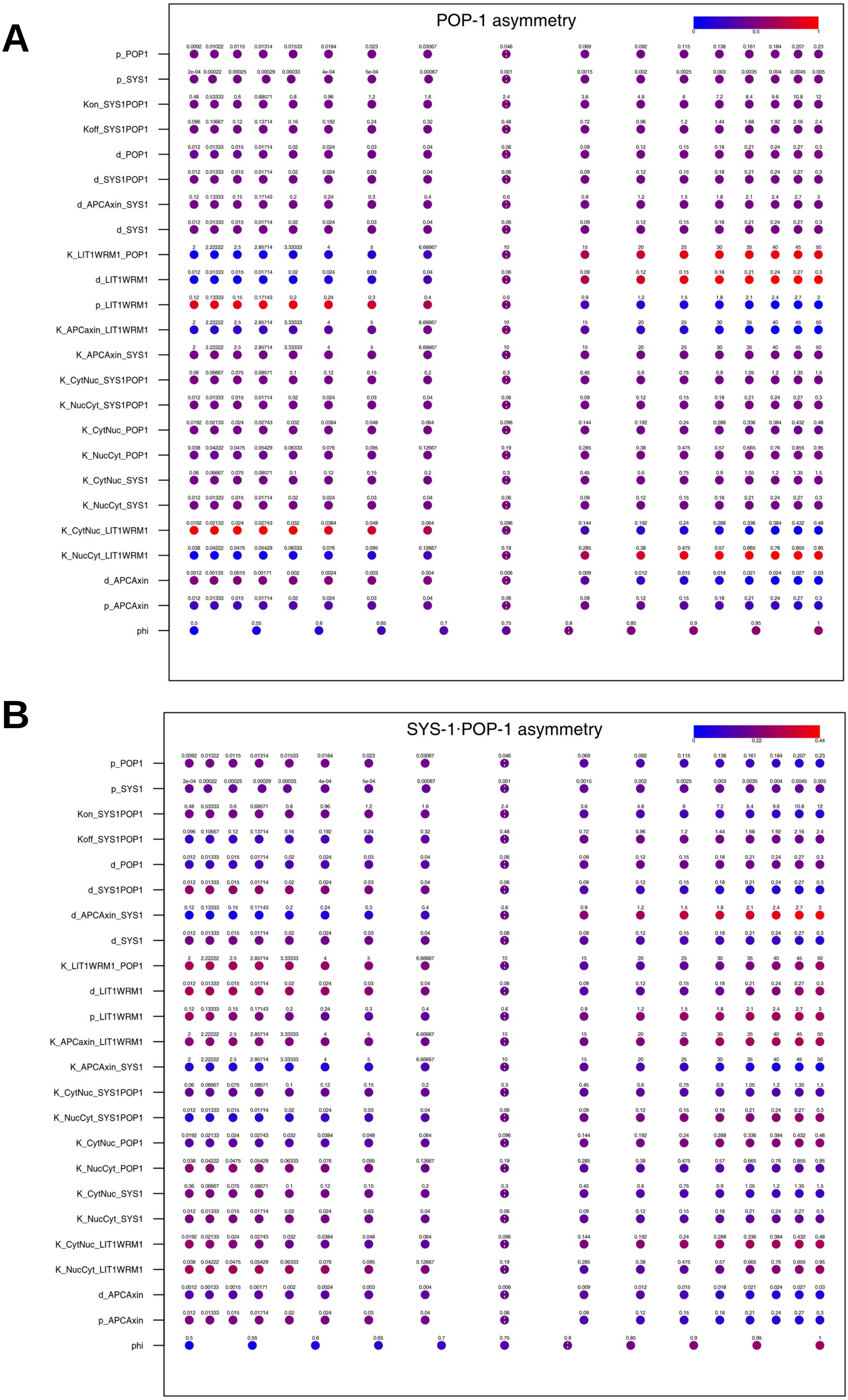


**Figure S3**. Sensitivities of different parameters on the (A) maximal relative asymmetry in nuclear POP-1 and (B) maximal relative asymmetry in nuclear SYS-1·POP-1. Each row represents a different model parameter. Selected reference parameters are shown with a dotted yellow line

A

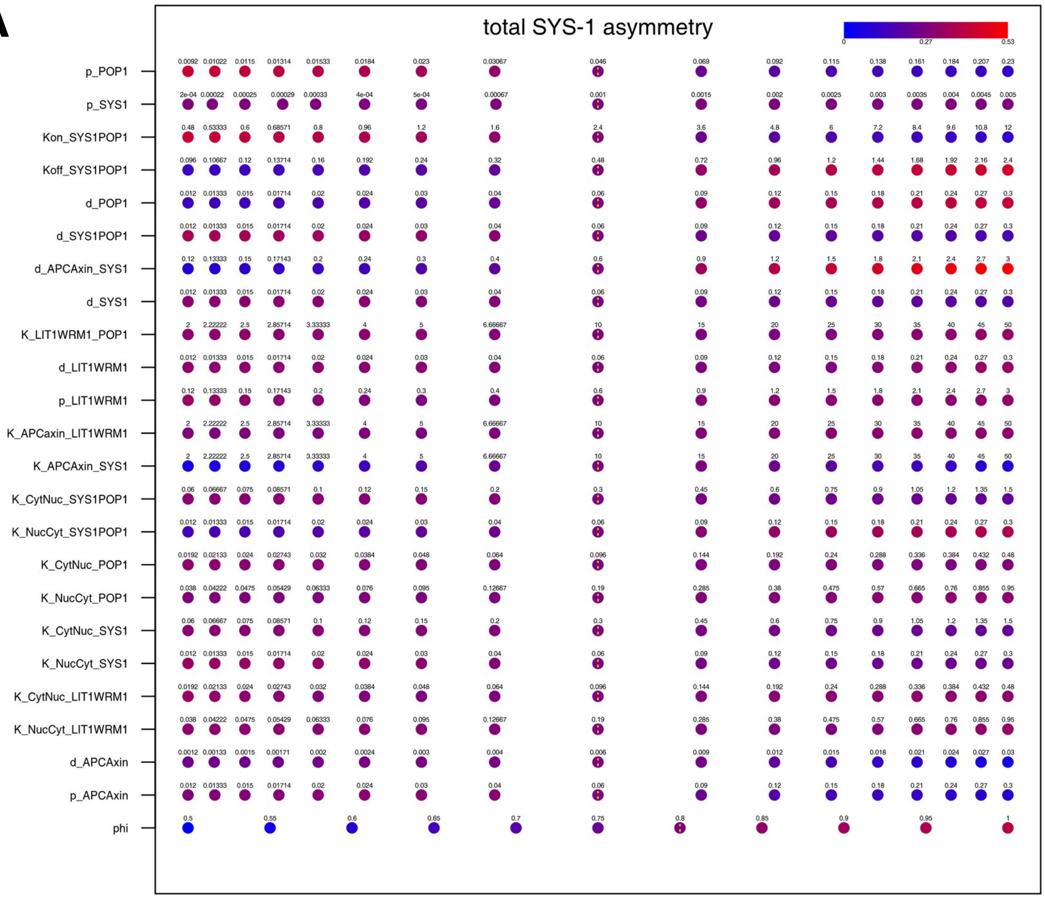


B

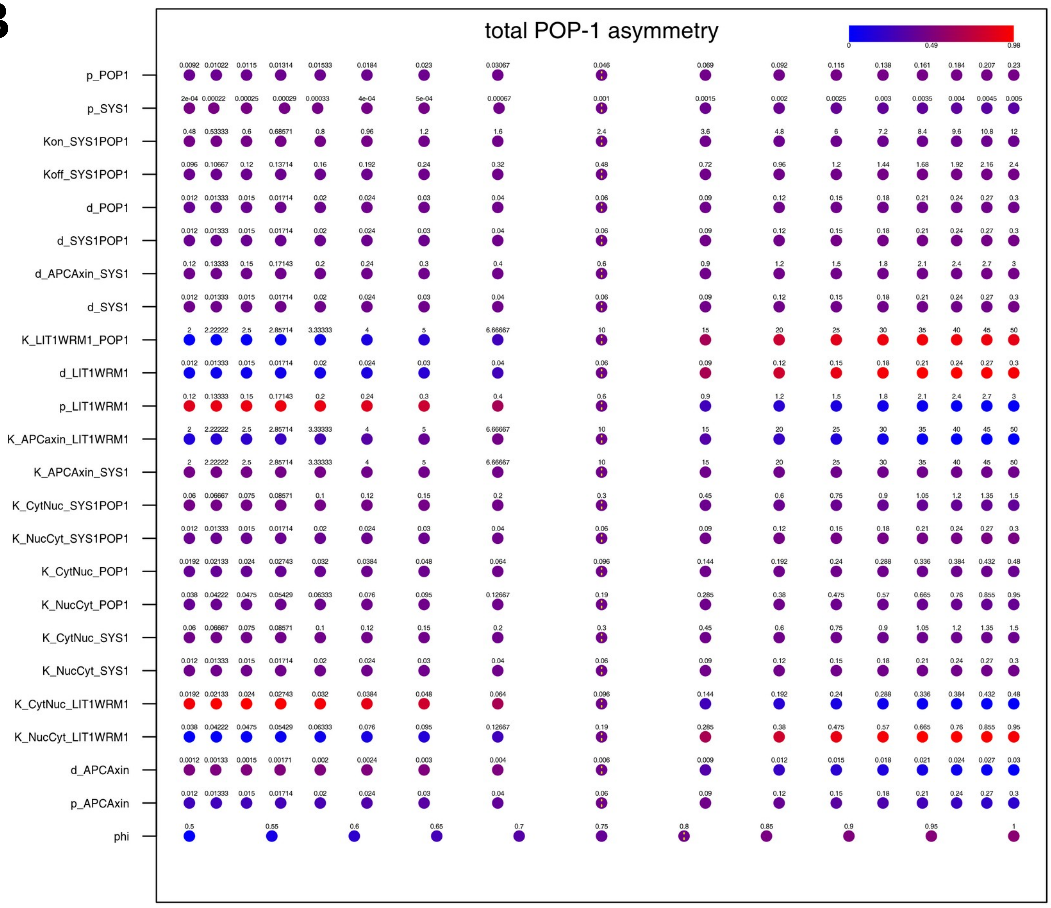


**Figure S4**. Sensitivities of different parameters on the (A) maximal relative asymmetry in total nuclear SYS-1 and (B) total POP-1.

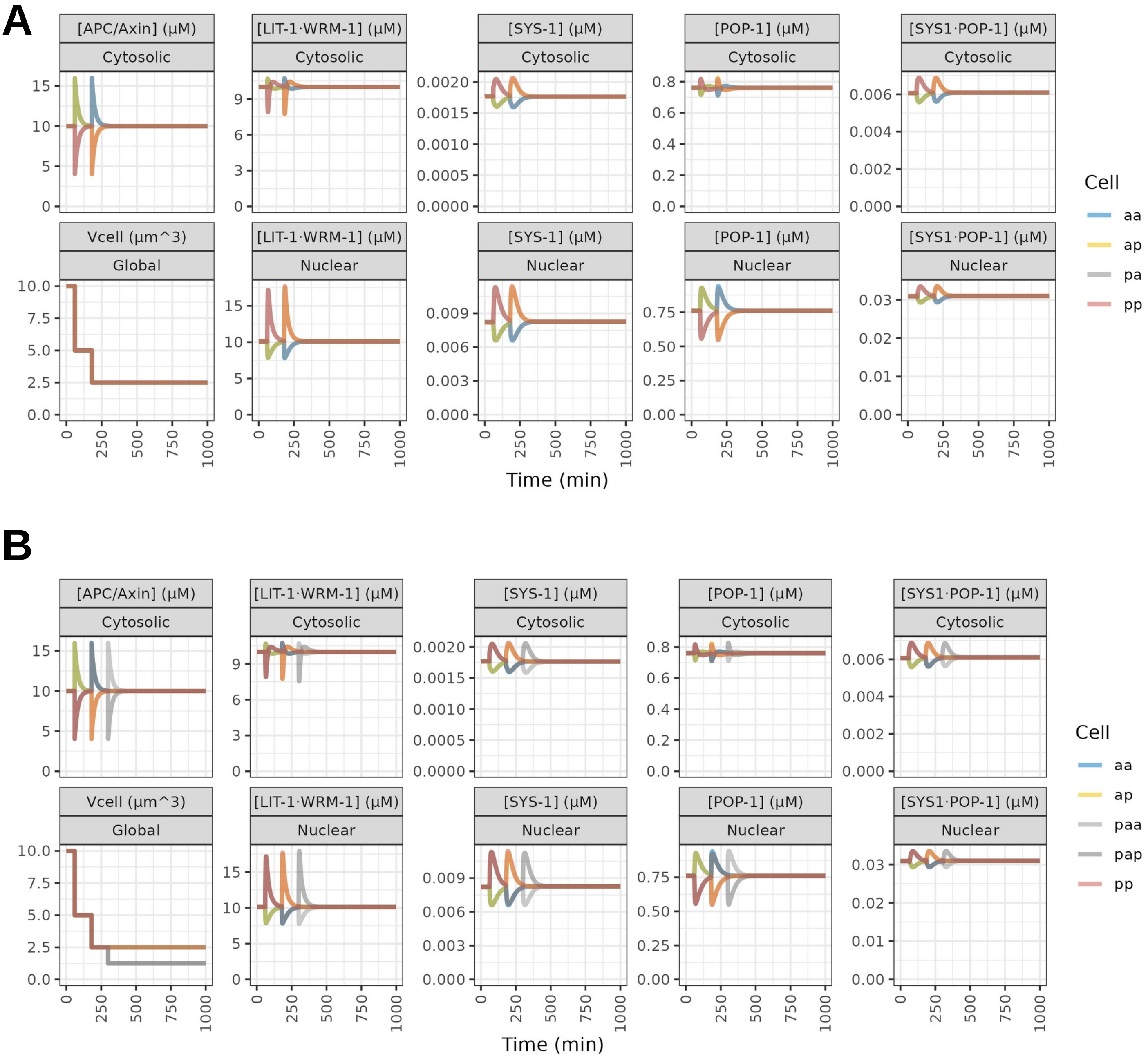


**Figure S5**. Numerical simulation of Model WβA in the non-cumulative regime.

A

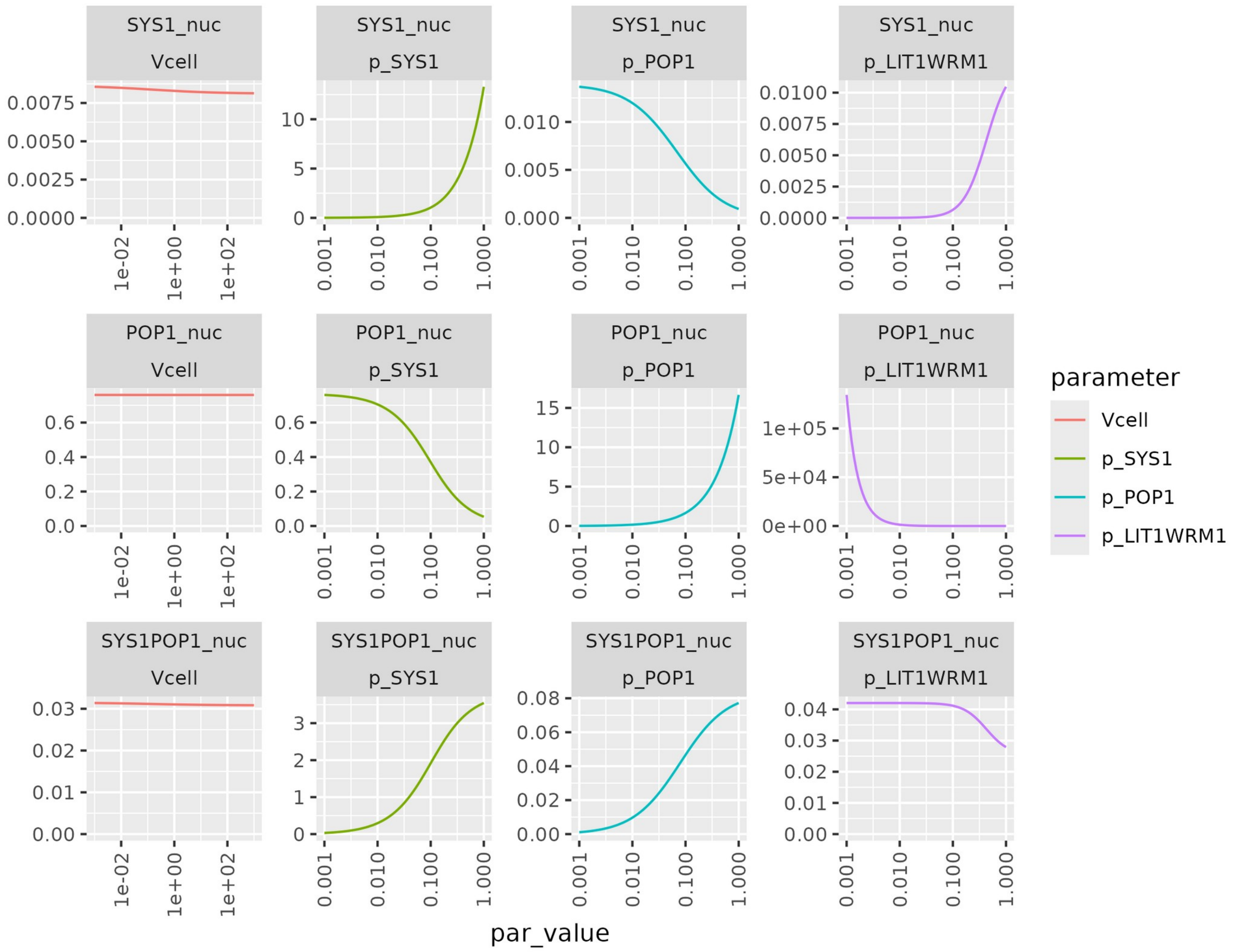


**Figure S6**. Steady state concentrations of nuclear SYS-1, POP-1 and SYS-1·POP-1 as a function of $V_{cell}$, $p_{SYS1}$ and $p_{POP1}$. The influence of $V_{cell}$ is negligible

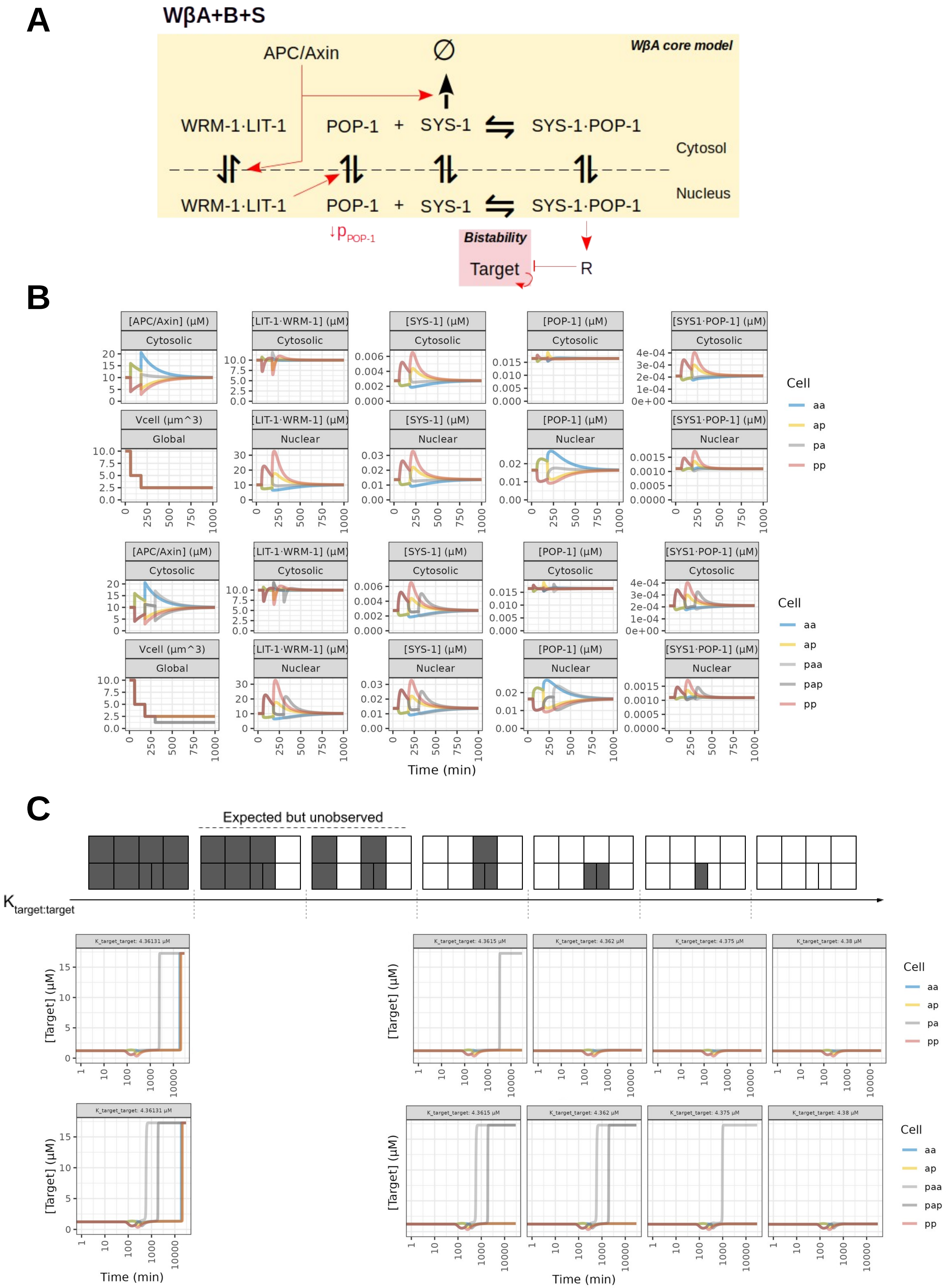


**Figure S7**. WBA+B+S model. (A) Model diagram. (B) Numerical simulations of time dynamics across WβA pathway components. (C) Target dynamics for different values of $K_{target:target}$. Steady-state target concentrations are represented as 2-D projections.

A

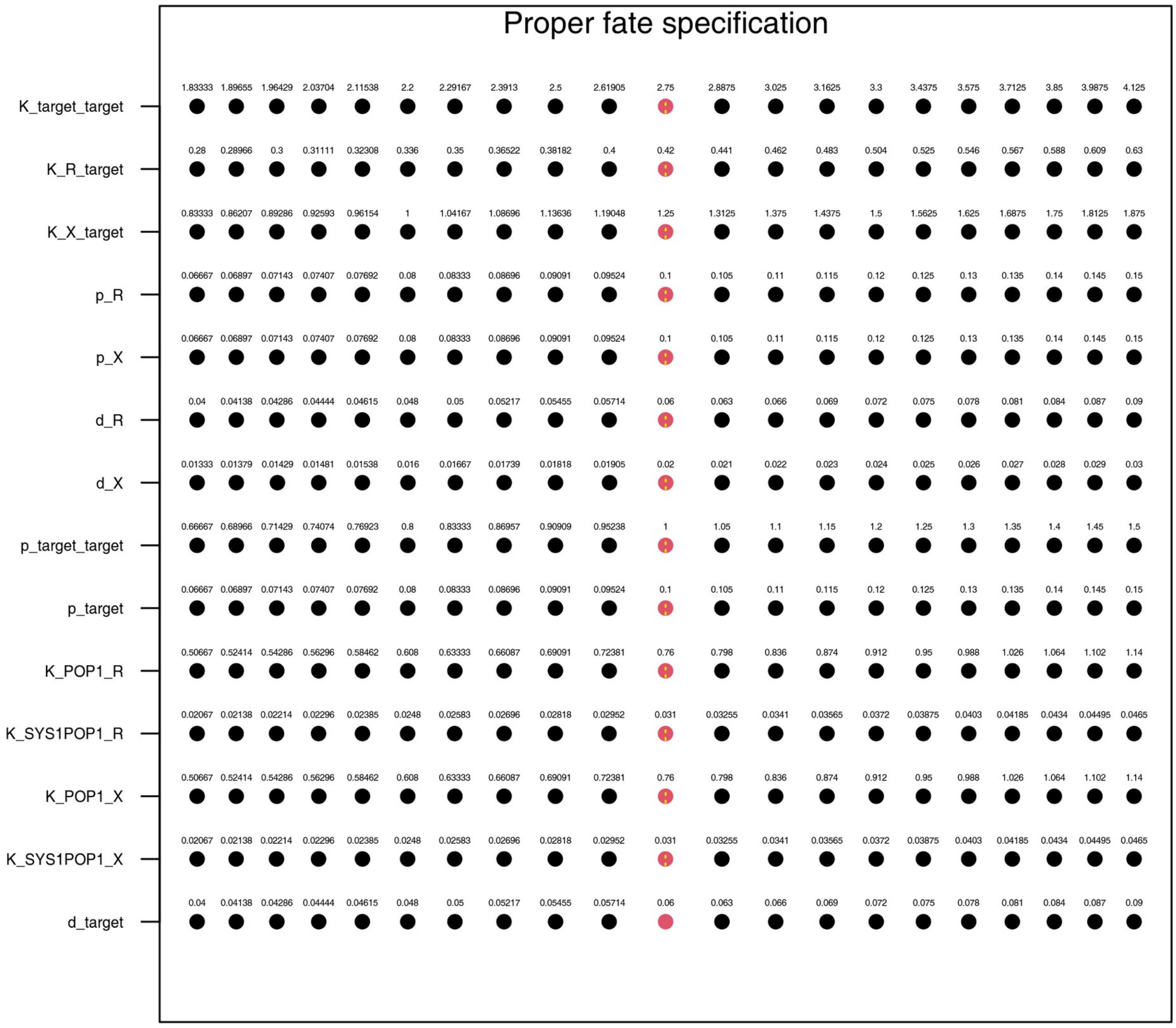


**Figure S8**. Parameter sensitivity analysis of the WBA+B+I model. Wildtype fate specification (red), other (black).

A

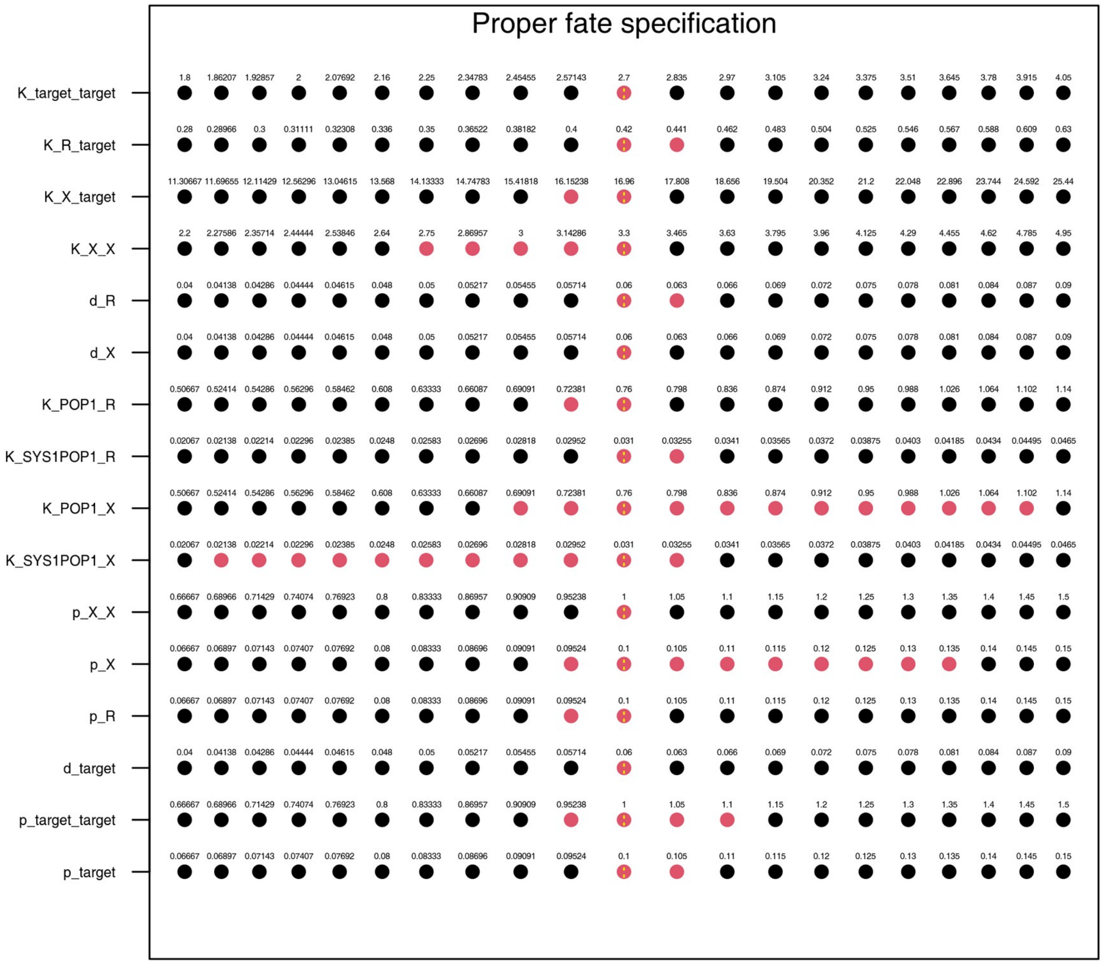


**Figure S9**. Parameter sensitivity analysis of the WBA+B+I+M model. Wildtype fate specification (red), other (black).

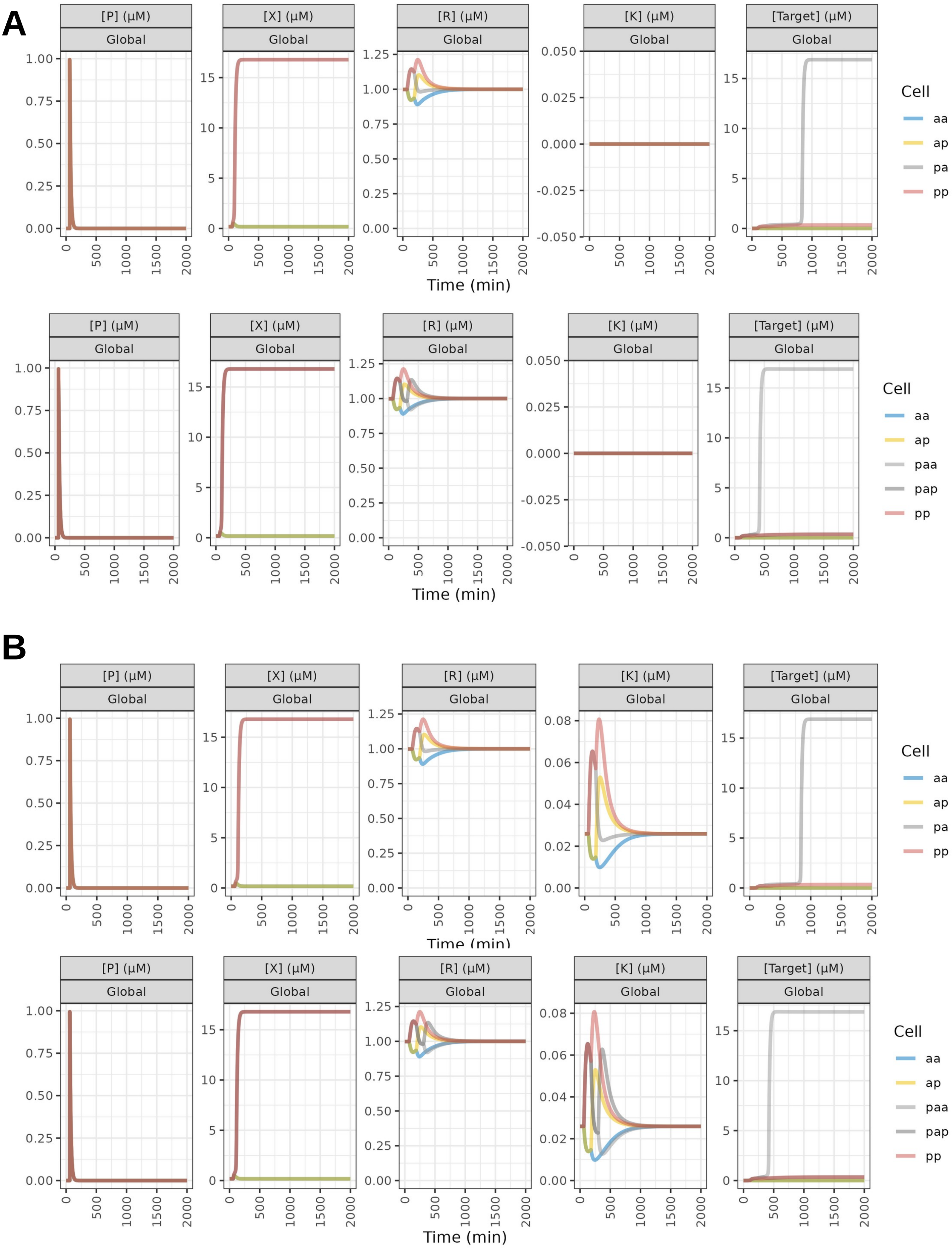


**Figure S10**. Temporal symmetry break (TSB) model time dynamics. Numerical simulations of time dynamics across components using a dorsal and ventral M lineage proliferation schemes (A) without killswitch or (B) with killswitch. Solely [P], [X], [R], [K] and [target] are shown.

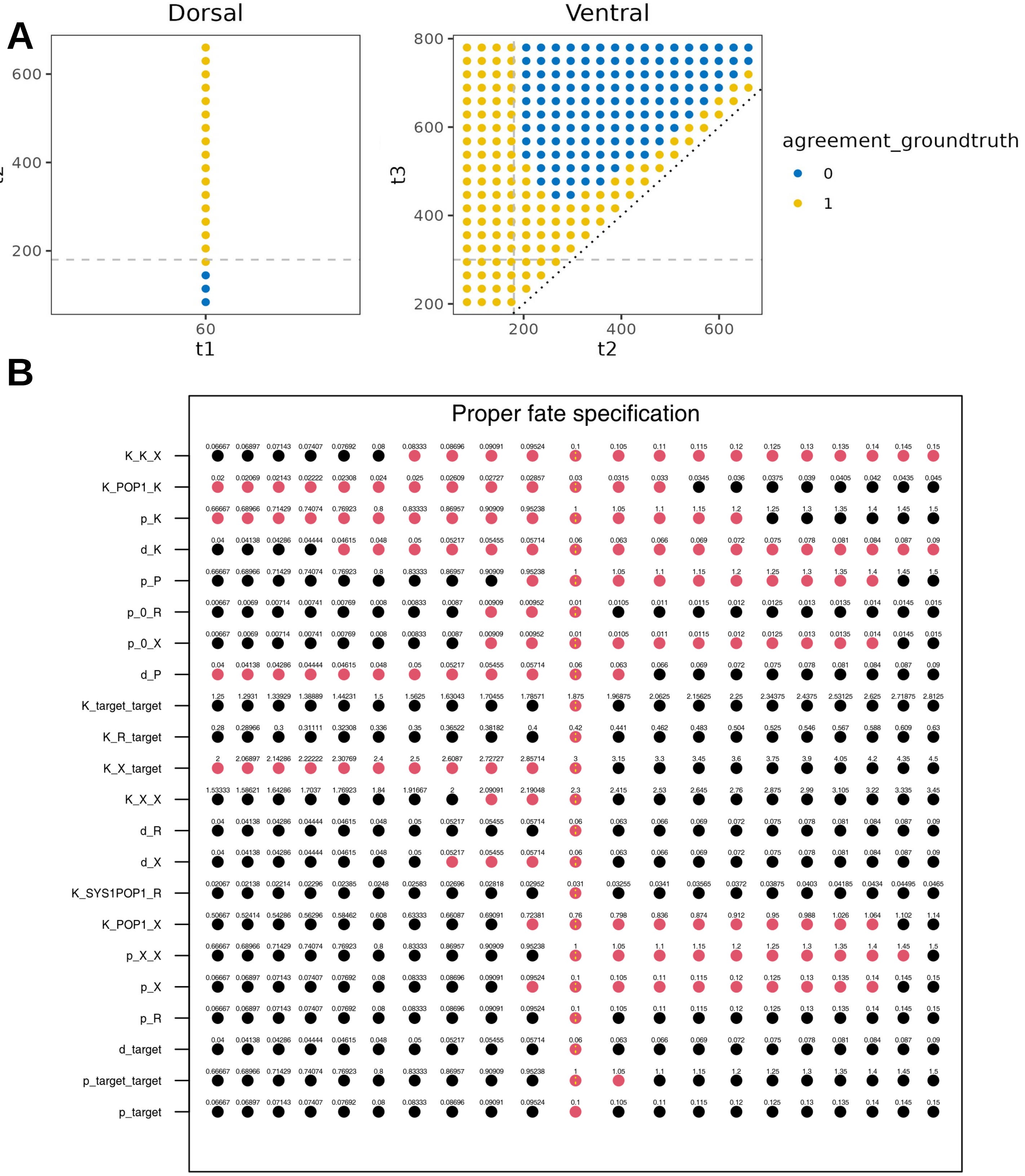


**Figure S11**. Robustness of the temporal symmetry break (TSB) model. B) Numerical simulation in the dorsal and ventral side proliferation schemes and timing robustness, where $\tau_1$ = 60 min; $\tau_2$ = 180 min; $\tau_3$ = 300 min, correspond to the different times of anterior-posterior cell divisions. (B) Parameter sensitivity analysis. Wildtype fate specification (red), other (black).

**A**

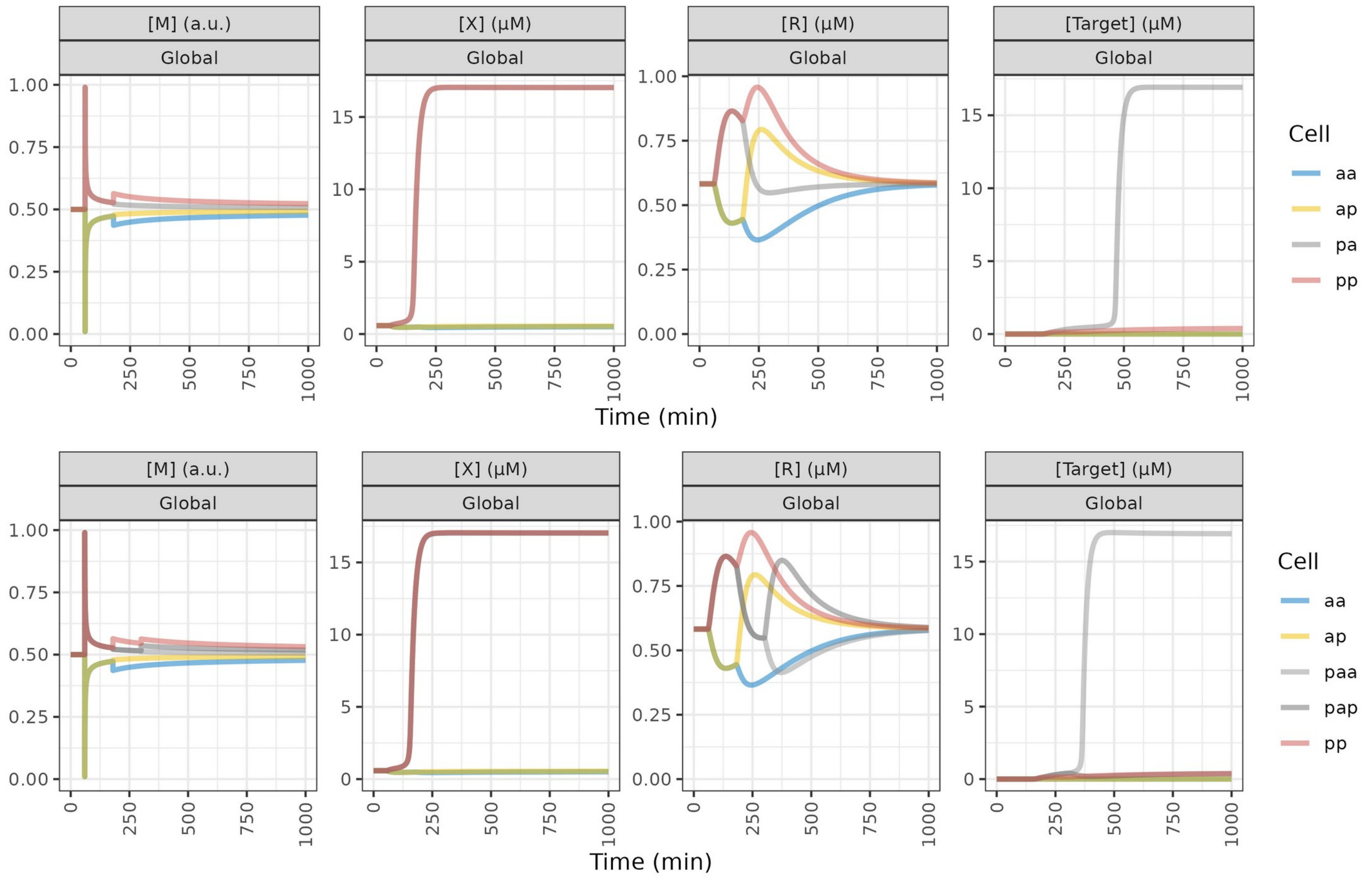


**Figure S12**. Spatial symmetry break (SSB) model time dynamics. Numerical simulations of time dynamics across WβA pathway components using a dorsal and ventral M lineage proliferation schemes. Solely [M], [X], [R] and [target] are shown.

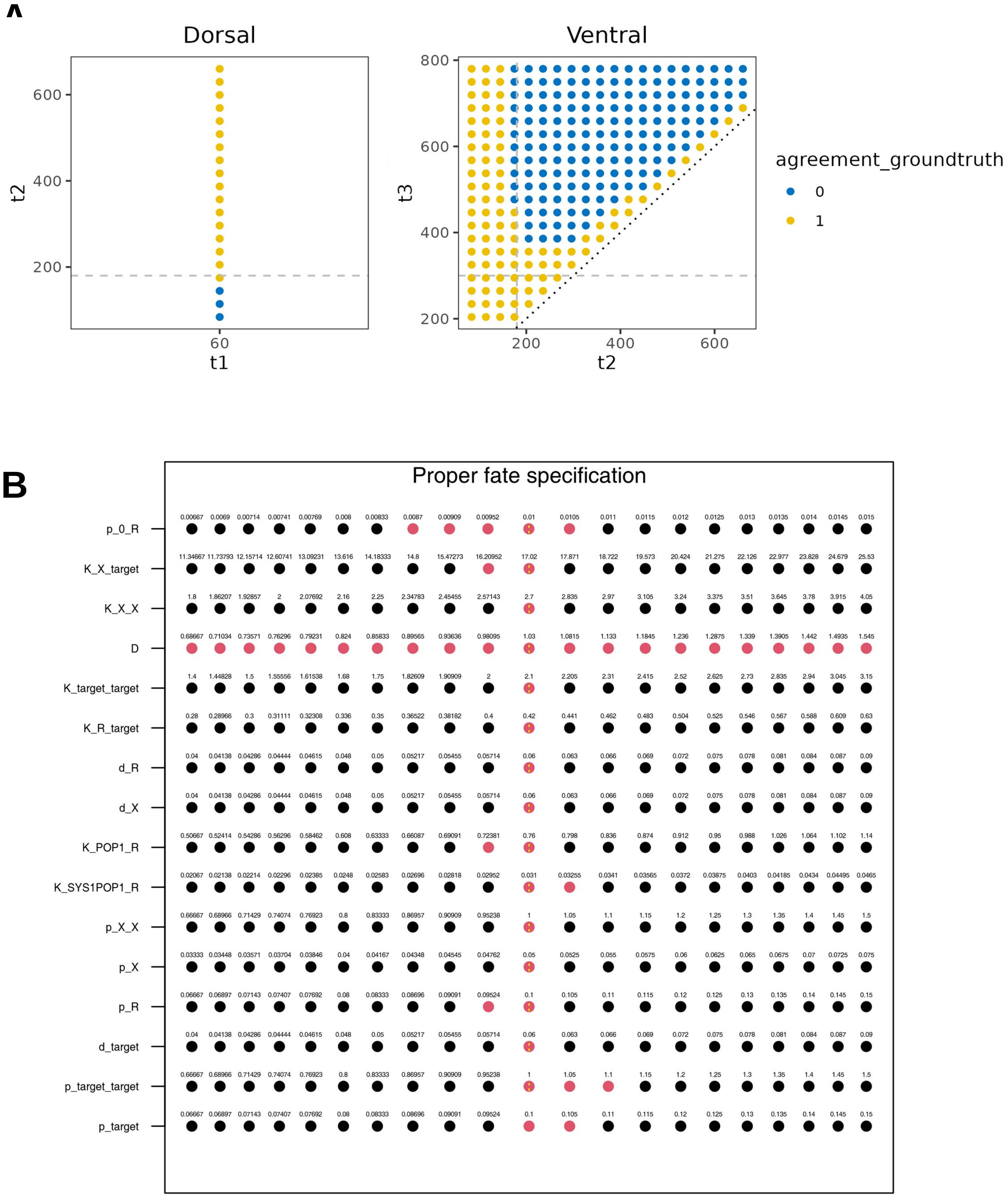


**Figure S13**. Robustness of the spatial symmetry break (SSB) model. B) Numerical simulation in the dorsal and ventral side proliferation schemes and timing robustness, where $\tau_1$ = 60 min; $\tau_2$ = 180 min; $\tau_3$ = 300 min, correspond to the different times of anterior-posterior cell divisions. (B) Parameter sensitivity analysis. Wildtype fate specification (red), other (black).
